# From Calcium-dependent currents to Multitarget Therapy: A Pharmacological Strategy for Glioblastoma

**DOI:** 10.64898/2026.11.16.765581

**Authors:** Cesar Adolfo Sanchez Trivino, Alessia Norbedo, Alice Scomparin, Licio Collavin, Tamara Ius, Ivana Manini, Anna Menini, Daniela Cesselli, Elena Rampazzo, Luca Persano, Loredana Casalis, Pietro Parisse, Miran Skrap, Renza Spelat, Vincent Torre

## Abstract

Glioblastoma (GBM) is an aggressive primary brain tumor with limited treatment options and substantial interpatient heterogeneity. Ion transport and cell-volume regulation may represent pharmacological vulnerabilities in glioma stem cells (GSCs). Here, we characterized membrane currents in patient-derived GSCs, identifying a calcium-dependent anion conductance consistent with volume-regulated anion channels (VRACs) and an inward current consistent with Na⁺/Ca²⁺ exchanger (NCX) activity. Based on these findings, we investigated individual compounds and developed a multitarget cocktail comprising niflumic acid (NFA), bumetanide (BTN), and R-ketorolac (R-K) to target complementary mechanisms involved in ion homeostasis, cell-volume regulation, and cellular dynamics. The cocktail affected cell viability and mechanical properties and showed synergistic effects in U87 cells and patient-derived GSCs with similar electrophysiological profiles. Treatment did not measurably alter spontaneous activity in cultured hippocampal neurons but caused substantial HMC3 cytotoxicity. Encapsulation in transferrin-functionalized liposomes attenuated HMC3 toxicity while preserving activity against glioblastoma cells, suggesting a potential strategy to improve treatment selectivity. Further pharmacological studies identified a potential vulnerability associated with NCX activity, although its contribution to cytotoxicity remains unclear. Overall, our findings highlight electrophysiological heterogeneity in patient-derived GSCs and support further investigation of ion transport as a therapeutic target in GBM. Combining multitarget pharmacology with targeted drug delivery may help address heterogeneous treatment responses, although further studies are needed to establish the underlying mechanisms, safety, and vivo efficacy.

## Introduction

Glioblastoma (GBM) is one of the most aggressive primary brain tumors in adults and remains associated with a poor prognosis despite multimodal treatment approaches combining surgery, radiotherapy, and chemotherapy. Its incidence is 2–5 cases per 100,000 people (1) and median survival remains limited, at approximately one year in many reported cohorts (2). The chemotherapy agents used in the standard treatment of GBM, such as temozolomide (1, 3), are primarily exerting their effects by inducing DNA damage, thereby interfering with cell proliferation (4). However, treatment efficacy is frequently limited by intrinsic or acquired resistance, extensive intratumoral heterogeneity, and the highly infiltrative nature of GBM (5, 6). These challenges underscore the need to investigate alternative therapeutic strategies that target cellular mechanisms beyond conventional DNA-damaging pathways.

Among the mechanisms that may contribute to glioma cell proliferation and invasion, ion transport and cell-volume regulation have received increasing attention (7). Changes in cell volume require coordinated interactions between transmembrane ion fluxes, osmotic water movement, and cytoskeletal remodeling. These processes are particularly relevant during cell division, when cells undergo substantial changes in shape and volume, and during migration, which requires dynamic regulation of cellular mechanics. Recent work has highlighted the contribution of intracellular Ca²⁺ signaling to the final stages of mitosis in glioma cells, where increased frequency and amplitude of cytoplasmic Ca²⁺ transients accompany changes in cell volume (8, 9). Calcium-dependent anion conductances may contribute to these processes by modulating transmembrane chloride fluxes and the associated osmotic movement of water. In addition, transporters involved in ionic homeostasis and intracellular Ca²⁺ regulation may influence cellular functions relevant to tumor progression (10–13). Characterizing these electrophysiological mechanisms in patient-derived glioma stem cells (GSCs) may therefore help identify potential therapeutic vulnerabilities.

In this study, we first characterized the calcium-activated membrane currents in patient-derived GSCs to investigate their electrophysiological properties and identify candidate pharmacological targets. This analysis revealed a calcium-dependent anion conductance with properties consistent with volume-regulated anion channels (VRACs) (14) that was sensitive to NFA (10), as well as an inward current with biophysical and pharmacological features consistent with Na⁺/Ca²⁺ exchanger (NCX) activity (15) that was partially inhibited by doxorubicin (Doxo).

We initially investigated niflumic acid (NFA), an inhibitor of several chloride conductances, including VRAC-like currents on U87 cells, as a potential pharmacological approach to impair glioblastoma cell viability However, because cell-volume regulation depends on the coordinated activity of membrane ion transporters, chloride conductances, and cytoskeleton-associated signaling pathways, we hypothesized that targeting complementary mechanisms could enhance the therapeutic effects. This rationale led us to explore a multitarget strategy combining FDA approved drugs: NFA with bumetanide (BTN), an inhibitor of Na⁺-K⁺-2Cl⁻ and K⁺-Cl cotransporters, particularly NKCC1 (16, 17), and R-ketorolac (R-K), which has been associated with the modulation of Rac1-and Cdc42-related signaling (18). By targeting complementary processes involved in ion homeostasis, cell-volume regulation, and cytoskeletal dynamics (19) this combination was designed to investigate whether a broader pharmacological intervention could improve the effects of individual compounds on glioblastoma cells.

We then extended our investigation to patient-derived GSCs cultures to assess the efficacy of the cocktail and determine whether treatment sensitivity was associated with electrophysiological profile. Finding that treatment responses showed a high variability across patient-derived cultures. We also explored NCX-associated pharmacological vulnerability (15) in a separate patient-derived culture as a potential alternative therapeutic target.

Finally, to address potential off-target effects of the multitarget cocktail, we investigated its encapsulation in transferrin-functionalized liposomes (20). Transferrin functionalization was explored as a strategy to improve the specificity of drug delivery toward glioma cells (21) while potentially limiting exposure of non-malignant cells 20. Overall, this study integrates electrophysiological characterization and pharmacological evaluation to investigate therapeutic vulnerabilities in GBM, assess a multitarget strategy directed at complementary cellular mechanisms, and explore targeted drug delivery as a complementary approach to improve treatment selectivity.

## Results

### SCs from some patients have only Ca-activated approximately linear currents or inward currents

To test the functional expression of calcium-dependent currents in GSCs, we performed electrical measurements using patch clamp in the whole cell configuration, in which the pipette contained CsCl instead of KCl to block potassium currents and the calcium concentration inside the patch pipette was 0.5 μM or zero nominally calcium by buffering with EGTA.

Under these experimental conditions we verified that under calcium-free conditions no appreciable current was measured (134.6 ± 3.82 pA at +100 mV). However, GSCs from some patients (S557) the elevation of intracellular calcium up to 0.5 μM activated large currents characterized by a slow kinetic of inactivation at very positive potentials (4378 ± 722.8 pA at +100 mV, Fig.1A-B). In some GSCs the activated current was outwardly rectified or approximately linear (violet trace, Fig.1A) but the degree of linearity varied among the recorded GSCs (Fig.1B). On the other hand, GSCs from other patients (S568) under identical experimental conditions exhibited only a larger inward current (-508 ± 89.1 pA at -100 mV) compared to cells pathed with a calcium-free internal solution (-115.4 ± 49.8 pA at -100 mV). However, GSCs from most patients had a mixture of both the outward and inward currents. In some GSCs we observed a time dependent increase of the outward current which was initially inward but in 4 minutes transformed into an approximately linear current (Fig.1E). This time dependent changes are ascribed to the Ca loading into the impaled GSC. In other experiments, when extracellular Na was replaced by NMDG the approximately linear current became an outward rectifying current (Fig.1F) suggesting that in most of the GSCs there is a concomitant presence of a VRAC-like Cl current (see Fig.2) and of a Na current associated to the activation of the Na-Ca exchanger (see Fig.3) which “linearize” the Ca activated current.

**Figure 1:**
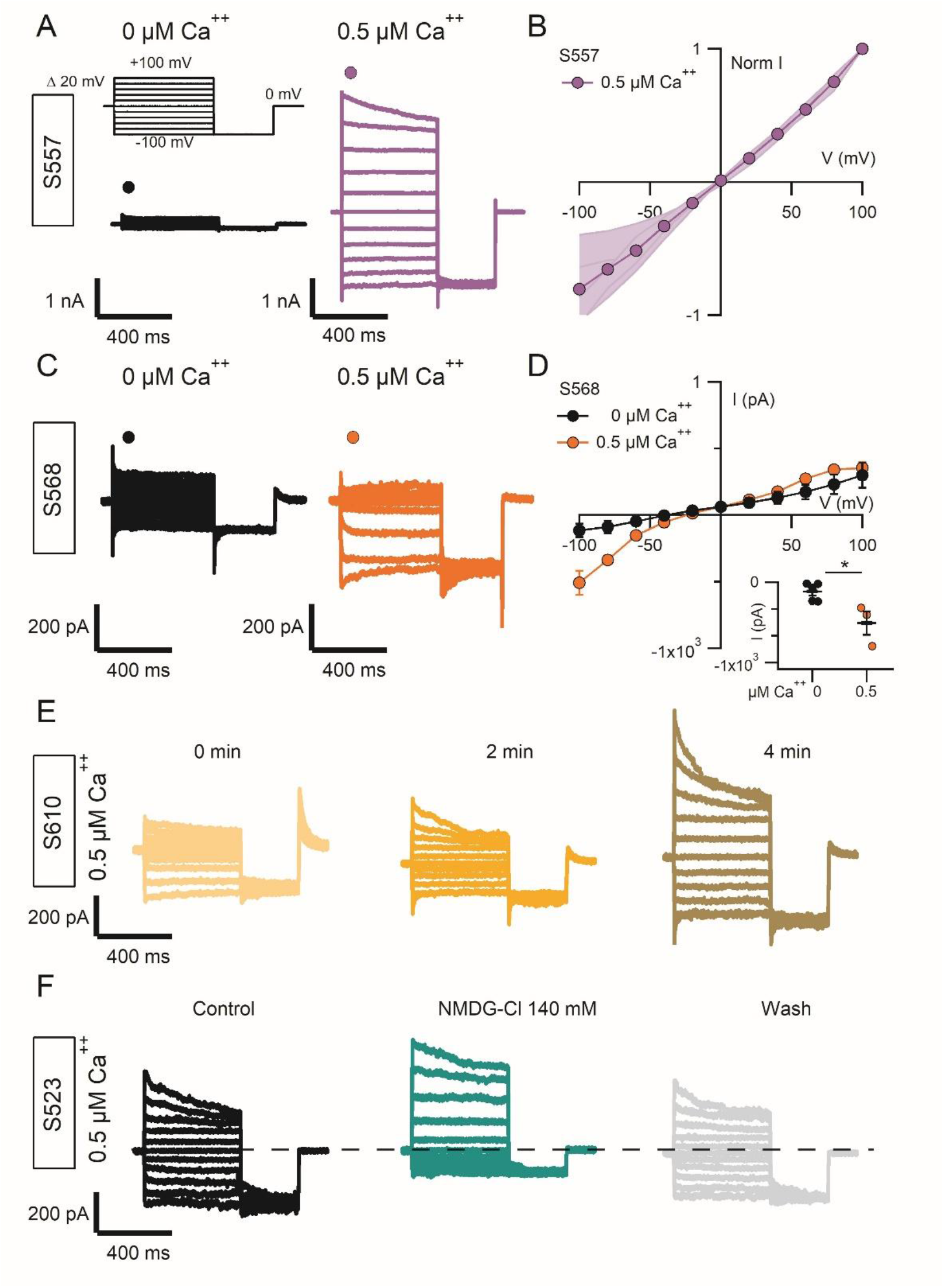
Whole-cell electrophysiological recordings from patient-derived GSCs. **(A)** Representative whole-cell current traces from S557 GSCs recorded with 0 μM (black) or 0.5 μM (violet) intracellular Ca²⁺ during hyperpolarizing and depolarizing voltage steps. **(B)** Normalized current–voltage (I–V) relationships from five S557 GSCs recorded with 0.5 μM intracellular Ca²⁺. **(C)** Representative recordings from S568 GSCs under the same conditions, showing a Ca²⁺-dependent inward current during hyperpolarization (orange). **(D)** Corresponding I–V relationships. Inset: peak inward current at −100 mV. **(E)** Time course of whole-cell currents in an S610 GSC, showing an early inward current followed by the development of a linear current component over 2–4 min after break-in. **(F)** Representative recordings from an S523 GSC before (black) and after extracellular Na⁺ replacement with 140 mM NMDG-Cl (green), showing suppression of the inward current and enhancement of the outward component

**Figure 2.**
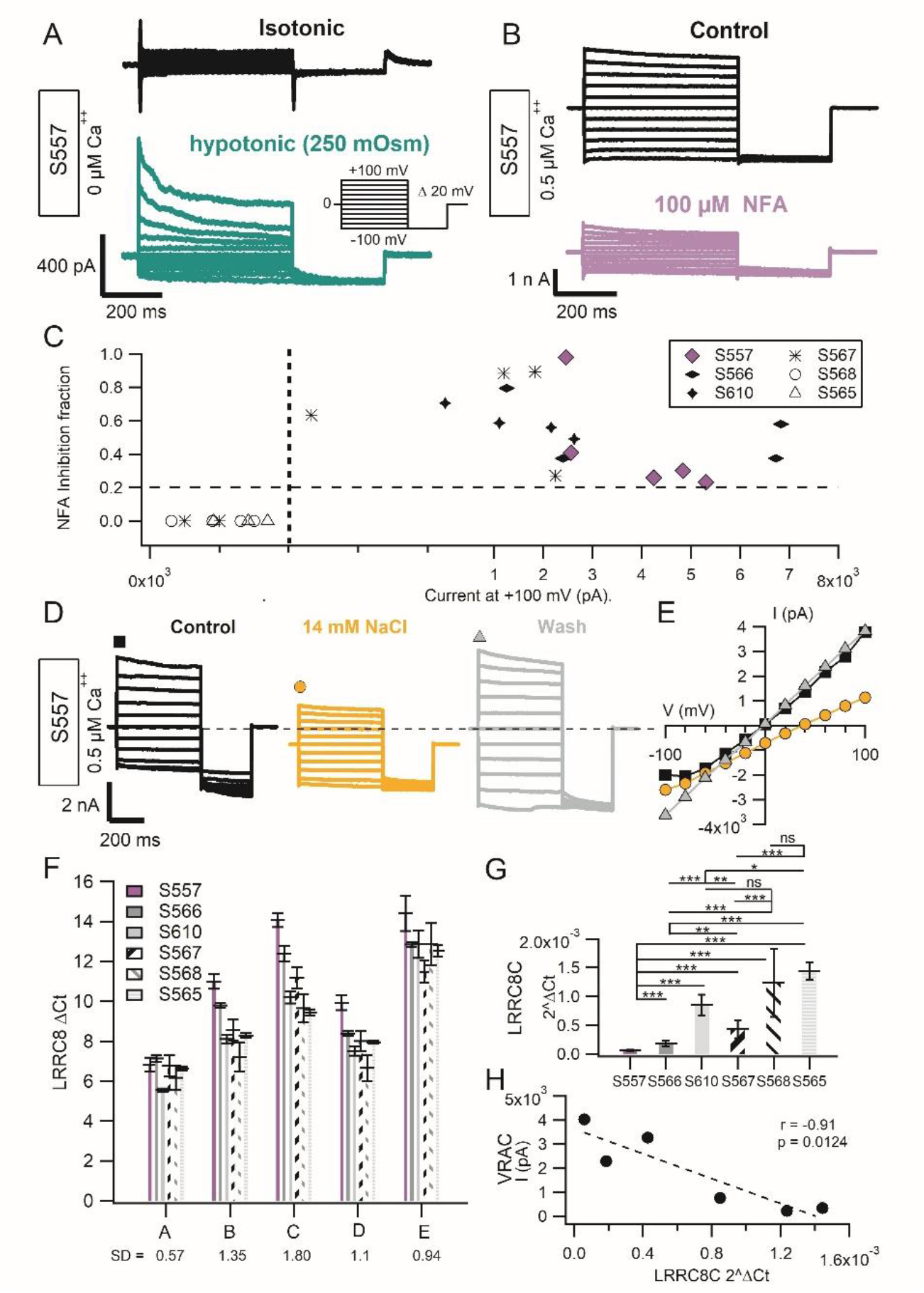
Identification of heterogeneous VRAC-like chloride currents in patient-derived GSCs. **(A)** Representative whole-cell recordings from an S557 GSC under isotonic (black) and hypotonic (250 mOsm; green) conditions, showing activation of an outwardly rectifying current with slow deactivation at positive potentials. Intracellular Ca²⁺ was nominally zero. Inset: voltage-step protocol (−100 to +100 mV). **(B)** Representative recordings from an S557 GSC with 0.5 μM intracellular Ca²⁺, showing inhibition by niflumic acid (NFA, 100 μM). **(C)** Relationship between NFA inhibition and outward current amplitude at +100 mV across patient-derived GSCs. Each symbol represents one cell. Dashed lines indicate empirical thresholds for current amplitude and inhibition. **(D)** Representative recordings from an S557 GSC under control conditions (black), following extracellular NaCl reduction to 14 mM (orange), and after washout (gray). **(E)** Corresponding current–voltage (I–V) relationships, showing reduced outward current and a positive shift in reversal potential under low-NaCl conditions. **(F)** Variability in LRRC8 subunit expression across patient-derived GSC cultures, with LRRC8C showing the greatest variability. **(G)** LRRC8C expression across patient-derived GSC cultures. **(H)** Correlation between maximal outward current amplitude at +100 mV and LRRC8C expression.

**Figure 3.**
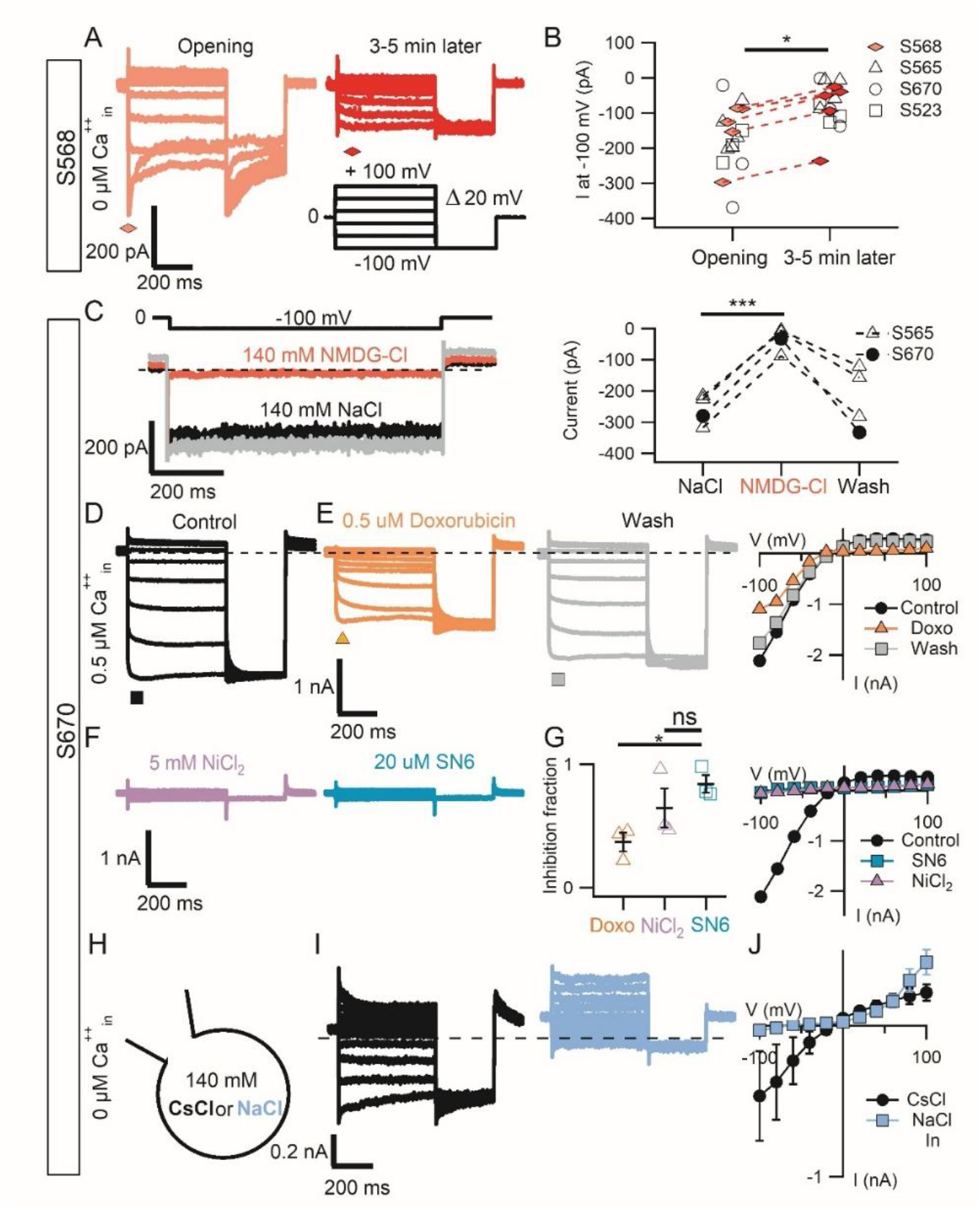
Identification Electrophysiological and pharmacological characterization of a Na^+^/Ca^2+^ exchanger-like current in patient-derived glioblastoma stem cells (GSCs). **(A)** Representative whole-cell recordings from GSCs under nominally Ca²⁺-free intracellular conditions, showing an inward current that progressively decreased after membrane break-in. **(B)** Quantification of inward current at −100 mV immediately after break-in and 3–5 min later in GSCs from patients S568 (n = 5), S565 (n = 5), S670 (n = 3), and S523 (n = 3) (p = 0.043114, Wilcoxon test). **(C)** Representative recordings with 0.5 μM intracellular Ca²⁺ showing suppression of the inward current following extracellular Na⁺ replacement with NMDG-Cl (left). Quantification under control, NMDG-Cl, and washout conditions (n = 4; p = 0.0008112, paired *t*-test; right). **(D)** Representative inward current recorded with 10 mM intracellular Na⁺, 0.5 μM intracellular Ca²⁺, and 140 mM extracellular Na⁺ and 2 mM Ca²⁺. **(E)** Partial and reversible inhibition of the inward current by 0.5 μM doxorubicin (left and middle), with the corresponding current–voltage (I–V) relationship (right). **(F)** Representative recordings showing inhibition of the inward current by 5 mM NiCl₂ and 20 μM SN-6. **(G)** Quantification of current inhibition by doxorubicin, NiCl₂, and SN-6 (p = 0.01707, Kruskal–Wallis test followed by Dunn’s post hoc test with Bonferroni correction; left). Corresponding I–V relationships showing suppression of the inward current by SN-6 and NiCl₂ (right). **(H)** Schematic representation of recording conditions with nominally Ca²⁺-free intracellular solution containing either CsCl (black) or 140 mM NaCl (blue). **(I)** Representative recordings under the conditions described in (H), showing loss of the inward current with intracellular NaCl (blue). **(J)** Corresponding I–V relationships, showing suppression of the inward current and predominance of the outward component. Inward current at −100 mV: p = 0.0357, Mann–Whitney test. Data are presented as mean ± SEM. Statistical significance was assessed using paired or unpaired tests, as appropriate. *p < 0.05; ***p < 0.001.

In summary, in GSCs extracted from patients with high grade brain tumors, the elevation of intracellular calcium drives two kinds of currents: an inward or an outward current and more often both.

### The VRAC-like outward current is carried by Cl ions and is blocked by NFA

In the absence of added Ca²⁺ in the patch pipette and when intracellular K⁺ was replaced by Cs⁺ - no appreciable ionic currents were typically observed in GSCs (>95% of recordings). However, when the extracellular solution was replaced with a hypo-osmotic solution, a classical stimulus used to activate the VRAC channel (11), a clear outwardly rectifying current was induced (Fig. 2A). his current displayed the characteristic slow and nearly linear deactivation at positive voltages that is typical of VRAC-mediated currents (22, 23).

To determine whether the outward current component activated by intracellular Ca²⁺ in GSCs shared electrophysiological properties with VRAC currents, we analyzed the deactivation kinetics by fitting the currents activated at +100 mV with a single exponential function. Recordings obtained from 67 cells derived from 11 patients revealed marked heterogeneity in the deactivation time constants, with cells segregating into two clearly distinct clusters (Supplementary Fig. 1). One group, characterized by larger linear outward currents, exhibited relatively fast deactivation kinetics (e.g., S566: 107.5 ± 32.4 ms, n = 9), whereas a second group displaying smaller currents (<1 nA) showed markedly slower deactivation time constants (e.g., S568: 3929 ± 389.4 ms, n = 8). These observations are consistent with the known variability in VRAC biophysical properties arising from differences in channel subunit composition (24).

Another hallmark of VRAC channels is their sensitivity to niflumic acid (NFA). We therefore evaluated the inhibitory effect of NFA on the outward current measured at +100 mV. In GSCs derived from patient S557, the outward current was strongly blocked by NFA (Fig. 2B). However, substantial variability in the degree of inhibition was observed both among cells from the same patient and across different patients (Fig. 2C). Again, two major populations could be distinguished: cells exhibiting large outward currents showed inhibition ranging from 20% to nearly complete blockade belonging to patients S557, S566 and S610, whereas cells from patients 565 and 568 showed smaller currents displayed little or no sensitivity to NFA.

To further assess whether these currents were predominantly carried by anions, as expected for VRAC channels, extracellular NaCl concentration was markedly reduced (14 mM NaCl; Fig. 2D). Under these conditions, the amplitude of the outward current decreased and the reversal potential shifted toward more positive values (ΔV = +37.94 ± 2.93 mV on 5 cells; Fig. 2E), consistent with a chloride-dependent conductance.

Taken together, these results support the presence of VRAC-like currents in a subset of GSCs, although with substantial electrophysiological heterogeneity. Such functional variability may reflect the differential expression of distinct VRAC channels subunits belonging to the LRRC8 family proteins, which can determine the biophysical properties and sensitivity to NFA To assess whether the expression of VRAC subunits reflects the observed functional variability, we performed qRT-PCR analysis of the LRRC8 subunits. Expression levels differed significantly among LRRC8 subunits, consistent with previous studies in GBM (13) (Supplementary Fig. 2, Friedman test, p < 0.001). Among the VRAC subunits, LRRC8C exhibited the greatest inter-individual variability in expression across the six patient-derived samples analyzed, based on the standard deviation (SD) calculated for each subunit (Fig. 2F; LRRC8C, SD = 1.8). We therefore focused our subsequent analysis on patient-to-patient variability in LRRC8C expression. Notably, LRRC8C expression provided greater separation among patients, with GSCs derived from patients S565 and S568 showing the highest expression levels, whereas those derived from patient S557 exhibited markedly lower expression (Fig. 2G). We next performed a linear correlation analysis between the mean VRAC-like current at +100mV in GSCs derived from each patient and their relative LRRC8C expression. This analysis revealed a strong negative correlation between LRRC8C expression and VRAC-like activated current (Fig. 2H), supporting the hypothesis that GSCs exhibit distinct patterns of LRRC8 subunit expression that may contribute to the biophysical and pharmacological differences observed in the electrophysiological experiments.

### The inward current is carried by the Na^+^ entry concomitant to the activity of the Na^+^-Ca^2+^ exchanger

In several experiments using a nominally Ca^2+^-free internal solution, we observed an inward current of some hundred pA, mainly in patients S658, S565, S523 and S670 (Fig. 3A), with a mean amplitude of −150.13 ± 38.82 pA at −100 mV (S568; Fig. 3B). Importantly, this inward current progressively decreased within 2–4 min after membrane break-in (−89.26 ± 38.55 pA at −100 mV; Fig. 3B), as expected for a current strongly dependent on intracellular Ca^2+^, which gradually equilibrates with the buffered pipette solution.

In subsequent experiments performed with 0.5 μM free Ca²⁺ in the patch pipette, cells in which the VRAC-like linear current did not develop, the inward current (−320.97 ± 91.68 pA) remained stable over time. To assess the ionic permeability underlying this conductance, extracellular Na⁺ was replaced with NMDG (initial current: −259.9 ± 23.79 pA; Fig. 3C, black trace). Under these conditions, the inward current was markedly reduced to −34.6 ± 18.7 pA (Fig. 3C, Red trace), indicating that the current is primarily carried by cations.

A current with similar characteristics was recorded in a cellular model of glioblastoma U251, which was associated to the Na⁺/Ca²⁺ exchanger (25). Under our recording conditions, the pipette solution contained 10 mM Na⁺ and 0.5 μM free Ca²⁺, whereas the extracellular solution contained 140 mM Na⁺ and 2 mM Ca²⁺, corresponding to a theoretical Na⁺/Ca²⁺ exchanger reversal potential of approximately −10 mV, consistent with the current–voltage relationship shown in (Fig. 3E–J). Together, these observations suggest that the conductance may reflect Na⁺/Ca²⁺ exchanger activity operating in the forward mode (26, 27).

To validate the expression of the exchanger, we performed immunofluorescence against the Na^+^/ Ca^2+^ exchanger in GSCs derived from patient S670 (Supplementary Fig. 3). Functional validation was then performed in GSCs from the same patient using pharmacological inhibitors of the exchanger. Among these, we tested doxorubicin, an FDA-approved chemotherapeutic agent previously used in breast and bladder cancer treatment (28, 29) that has also been reported to inhibit the Na^+^/Ca^2+^ exchanger (30, 31). Application of 0.5 μM doxorubicin produced a reversible partial inhibition of the inward current (Fig. 3D–E), together with a complete suppression of the small outward component (Fig. 3E), suggesting that doxorubicin affects both the forward and reverse operating modes of the exchanger.

As additional pharmacological characterization, we tested the classical Na^+^/Ca^2+^ exchanger inhibitors NiCl_2_ (32) and SN6 (33). NiCl_2_ produced a 64.5 ± 15.9% inhibition, whereas the more selective inhibitor SN-6 reduced the current by 83.9 ± 7% (Fig. 3F–G). Representative traces show that both Ni^2+^ and SN-6 nearly abolished the inward current (Fig. 3F).

To further characterize the Na^+^/Ca^2+^ exchanger-like inward current, we replaced intracellular CsCl with 140 mM NaCl while maintaining nominally 0 μM Ca^2+^ in the pipette solution (Fig. 3H), thereby collapsing the Na^+^ gradient and shifting the exchanger reversal potential. Assuming a free intracellular Ca^+^ concentration of ∼200 nM, estimated from the EGTA-buffered internal solution, the theoretical reversal potential under these conditions was calculated to be approximately −246 mV. As expected from this large negative shift in reversal potential, the inward current disappeared and was replaced by a net outward current at 0 mV together with an incipient outward rectification favored by the strong electrochemical driving force (blue traces, Fig. 3I–J).

### GSCs from the great majority of patients have a mixture of both currents

To broadly assess the presence of calcium-triggered currents, the mean currents recorded from the 10 patients were compared. First, the mean amplitude of the VRAC-like current was calculated at the beginning of the +100 mV voltage step in cells dialyzed with 0.5 µM Ca²⁺ in the pipette solution and displaying the characteristic inactivating kinetics (Fig. 1A). To identify the presence of the inward current carried by the Na–Ca exchanger was more difficult, due to the fact that in cells with both current, the linearity of VRAC contaminate the current at negative potentials, to avoid that, the inward current at −100 mV was measured and verified its disappearance when GSCs were patched with electrodes containing free Ca²⁺; the mean amplitude of this initial current was then calculated (Fig. 3A).

Our analysis showed that GSCs from most patients exhibited both currents. To evaluate their relative contribution, we calculated the ratio between the VRAC-like current and the Na–Ca exchanger current. Ratios greater than 5 were observed in patients S557, S610, S566, S567, S541, and S523, indicating that the outward component was several-fold larger than the inward current. In contrast, patients S562, S670, S565, and S568 showed ratios close to 1, suggesting a comparable contribution of both currents (Supplementary Fig. 4).

Based on these findings, we conclude that GSCs from most patients display a mixed pattern of Ca²⁺-activated Cl⁻ currents and inward Na–Ca exchanger currents.

### Heterogeneity of ionic currents in GSCs of some patients

The electrical properties of GSCs were generally reproducible among cells derived from the same patient. However, in some cases we observed a marked heterogeneity within the same culture. A representative example is provided by GSCs from patient S567, which displayed substantial variability in the degree of linearity versus rectification of their membrane currents (Fig. 4A). The current amplitudes measured at +100 mV and −60 mV were broadly distributed across cells (Fig. 4B), and the outward currents recorded at +100 mV exhibited distinct inactivation kinetics (Fig. 4C).

**Figure 4.**
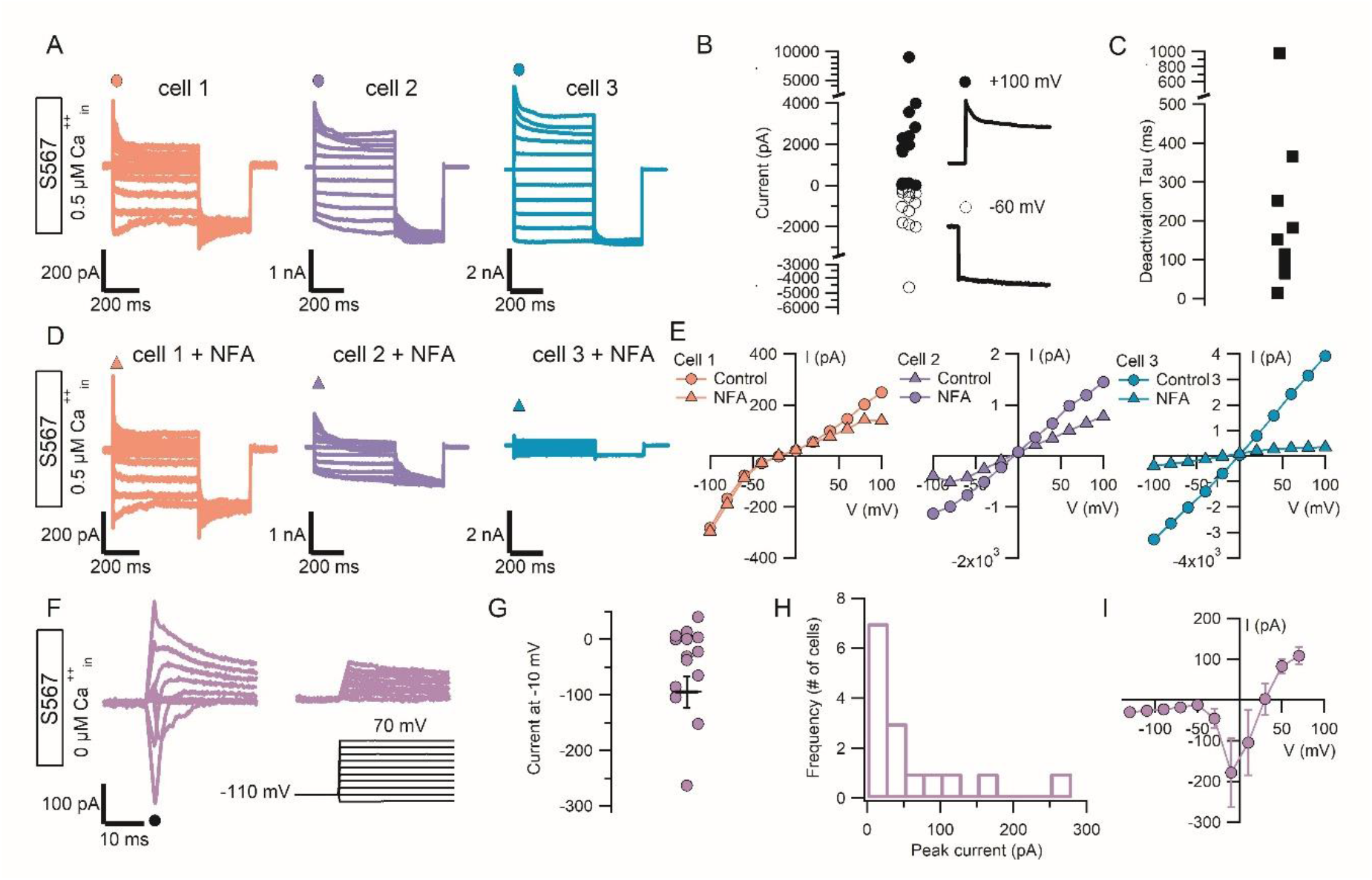
Functional heterogeneity of GSCs from patient S567. **(A)** Representative current traces from three S567 GSCs showing distinct current profiles: predominantly inward (Cell 1), mixed (Cell 2), and nearly linear (Cell 3). **(B)** Distribution of current amplitudes at +100 and −60 mV across individual S567 GSCs, illustrating intercellular variability. **(C)** Representative outward currents at +100 mV showing variable inactivation kinetics. **(D)** Current–voltage (I–V) relationships from the cells in (A) before (open circles) and after NFA application (filled triangles). **(E)** Differential NFA effects: minimal inhibition in Cell 1, preferential suppression of the outward component in Cell 2, and marked current inhibition in Cell 3. The residual inward current in Cell 2 is consistent with a distinct current component, potentially associated with NCX activity. **(F)** Representative voltage-clamp recordings showing transient inward Na⁺ currents in a subset of S567 GSCs. **(G–H)** Distribution of peak transient inward current amplitudes across individual cells. **(I)** I–V relationship of the transient inward Na⁺ current, showing activation at approximately −30 to −40 mV. Data are presented as mean ± SEM where indicated.

This variability was also reflected in the sensitivity to NFA blockade. In cells displaying predominantly inward currents with little or no detectable outward component (Cell 1, Fig. 4A–D), NFA produced little or no significant inhibition (Fig. 4E). In contrast, cells exhibiting an almost linear I–V relationship (Cell 3, Fig. 4A–D) showed a strong suppression of the current by NFA. We also identified cells with intermediate electrophysiological properties (Cell 2, Fig. 4A–D), in which NFA partially inhibited the outward current while leaving a residual inwardly rectifying component. This observation suggests the coexistence of both VRAC-like currents and inward Na–Ca exchanger currents within the same cell population (Fig. 4E).

To determine whether this heterogeneity was restricted to Ca²⁺-triggered currents, we also investigated voltage-dependent inward currents using a holding potential of −110 mV (Fig. 4F). Under these conditions, brief transient inward currents were observed, with a broad distribution of peak amplitudes among cells (Fig. 4G–H). These currents displayed activation kinetics and voltage dependence characteristic of high-threshold voltage-gated Na⁺ currents, consistent with NaV1.2 or NaV1.3 channels (34, 35), with threshold activation occurring around −30 to −40 mV (Fig. 4I). Notably, these transient inward currents were not detected in all GSCs from patient S567, but only in 50% of the recorded cells (Fig. 4F), further highlighting the pronounced functional heterogeneity within this patient-derived GSC population.

### Multi-target strategy improved the cytotoxic effect induced by NFA

A pharmacological strategy targeting cellular volume regulation and cytoskeletal dynamics in GSCs was investigated, starting with the cellular response to NFA, performing cell viability assays in U87 glioblastoma cells, which functionally express a VRAC-like current (Supplementary Fig. 5) and have been reported to respond to NFA (8). First, the optimal treatment window was established by evaluating cell viability after 24, 48, and 72 h of exposure (Supplementary Fig. 6). Subsequently the concentration-dependent effects of NFA were evaluated. After 72 h of treatment, a concentration of 200 µM was required to significantly reduce cell viability compared with the control condition, whereas treatment with 800 µM reduced cell viability to 17.6 ± 2.3% (Fig. 5A). To further investigate the effects of NFA on cellular mechanical properties, Young’s modulus of treated cells was measured by Atomic Force Microscopy (AFM). NFA treatment increased the Young’s modulus by 0.4 ± 0.05 kPa (Fig. 5B), indicating increased cellular stiffness. Which is consistent with reduced cellular deformability, a mechanical state that has been associated with reduced migratory and invasive capacity (36). Together with the observed reduction in cell viability, these findings suggest that NFA may affect both cellular survival and the mechanical properties of glioma cells.

**Figure 5.**
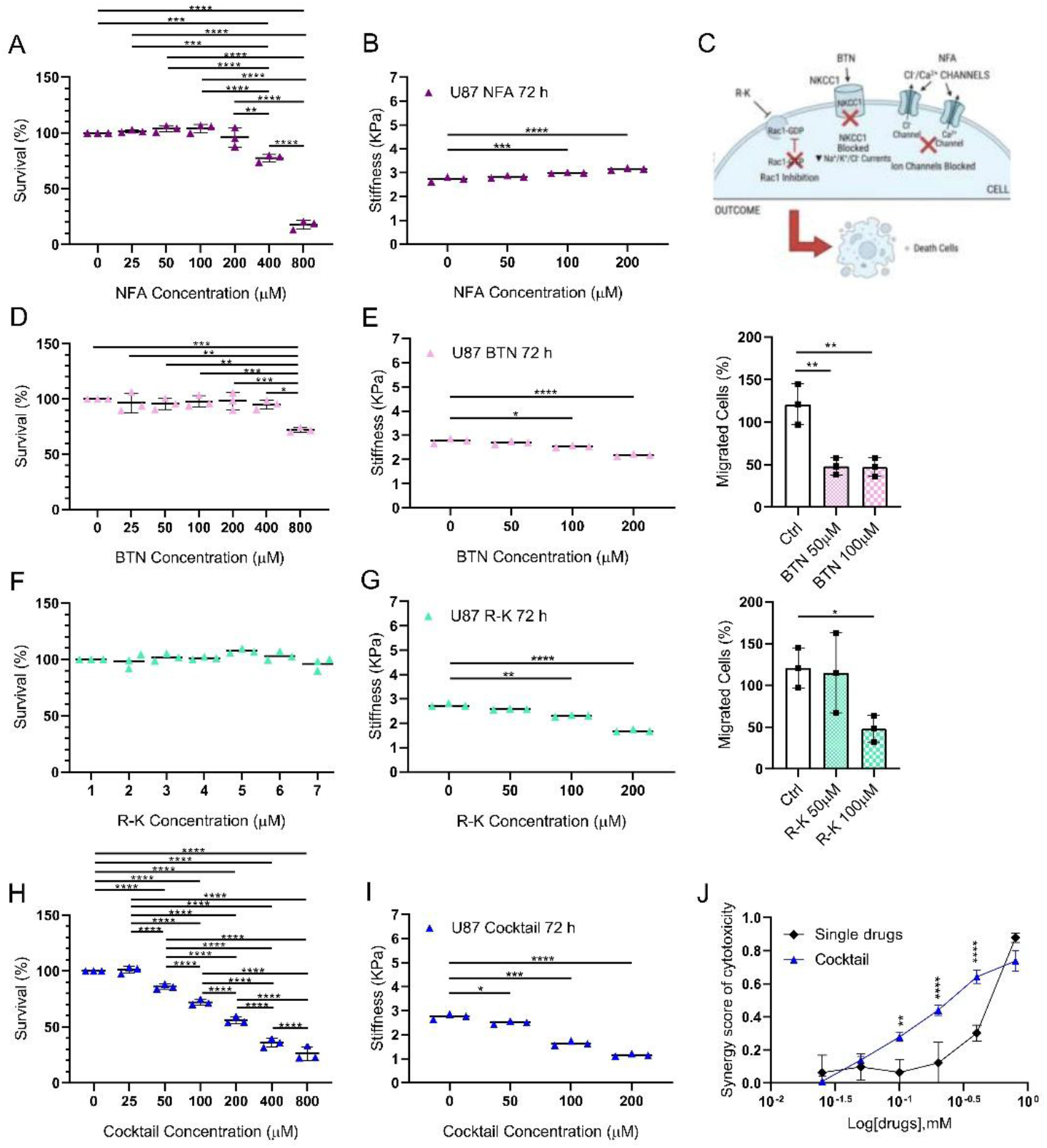
Effects of individual compounds and a multitarget cocktail on U87 cell viability and biomechanical properties. **(A–C)** Effects of niflumic acid (NFA). **(A)** Cell viability following 72 h of treatment, showing significant reductions at 400 and 800 μM (p = 0.0010 and p = 0.00013, respectively; one-way ANOVA followed by Sidak’s post hoc test). **(B)** Apparent Young’s modulus, showing significant increases at 100 and 200 μM (p = 0.00002 and p < 0.000001, respectively; one-way ANOVA followed by Tukey’s post hoc test). **(C)** Schematic rationale for the multitarget pharmacological strategy. **(D–E)** Effects of bumetanide (BTN). **(D)** Cell viability following 72 h of treatment, showing a significant reduction at 800 μM. **(E)** Apparent Young’s modulus (left), showing reductions at 100 and 200 μM (p = 0.0020 and p < 0.000001, respectively), and Transwell migration (right), showing reduced migration at 50 and 100 μM (p = 0.00280 and p = 0.0026, respectively). **(F–G)** Effects of R-ketorolac (R-K). **(F)** Cell viability following 72 h of treatment, showing no significant cytotoxicity. **(G)** Apparent Young’s modulus (left), showing reductions at 100 and 200 μM (both p < 0.000001), and Transwell migration (right), showing reduced migration at 100 μM (p = 0.0500). **(H–J)** Effects of the 1:1:1 NFA, BTN and R-K cocktail. **(H)** Cell viability following 72 h of treatment, showing significant reductions at 100 and 800 μM (both p = 0.0001). **(I)** Apparent Young’s modulus, showing reductions at 50 and 200 μM (p = 0.02 and p < 0.00001, respectively). **(J)** Bliss independence analysis indicating potential synergistic effects at 100, 200, and 400 μM (p = 0.00111, p = 0.000003, and p = 0.000001, respectively). Statistical results are provided in Supplementary Table 1. Data are presented as mean ± SEM. Statistical tests and post hoc analyses are indicated in the text. *p < 0.05; **p < 0.01; ***p < 0.001; ****p < 0.0001.

Although NFA has been reported to exert cytotoxic effects at lower concentrations in other cellular models (37, 38) its relatively high concentration requirement in our U87 cell viability assays prompted us to explore a complementary pharmacological strategy. Given that cellular volume regulation and mechanical stability involve multiple interacting pathways, we hypothesized that simultaneously targeting ion homeostasis and cytoskeletal organization could provide a more comprehensive approach than targeting a single component of these processes. Accordingly, we selected bumetanide, an inhibitor of the Na⁺-K⁺-2Cl⁻ and K⁺-Cl⁻ cotransporter, to interfere with chloride and intracellular ion homeostasis, and R-ketorolac, which has been associated with Rac1 inhibition and alterations in cytoskeletal regulation, to target cytoskeletal dynamics (Fig. 5C).

To characterize the individual effects of bumetanide and R-ketorolac on cell viability, biomechanical properties, and migratory capacity, we evaluated each compound separately in U87 cells. Bumetanide induced a moderate cytotoxic effect at the highest concentration tested, reducing cell viability to 72 ± 1.2% (Fig. 5D). Interestingly, biomechanical measurements revealed that bumetanide treatment decreased the Young’s modulus from 2.7 ± 0.06 kPa to 2.1 ± 0.03 kPa (Fig. 5E, left panel), indicating a reduction in cellular stiffness, in contrast to the increase observed following NFA treatment. To determine whether this treatment-induced softening was associated with altered migratory behavior, we performed transwell migration assays. Despite the reduction in cellular stiffness, treatment with 100 µM bumetanide significantly decreased cell migration, from 121 ± 13 % to 47 ± 6% (Fig. 5E, right panel). These findings indicate that bumetanide-mediated softening does not translate into increased migratory capacity and suggest that its effects on cell migration may involve mechanisms beyond changes in cellular stiffness alone.

In contrast, R-ketorolac did not significantly affect U87 cell viability at any of the concentrations tested (Fig. 5F). Nevertheless, its effects on cellular mechanics were more pronounced than those observed with bumetanide. Treatment with R-ketorolac reduced the Young’s modulus from 2.7 ± 0.04 kPa in control cells to 2.3 ± 0.02 kPa and 1.7 ± 0.03 kPa at 100 and 200 µM, respectively (Fig. 5G, left panel), indicating a marked decrease in cellular stiffness. To assess whether these biomechanical changes were accompanied by alterations in migratory behavior, we performed transwell migration assays. Consistent with the results obtained with bumetanide, R-ketorolac treatment at the tested concentration significantly reduced cell migration compared with control conditions from 121 ± 13 % to 48 ± 9% (Fig. 5G, right panel). Together, these findings demonstrate that both bumetanide and R-ketorolac reduce cellular stiffness while impairing migratory capacity, despite their distinct effects on cell viability.

Based on the results obtained with the individual compounds, we subsequently treated U87 cells with a pharmacological cocktail comprising NFA, bumetanide, and R-ketorolac at a 1:1:1 ratio to investigate whether the combined modulation of ion transport and cytoskeletal organization could enhance the effects on cell viability and biomechanical properties. The combined treatment significantly reduced cell viability across the tested concentration range of 50–800 µM, producing a greater reduction in survival than that observed with the individual compounds (Fig. 5H). In parallel, biomechanical measurements revealed a pronounced decrease in cellular stiffness following cocktail treatment. Notably, a significant reduction was already observed at 50 µM, with the Young’s modulus decreasing from 2.7 ± 0.07 kPa in control cells to 2.5 ± 0.04 kPa in treated cells (Fig. 5I).

To investigate whether the enhanced effects of the pharmacological cocktail could be attributed to synergistic interactions among its components, we applied the Bliss independence model to compare the experimentally observed effects on cell viability with those predicted under the assumption of independent drug action (Fig. 5J). Across cocktail concentrations ranging from 100 to 400 µM, the observed effects exceeded the Bliss-predicted values, with corresponding Bliss scores of 0.06, 0.12, and 0.3 and observed effects of 0.27, 0.44, and 0.64 at the respective concentrations. These results support synergistic interaction among NFA, bumetanide, and R-ketorolac within this concentration range, providing a rationale for investigating the combined treatment in patient-derived glioma stem cells (GSCs).

### GSCs showed a greater sensitivity to a multi-target strategy cocktail

We next assessed the efficacy of the cocktail in GSC cultures derived from 11 patients, using an initial concentration of 100 µM, corresponding to the lowest concentration at which a synergistic effect was observed in U87 cells. Time-lapse imaging over 72 h was used to quantify mitotic events in each culture (Supplementary Fig. 7A). Five of the 11 cultures (45%) exhibited a significant response to treatment, revealing substantial interpatient variability in sensitivity.

To explore the relationship between electrophysiological properties and treatment response (Supplementary Fig. 7B), we compared VRAC-like currents across patient-derived cultures. Cocktail-responsive cultures S557, S566, and S610 generally exhibited larger VRAC-like currents, although S565 was an exception. In contrast, VRAC-like current amplitudes varied among non-responsive cultures. These findings should be interpreted in the context of the substantial interpatient variability in NFA-mediated inhibition (see Fig.2), which complicates the establishment of a direct association between VRAC-like current amplitude and cocktail responsiveness at 100 uM.

Based on these findings, we selected the S82 patient-derived GSC line, which exhibits a prominent VRAC-like current and no evidence of NCX exchanger current (Supplementary Fig. 8A), for further dose–response characterization. S82 cells displayed sensitivity to the cocktail comparable to that observed in U87 cells, with cell viability decreasing to 52 ± 1.04% following treatment with 100 µM of the cocktail (Fig. 6A). Notably, we had previously characterized a calcium-dependent VRAC-like current in this GSC line, which was inhibited by approximately 85% upon application of 100 µM NFA (Supplementary Fig. 8B). This finding provides a functional link between the electrophysiological properties of the selected GSC model and the pharmacological target investigated in this study.

**Figure 6.**
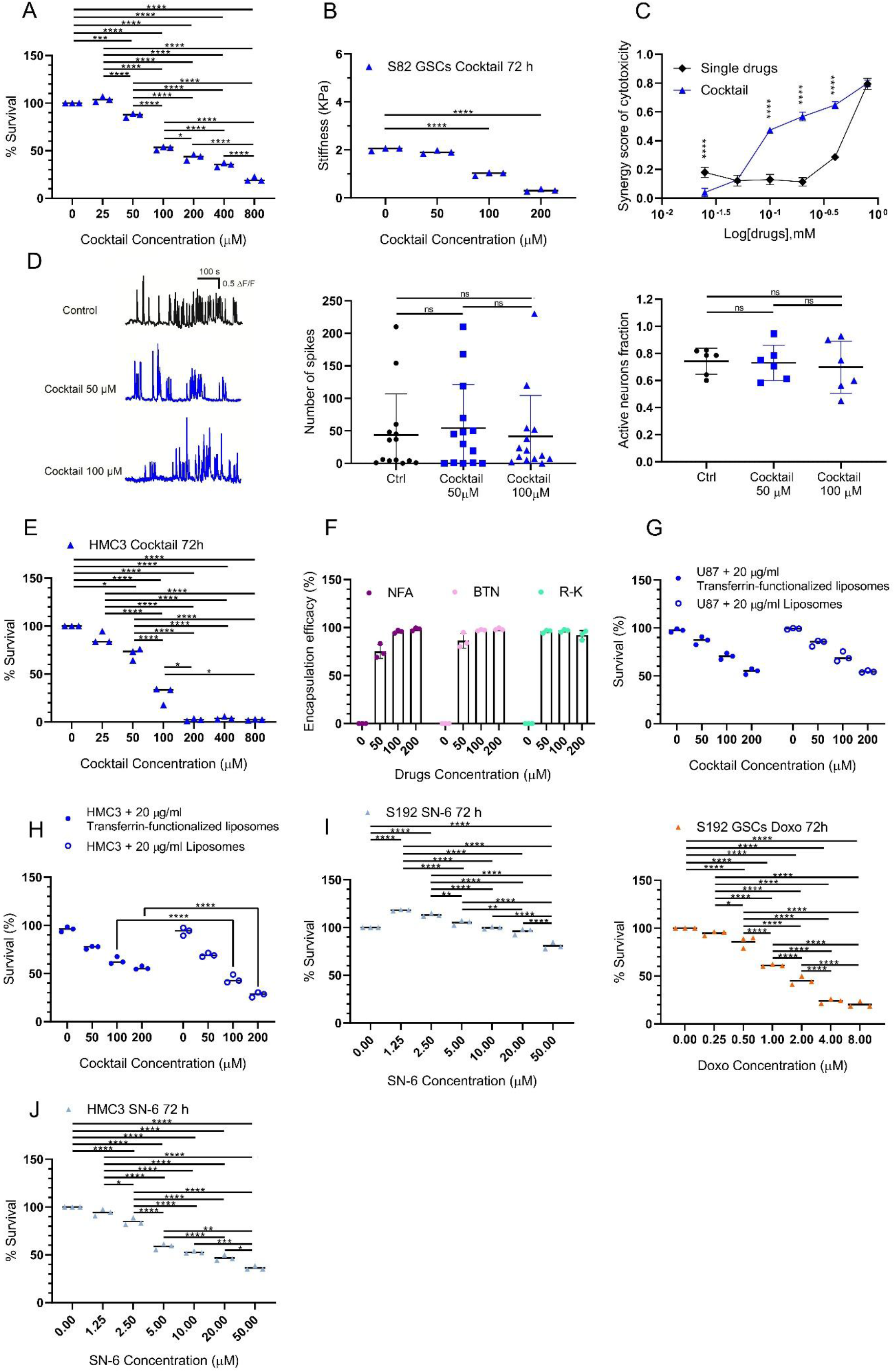
Evaluation of the therapeutic effects of the drug cocktail, its impact on neuronal activity, and the efficacy and selectivity of liposome-encapsulated formulations. **(A–C)** Effects of the 1:1:1 cocktail of niflumic acid (NFA), bumetanide, and R-ketorolac. **(A)** Cell viability following 72 h of treatment, showing a significant reduction at 100 and 800 μM (both p = 0.00001). **(B)** Apparent Young’s modulus following cocktail treatment, showing a significant reduction at 100 and 200 μM (both p < 0.000001; one-way ANOVA followed by Tukey’s post hoc test). **(C)** Bliss independence analysis indicating effects greater than those predicted for individual compounds at 100, 200, and 400 μM (p = 0.000007, p = 0.000001, and p = 0.000001, respectively; two-way ANOVA followed by Tukey’s post hoc test). **(D)** Assessment of spontaneous neuronal activity by calcium imaging in primary mouse hippocampal neurons. Left: representative calcium traces under the experimental conditions tested. Middle: frequency of spontaneous calcium transients recorded over 10 min following treatment with 50 or 100 μM cocktail, showing no significant differences compared with vehicle-treated controls. Right: percentage of neurons displaying spontaneous calcium activity following treatment with 100 μM cocktail (70 ± 0.08%), with no significant difference compared with controls. **(E)** HMC3 cell viability following 72 h of treatment with the 1:1:1 cocktail, showing concentration-dependent cytotoxicity (p = 0.00127 at 25 μM and p < 0.00001 at 200 μM; one-way ANOVA followed by Sidak’s post hoc test). **(F–H)** Characterization and biological effects of cocktail-loaded liposomes. **(F)** Drug encapsulation efficiency determined by UV– Vis spectrophotometry following liposome disruption, using absorbance wavelengths of 230 nm for bumetanide (BTN), 290 nm for NFA, and 315 nm for R-ketorolac (R-K). **(G)** U87 cell viability following treatment with cocktail-loaded non-functionalized or transferrin-functionalized liposomes (20 μg/mL), showing no significant difference between formulations (two-way ANOVA followed by Tukey’s post hoc test). **(H)** HMC3 cell viability under the same treatment conditions, showing significantly lower cytotoxicity with transferrin-functionalized liposomes compared with non-functionalized liposomes (p < 0.00001; two-way ANOVA followed by Tukey’s post hoc test). **(I)** Effects of SN-6 and doxorubicin on S192-derived GSC viability following 72 h of treatment. Left: concentration-dependent effects of SN-6 (0–50 μM; one-way ANOVA followed by Sidak’s post hoc test). Right: doxorubicin-induced reduction in cell viability at concentrations of 0.5–8 μM compared with untreated controls (p = 0.000014 and p < 0.000001, respectively; one-way ANOVA followed by Sidak’s post hoc test).**(J)** HMC3 cell viability following 72 h of SN-6 treatment, showing significant cytotoxicity at concentrations of 1–50 μM (p = 0.000095 and p < 0.00001, respectively; one-way ANOVA followed by Sidak’s post hoc test). Statistical results for the comparisons shown are provided in Table 1. Data are presented as mean ± SEM. Statistical significance was assessed using paired tests, where appropriate. *p < 0.05; **p < 0.01; ***p < 0.001; ****p < 0.0001.

**Table 1.**
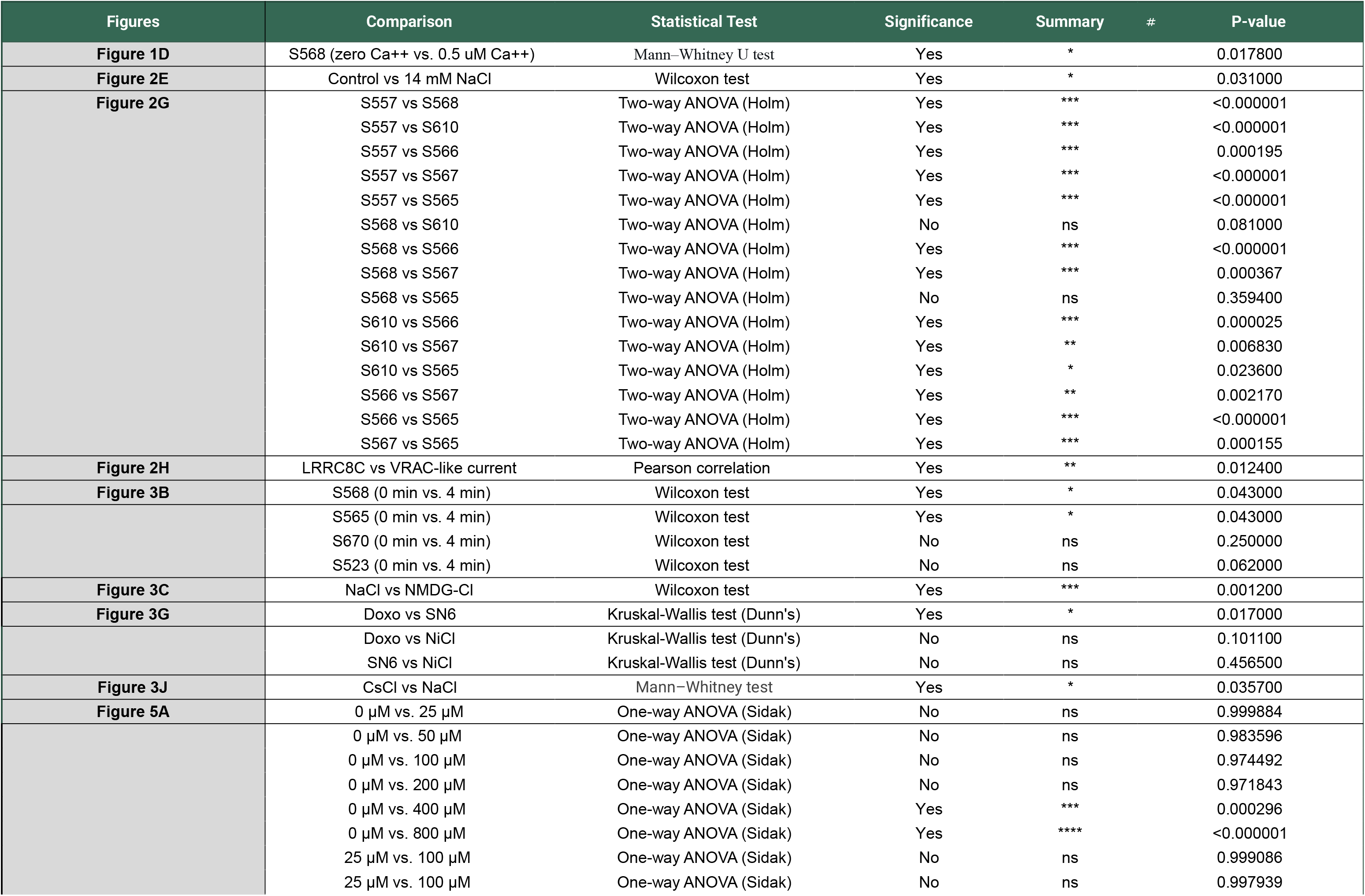
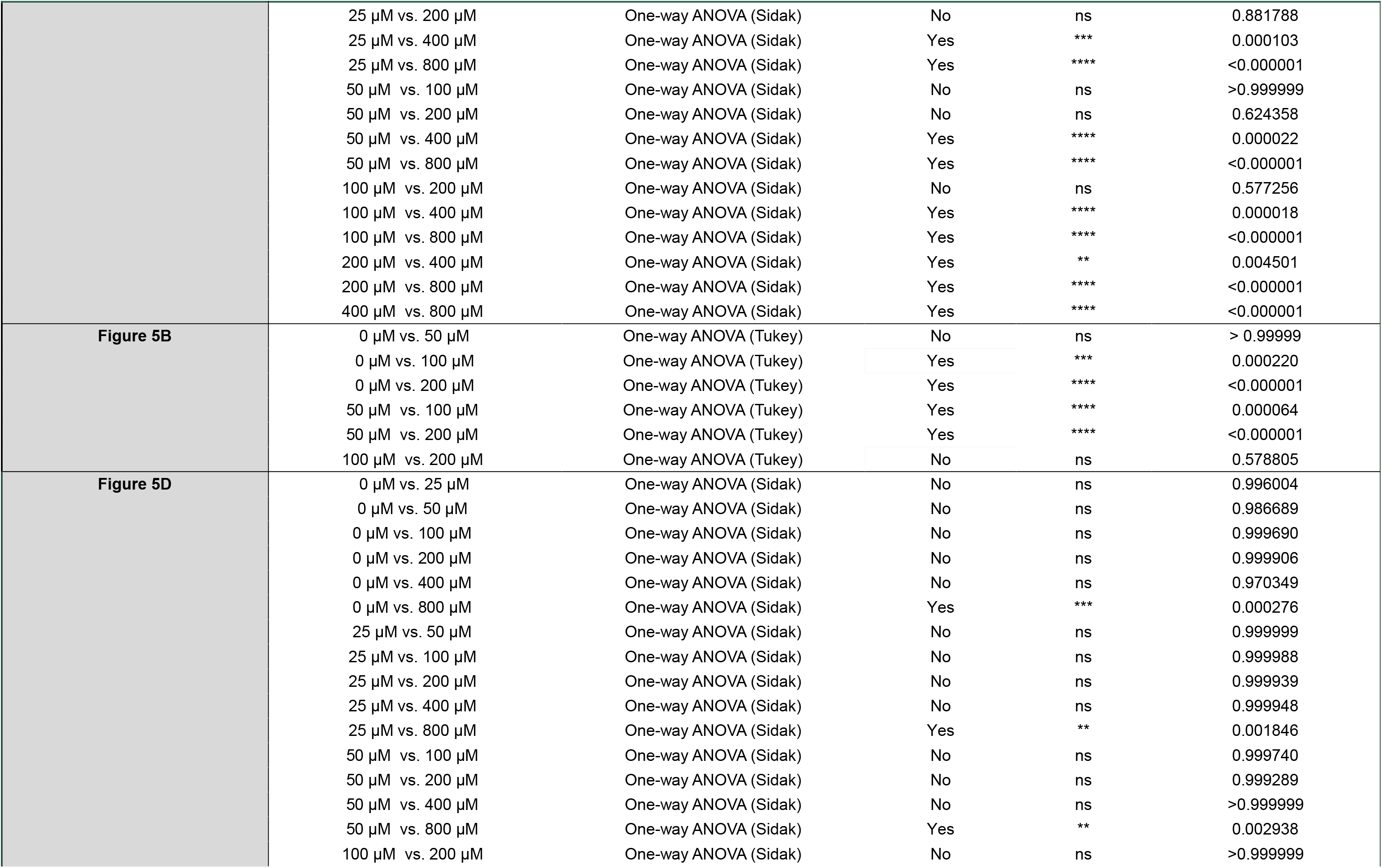
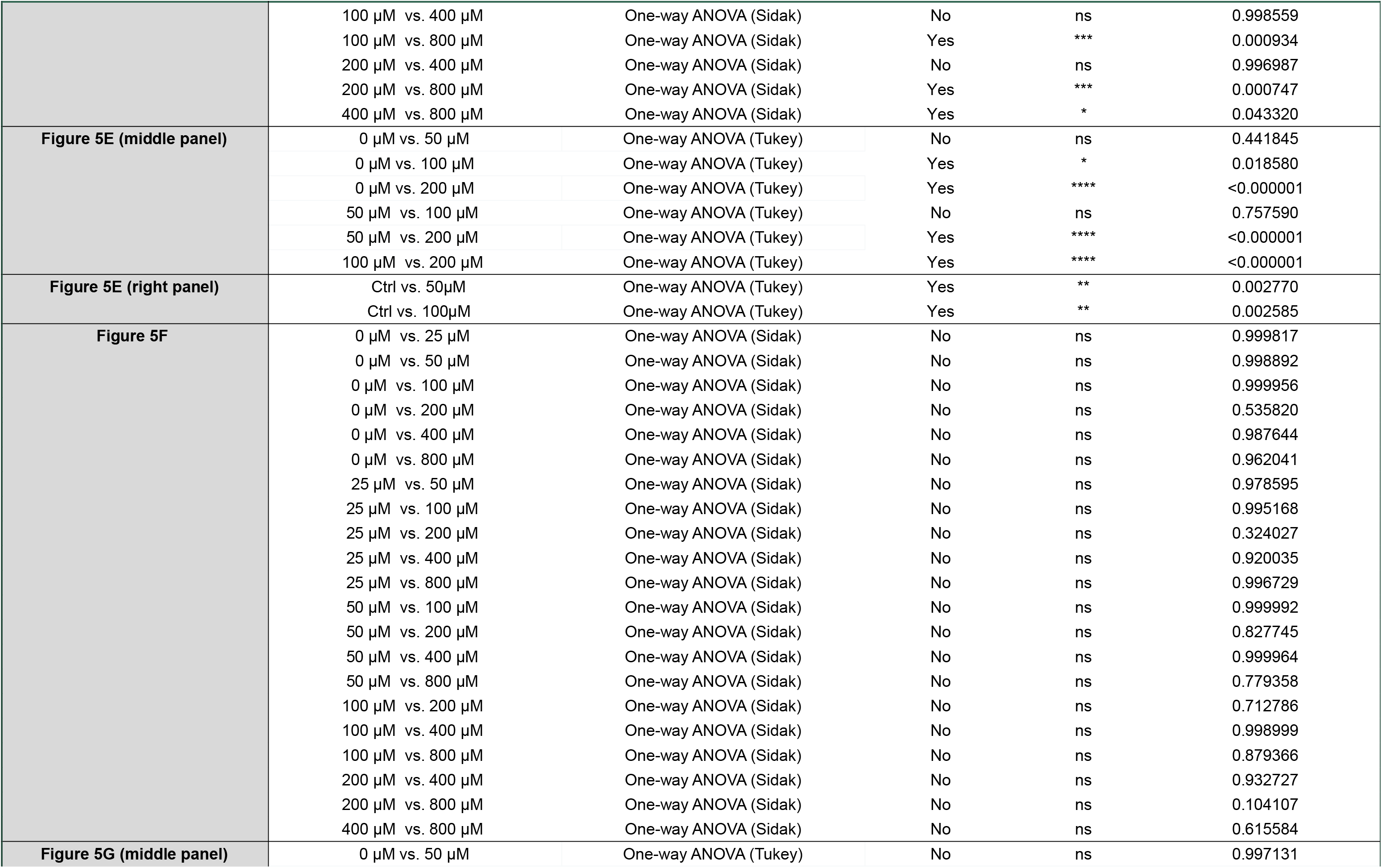
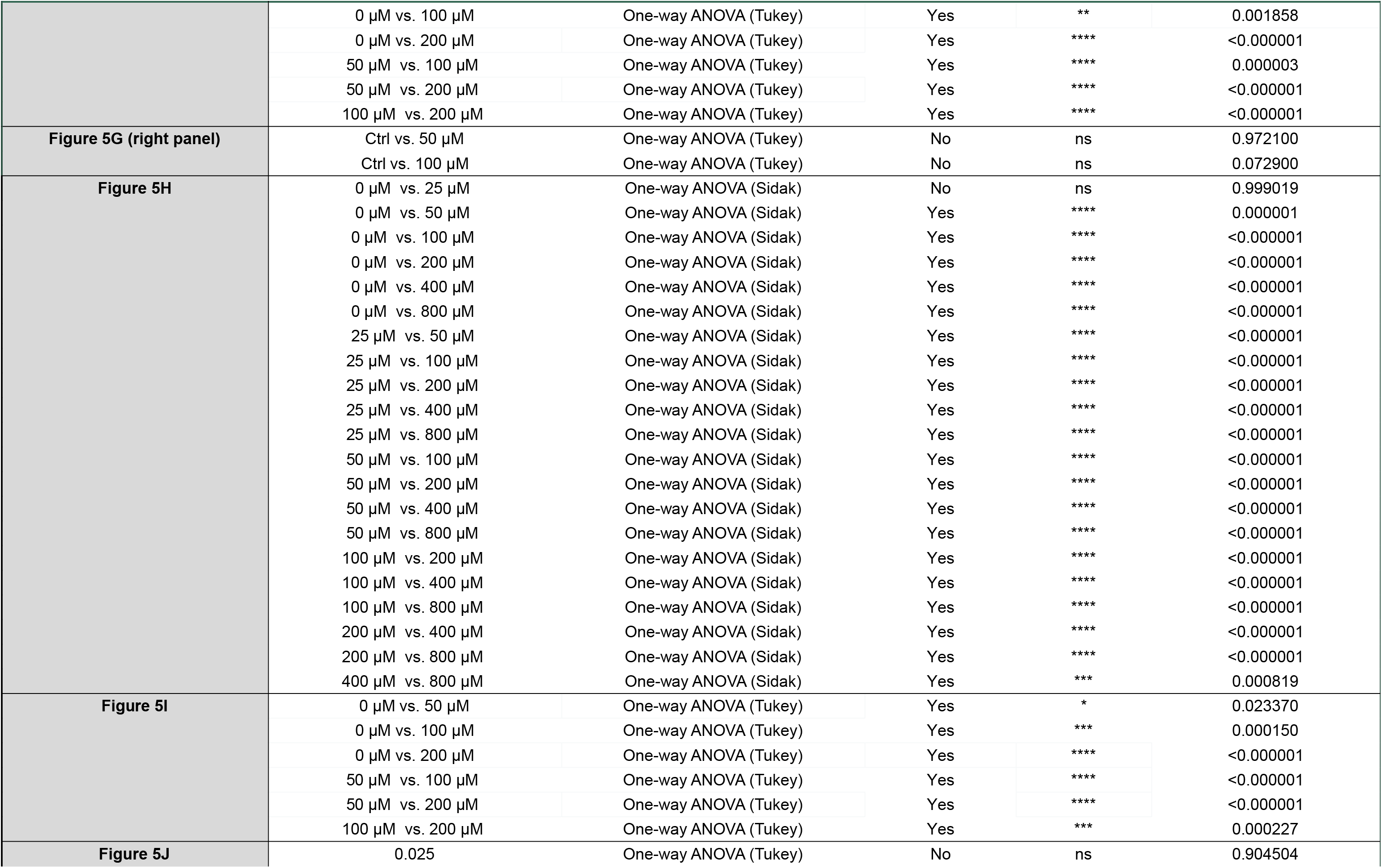
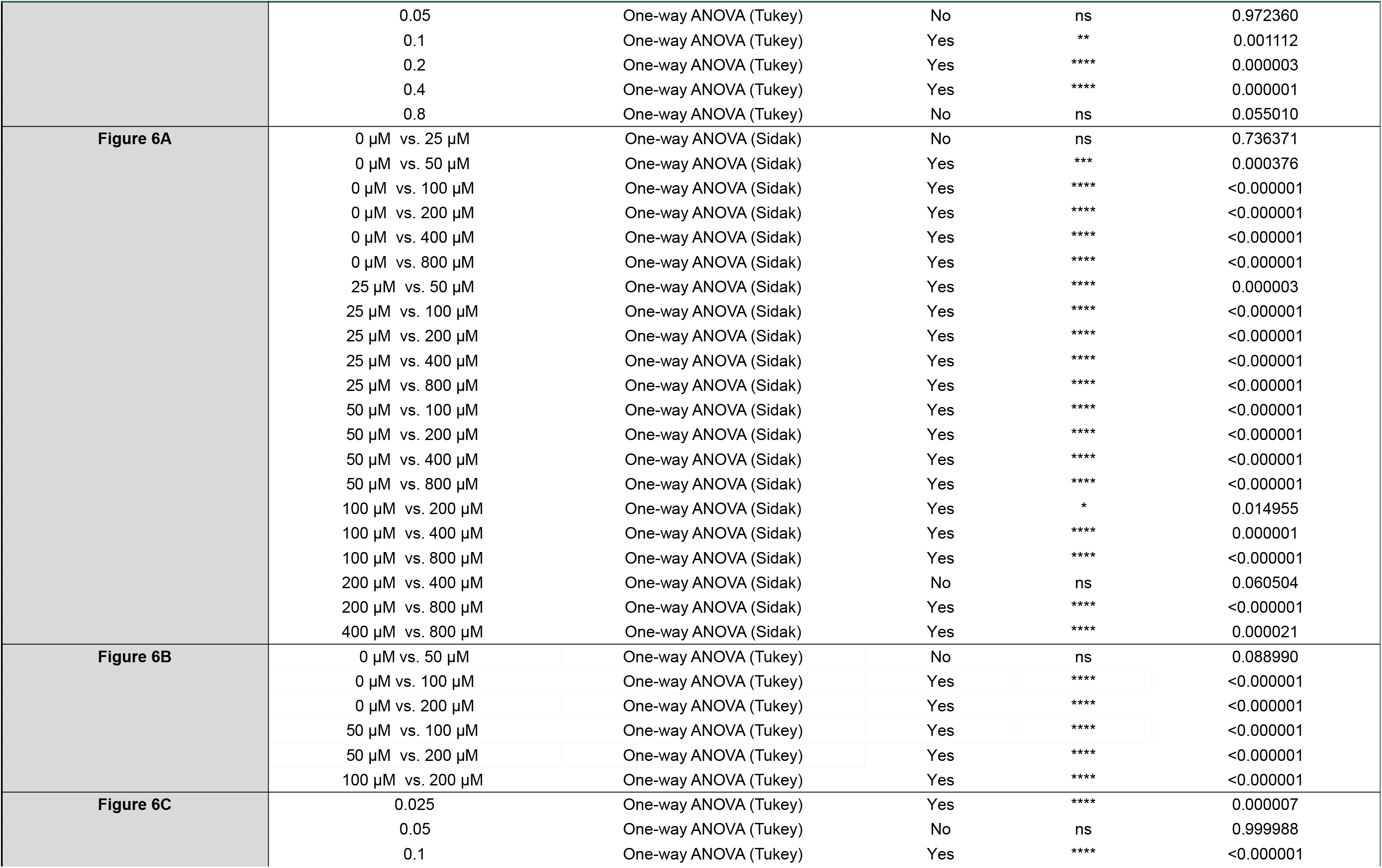
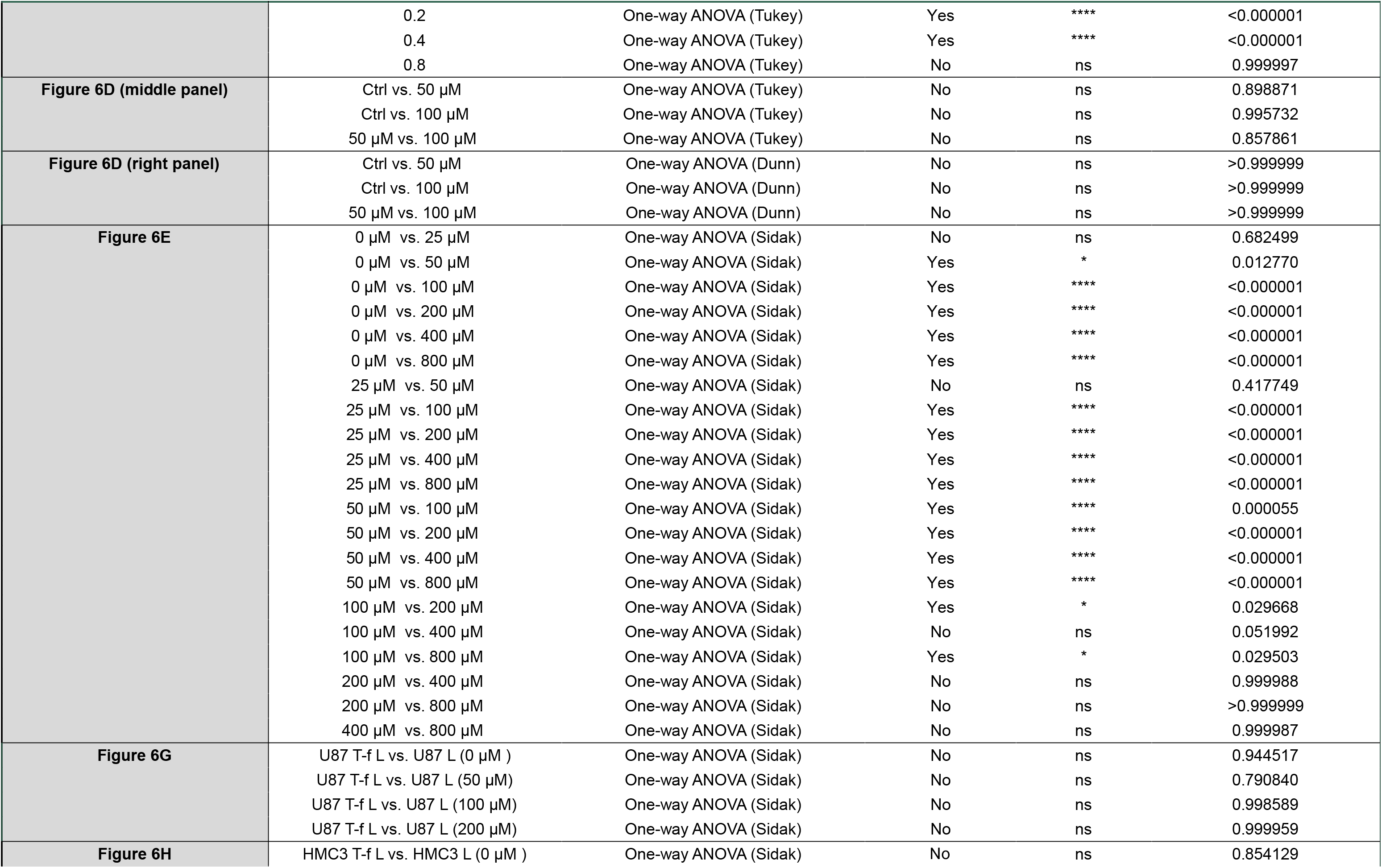
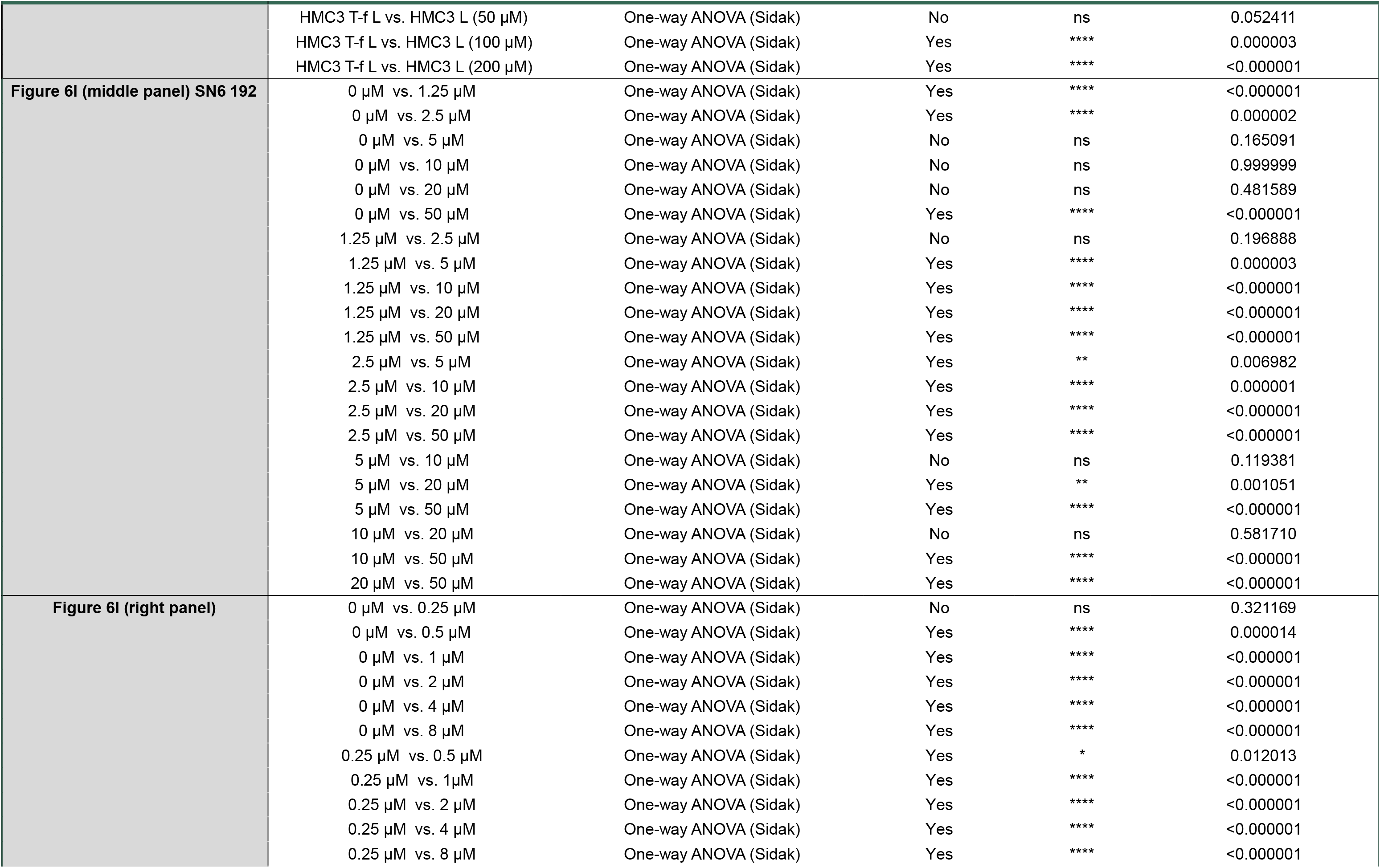
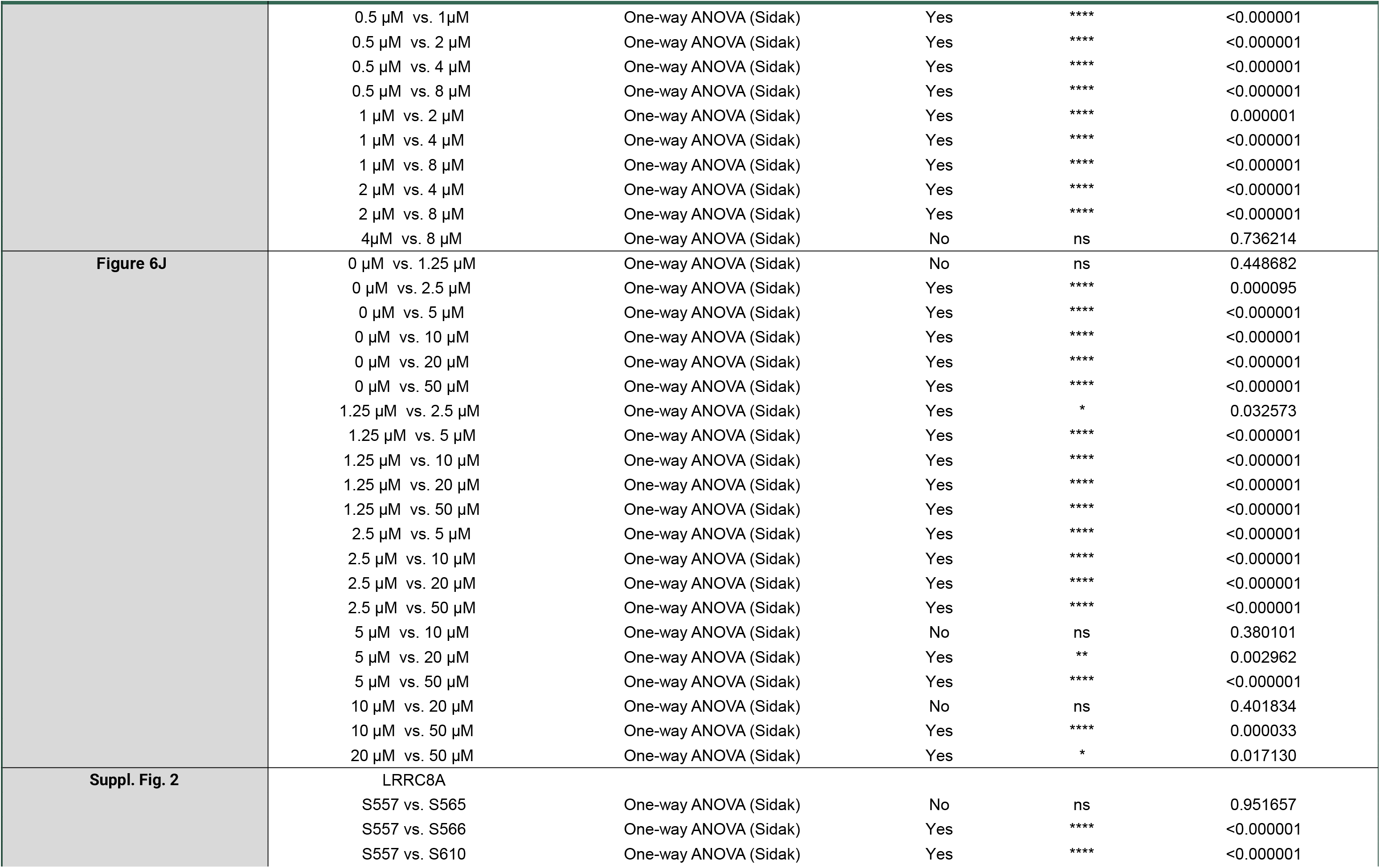
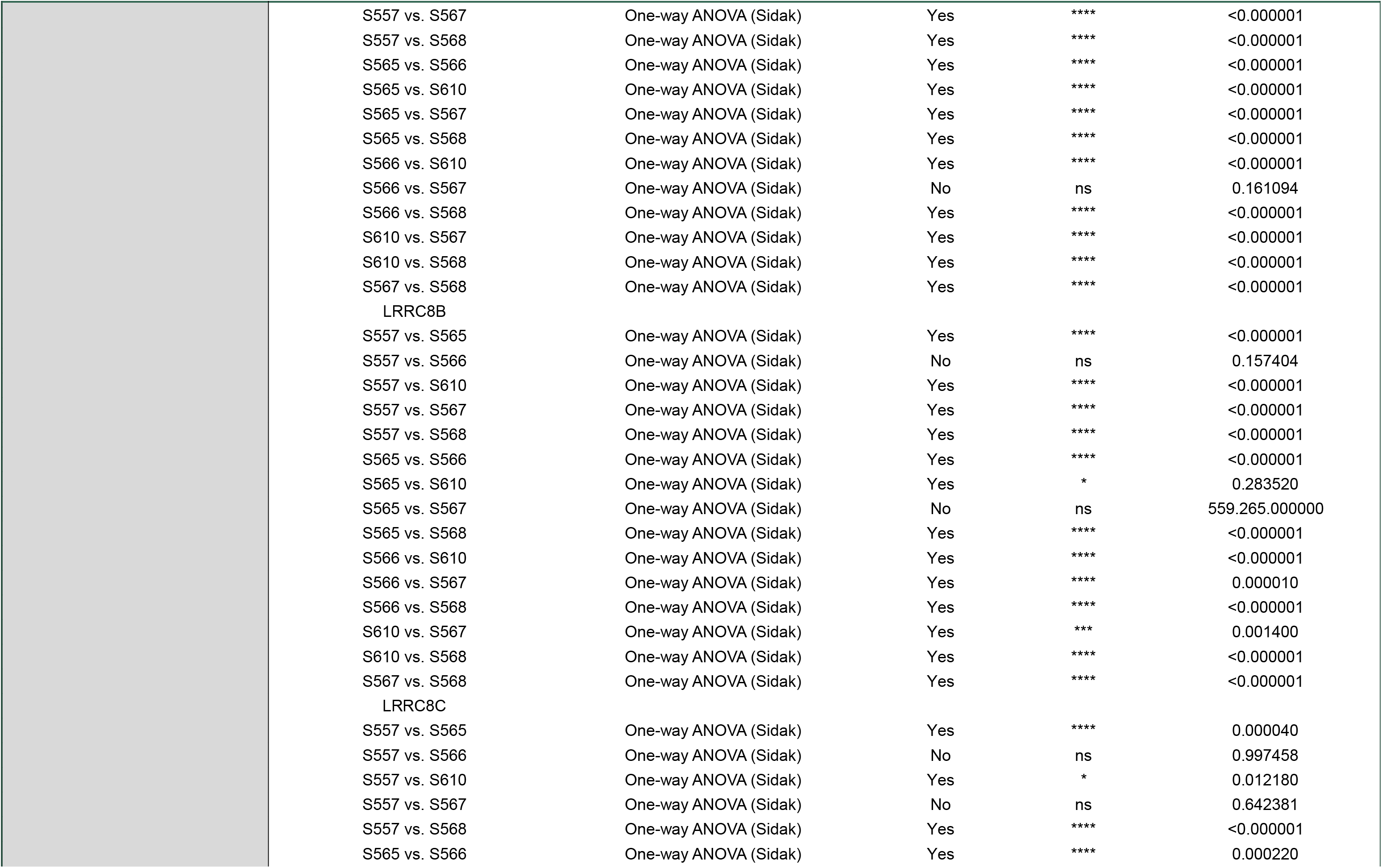
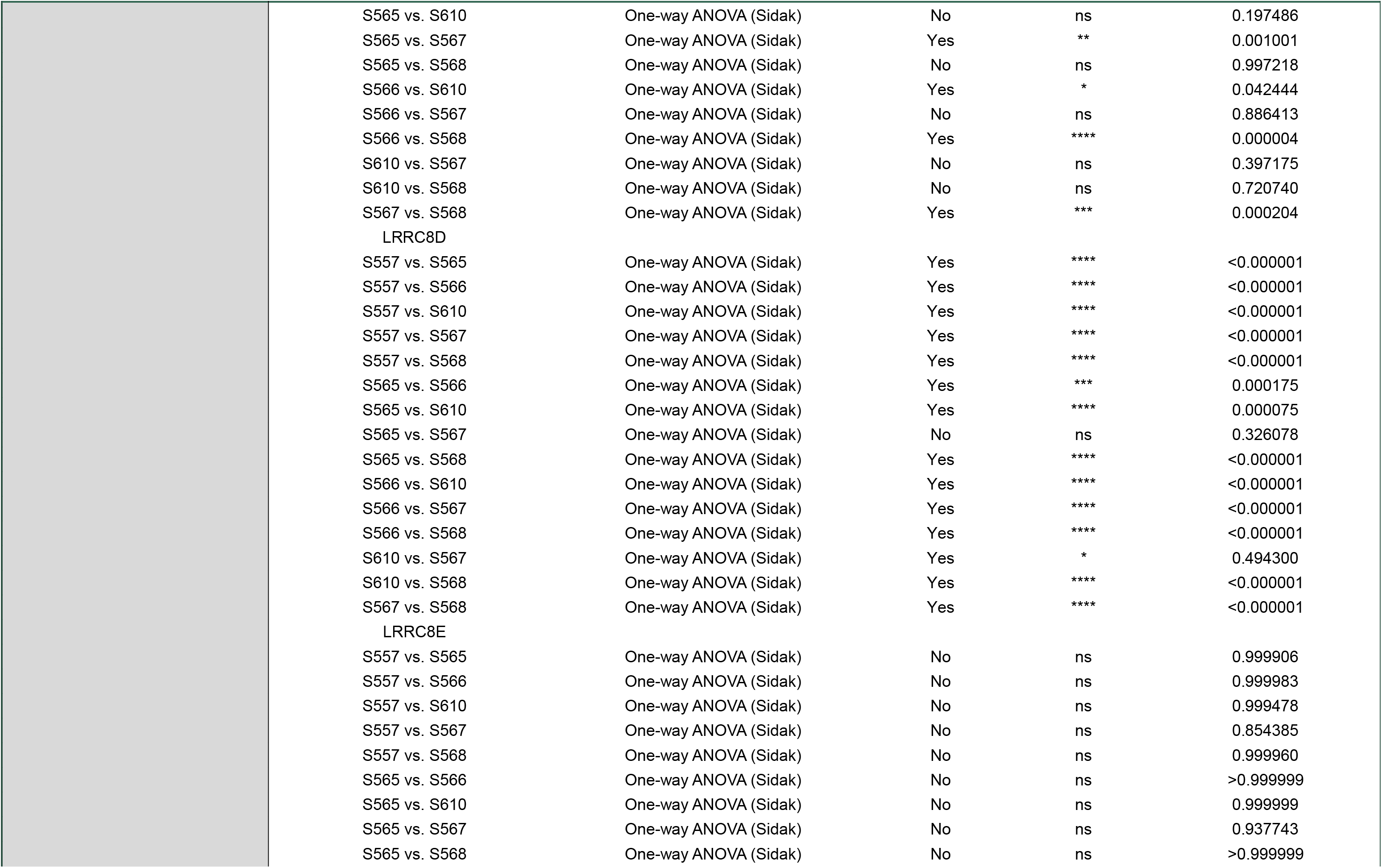
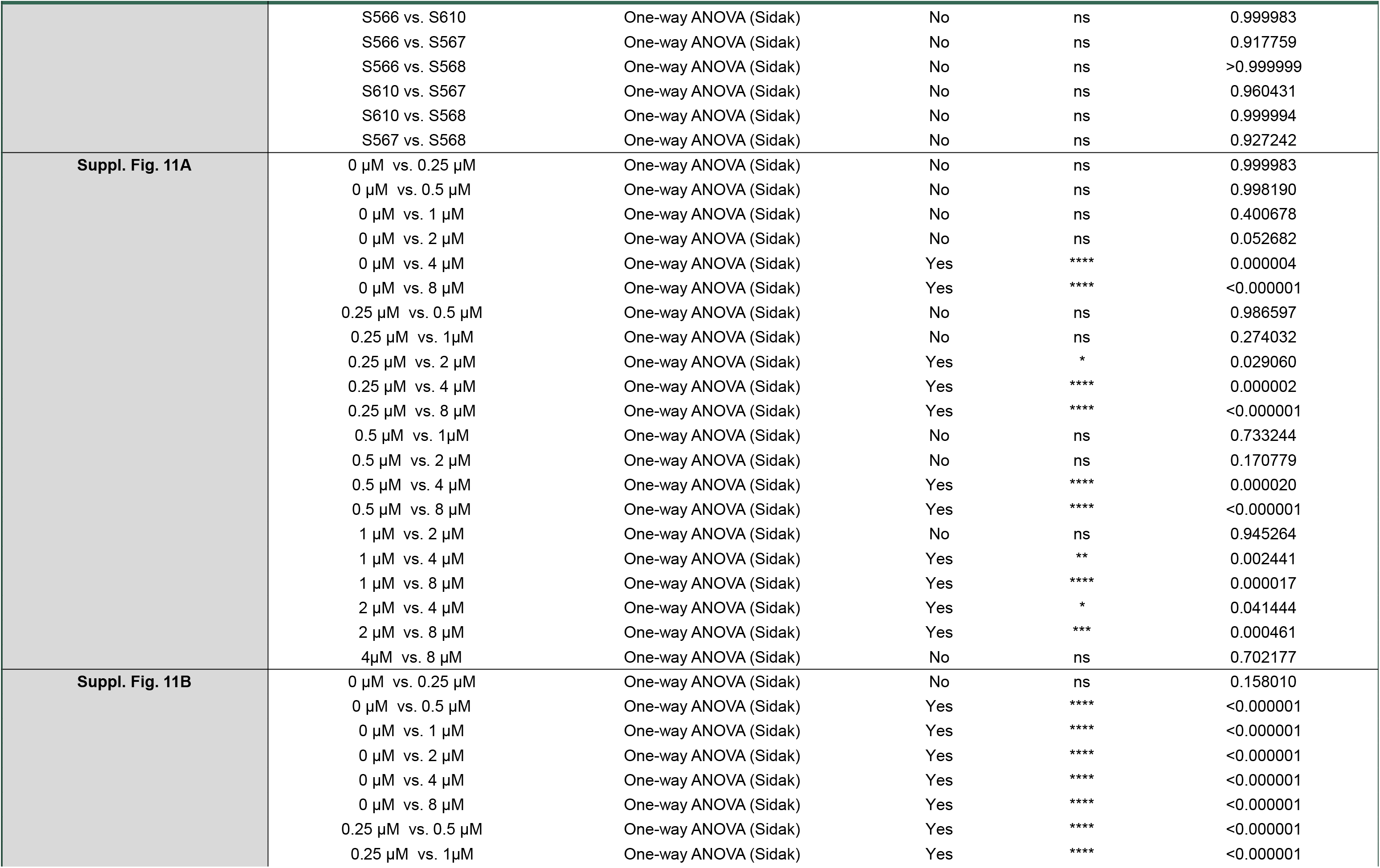
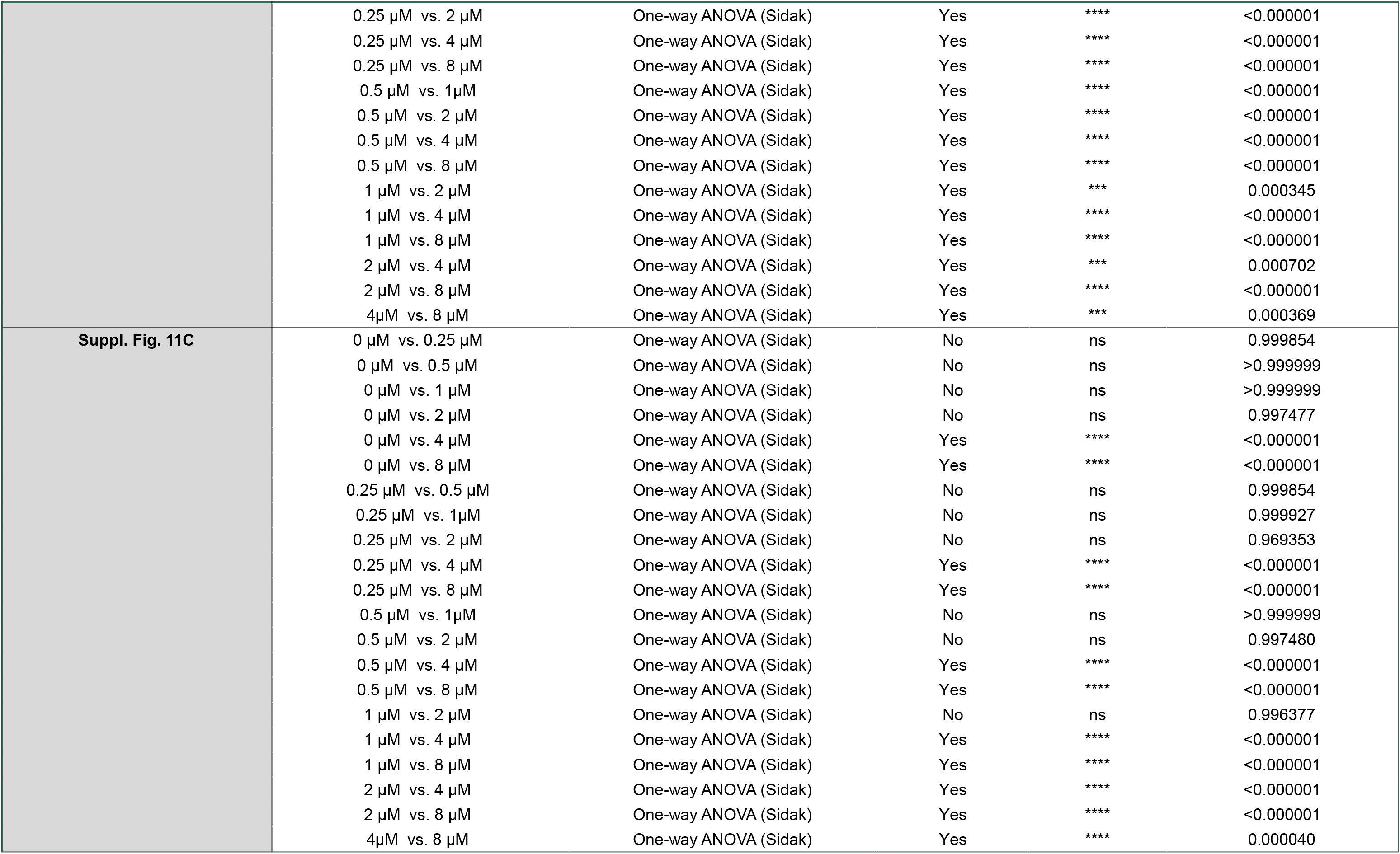

| Figures | Comparison | Statistical Test | Significance | Summary # | P-value |
|---|---|---|---|---|---|
| **Figure 1D** | S568 (zero Ca++ vs. 0.5 uM Ca++) | Mann–Whitney U test | Yes | * | 0.017800 |
| **Figure 2E** | Control vs 14 mM NaCl | Wilcoxon test | Yes | * | 0.031000 |
| **Figure 2G** | S557 vs S568 | Two-way ANOVA (Holm) | Yes | *** | <0.000001 |
|  | S557 vs S610 | Two-way ANOVA (Holm) | Yes | *** | <0.000001 |
|  | S557 vs S566 | Two-way ANOVA (Holm) | Yes | *** | 0.000195 |
|  | S557 vs S567 | Two-way ANOVA (Holm) | Yes | *** | <0.000001 |
|  | S557 vs S565 | Two-way ANOVA (Holm) | Yes | *** | <0.000001 |
|  | S568 vs S610 | Two-way ANOVA (Holm) | No | ns | 0.081000 |
|  | S568 vs S566 | Two-way ANOVA (Holm) | Yes | *** | <0.000001 |
|  | S568 vs S567 | Two-way ANOVA (Holm) | Yes | *** | 0.000367 |
|  | S568 vs S565 | Two-way ANOVA (Holm) | No | ns | 0.359400 |
|  | S610 vs S566 | Two-way ANOVA (Holm) | Yes | *** | 0.000025 |
|  | S610 vs S567 | Two-way ANOVA (Holm) | Yes | ** | 0.006830 |
|  | S610 vs S565 | Two-way ANOVA (Holm) | Yes | * | 0.023600 |
|  | S566 vs S567 | Two-way ANOVA (Holm) | Yes | ** | 0.002170 |
|  | S566 vs S565 | Two-way ANOVA (Holm) | Yes | *** | <0.000001 |
|  | S567 vs S565 | Two-way ANOVA (Holm) | Yes | *** | 0.000155 |
| **Figure 2H** | LRRC8C vs VRAC-like current | Pearson correlation | Yes | ** | 0.012400 |
| **Figure 3B** | S568 (0 min vs. 4 min) | Wilcoxon test | Yes | * | 0.043000 |
|  | S565 (0 min vs. 4 min) | Wilcoxon test | Yes | * | 0.043000 |
|  | S670 (0 min vs. 4 min) | Wilcoxon test | No | ns | 0.250000 |
|  | S523 (0 min vs. 4 min) | Wilcoxon test | No | ns | 0.062000 |
| **Figure 3C** | NaCl vs NMDG-Cl | Wilcoxon test | Yes | *** | 0.001200 |
| **Figure 3G** | Doxo vs SN6 | Kruskal-Wallis test (Dunn's) | Yes | * | 0.017000 |
|  | Doxo vs NiCl | Kruskal-Wallis test (Dunn's) | No | ns | 0.101100 |
|  | SN6 vs NiCl | Kruskal-Wallis test (Dunn's) | No | ns | 0.456500 |
| **Figure 3J** | CsCl vs NaCl | Mann–Whitney test | Yes | * | 0.035700 |
| **Figure 5A** | 0 µM vs. 25 µM | One-way ANOVA (Sidak) | No | ns | 0.999884 |
|  | 0 µM vs. 50 µM | One-way ANOVA (Sidak) | No | ns | 0.983596 |
|  | 0 µM vs. 100 µM | One-way ANOVA (Sidak) | No | ns | 0.974492 |
|  | 0 µM vs. 200 µM | One-way ANOVA (Sidak) | No | ns | 0.971843 |
|  | 0 µM vs. 400 µM | One-way ANOVA (Sidak) | Yes | *** | 0.000296 |
|  | 0 µM vs. 800 µM | One-way ANOVA (Sidak) | Yes | **** | <0.000001 |
|  | 25 µM vs. 100 µM | One-way ANOVA (Sidak) | No | ns | 0.999086 |
|  | 25 µM vs. 100 µM | One-way ANOVA (Sidak) | No | ns | 0.997939 |
|  | 25 µM vs. 200 µM | One-way ANOVA (Sidak) | No | ns | 0.881788 |
|  | 25 µM vs. 400 µM | One-way ANOVA (Sidak) | Yes | *** | 0.000103 |
|  | 25 µM vs. 800 µM | One-way ANOVA (Sidak) | Yes | **** | <0.000001 |
|  | 50 µM vs. 100 µM | One-way ANOVA (Sidak) | No | ns | >0.999999 |
|  | 50 µM vs. 200 µM | One-way ANOVA (Sidak) | No | ns | 0.624358 |
|  | 50 µM vs. 400 µM | One-way ANOVA (Sidak) | Yes | **** | 0.000022 |
|  | 50 µM vs. 800 µM | One-way ANOVA (Sidak) | Yes | **** | <0.000001 |
|  | 100 µM vs. 200 µM | One-way ANOVA (Sidak) | No | ns | 0.577256 |
|  | 100 µM vs. 400 µM | One-way ANOVA (Sidak) | Yes | **** | 0.000018 |
|  | 100 µM vs. 800 µM | One-way ANOVA (Sidak) | Yes | **** | <0.000001 |
|  | 200 µM vs. 400 µM | One-way ANOVA (Sidak) | Yes | ** | 0.004501 |
|  | 200 µM vs. 800 µM | One-way ANOVA (Sidak) | Yes | **** | <0.000001 |
|  | 400 µM vs. 800 µM | One-way ANOVA (Sidak) | Yes | **** | <0.000001 |
| **Figure 5B** | 0 µM vs. 50 µM | One-way ANOVA (Tukey) | No | ns | > 0.99999 |
|  | 0 µM vs. 100 µM | One-way ANOVA (Tukey) | Yes | *** | 0.000220 |
|  | 0 µM vs. 200 µM | One-way ANOVA (Tukey) | Yes | **** | <0.000001 |
|  | 50 µM vs. 100 µM | One-way ANOVA (Tukey) | Yes | **** | 0.000064 |
|  | 50 µM vs. 200 µM | One-way ANOVA (Tukey) | Yes | **** | <0.000001 |
|  | 100 µM vs. 200 µM | One-way ANOVA (Tukey) | No | ns | 0.578805 |
| **Figure 5D** | 0 µM vs. 25 µM | One-way ANOVA (Sidak) | No | ns | 0.996004 |
|  | 0 µM vs. 50 µM | One-way ANOVA (Sidak) | No | ns | 0.986689 |
|  | 0 µM vs. 100 µM | One-way ANOVA (Sidak) | No | ns | 0.999690 |
|  | 0 µM vs. 200 µM | One-way ANOVA (Sidak) | No | ns | 0.999906 |
|  | 0 µM vs. 400 µM | One-way ANOVA (Sidak) | No | ns | 0.970349 |
|  | 0 µM vs. 800 µM | One-way ANOVA (Sidak) | Yes | *** | 0.000276 |
|  | 25 µM vs. 50 µM | One-way ANOVA (Sidak) | No | ns | 0.999999 |
|  | 25 µM vs. 100 µM | One-way ANOVA (Sidak) | No | ns | 0.999988 |
|  | 25 µM vs. 200 µM | One-way ANOVA (Sidak) | No | ns | 0.999939 |
|  | 25 µM vs. 400 µM | One-way ANOVA (Sidak) | No | ns | 0.999948 |
|  | 25 µM vs. 800 µM | One-way ANOVA (Sidak) | Yes | ** | 0.001846 |
|  | 50 µM vs. 100 µM | One-way ANOVA (Sidak) | No | ns | 0.999740 |
|  | 50 µM vs. 200 µM | One-way ANOVA (Sidak) | No | ns | 0.999289 |
|  | 50 µM vs. 400 µM | One-way ANOVA (Sidak) | No | ns | >0.999999 |
|  | 50 µM vs. 800 µM | One-way ANOVA (Sidak) | Yes | ** | 0.002938 |
|  | 100 µM vs. 200 µM | One-way ANOVA (Sidak) | No | ns | >0.999999 |
|  | 100 µM vs. 400 µM | One-way ANOVA (Sidak) | No | ns | 0.998559 |
|  | 100 µM vs. 800 µM | One-way ANOVA (Sidak) | Yes | *** | 0.000934 |
|  | 200 µM vs. 400 µM | One-way ANOVA (Sidak) | No | ns | 0.996987 |
|  | 200 µM vs. 800 µM | One-way ANOVA (Sidak) | Yes | *** | 0.000747 |
|  | 400 µM vs. 800 µM | One-way ANOVA (Sidak) | Yes | * | 0.043320 |
| **Figure 5E (middle panel)** | 0 µM vs. 50 µM | One-way ANOVA (Tukey) | No | ns | 0.441845 |
|  | 0 µM vs. 100 µM | One-way ANOVA (Tukey) | Yes | * | 0.018580 |
|  | 0 µM vs. 200 µM | One-way ANOVA (Tukey) | Yes | **** | <0.000001 |
|  | 50 µM vs. 100 µM | One-way ANOVA (Tukey) | No | ns | 0.757590 |
|  | 50 µM vs. 200 µM | One-way ANOVA (Tukey) | Yes | **** | <0.000001 |
|  | 100 µM vs. 200 µM | One-way ANOVA (Tukey) | Yes | **** | <0.000001 |
| **Figure 5E (right panel)** | Ctrl vs. 50µM | One-way ANOVA (Tukey) | Yes | ** | 0.002770 |
|  | Ctrl vs. 100µM | One-way ANOVA (Tukey) | Yes | ** | 0.002585 |
| **Figure 5F** | 0 µM vs. 25 µM | One-way ANOVA (Sidak) | No | ns | 0.999817 |
|  | 0 µM vs. 50 µM | One-way ANOVA (Sidak) | No | ns | 0.998892 |
|  | 0 µM vs. 100 µM | One-way ANOVA (Sidak) | No | ns | 0.999956 |
|  | 0 µM vs. 200 µM | One-way ANOVA (Sidak) | No | ns | 0.535820 |
|  | 0 µM vs. 400 µM | One-way ANOVA (Sidak) | No | ns | 0.987644 |
|  | 0 µM vs. 800 µM | One-way ANOVA (Sidak) | No | ns | 0.962041 |
|  | 25 µM vs. 50 µM | One-way ANOVA (Sidak) | No | ns | 0.978595 |
|  | 25 µM vs. 100 µM | One-way ANOVA (Sidak) | No | ns | 0.995168 |
|  | 25 µM vs. 200 µM | One-way ANOVA (Sidak) | No | ns | 0.324027 |
|  | 25 µM vs. 400 µM | One-way ANOVA (Sidak) | No | ns | 0.920035 |
|  | 25 µM vs. 800 µM | One-way ANOVA (Sidak) | No | ns | 0.996729 |
|  | 50 µM vs. 100 µM | One-way ANOVA (Sidak) | No | ns | 0.999992 |
|  | 50 µM vs. 200 µM | One-way ANOVA (Sidak) | No | ns | 0.827745 |
|  | 50 µM vs. 400 µM | One-way ANOVA (Sidak) | No | ns | 0.999964 |
|  | 50 µM vs. 800 µM | One-way ANOVA (Sidak) | No | ns | 0.779358 |
|  | 100 µM vs. 200 µM | One-way ANOVA (Sidak) | No | ns | 0.712786 |
|  | 100 µM vs. 400 µM | One-way ANOVA (Sidak) | No | ns | 0.998999 |
|  | 100 µM vs. 800 µM | One-way ANOVA (Sidak) | No | ns | 0.879366 |
|  | 200 µM vs. 400 µM | One-way ANOVA (Sidak) | No | ns | 0.932727 |
|  | 200 µM vs. 800 µM | One-way ANOVA (Sidak) | No | ns | 0.104107 |
|  | 400 µM vs. 800 µM | One-way ANOVA (Sidak) | No | ns | 0.615584 |
| **Figure 5G (middle panel)** | 0 µM vs. 50 µM | One-way ANOVA (Tukey) | No | ns | 0.997131 |
|  | 0 µM vs. 100 µM | One-way ANOVA (Tukey) | Yes | ** | 0.001858 |
|  | 0 µM vs. 200 µM | One-way ANOVA (Tukey) | Yes | **** | <0.000001 |
|  | 50 µM vs. 100 µM | One-way ANOVA (Tukey) | Yes | **** | 0.000003 |
|  | 50 µM vs. 200 µM | One-way ANOVA (Tukey) | Yes | **** | <0.000001 |
|  | 100 µM vs. 200 µM | One-way ANOVA (Tukey) | Yes | **** | <0.000001 |
| **Figure 5G (right panel)** | Ctrl vs. 50 µM | One-way ANOVA (Tukey) | No | ns | 0.972100 |
|  | Ctrl vs. 100 µM | One-way ANOVA (Tukey) | No | ns | 0.072900 |
| **Figure 5H** | 0 µM vs. 25 µM | One-way ANOVA (Sidak) | No | ns | 0.999019 |
|  | 0 µM vs. 50 µM | One-way ANOVA (Sidak) | Yes | **** | 0.000001 |
|  | 0 µM vs. 100 µM | One-way ANOVA (Sidak) | Yes | **** | <0.000001 |
|  | 0 µM vs. 200 µM | One-way ANOVA (Sidak) | Yes | **** | <0.000001 |
|  | 0 µM vs. 400 µM | One-way ANOVA (Sidak) | Yes | **** | <0.000001 |
|  | 0 µM vs. 800 µM | One-way ANOVA (Sidak) | Yes | **** | <0.000001 |
|  | 25 µM vs. 50 µM | One-way ANOVA (Sidak) | Yes | **** | <0.000001 |
|  | 25 µM vs. 100 µM | One-way ANOVA (Sidak) | Yes | **** | <0.000001 |
|  | 25 µM vs. 200 µM | One-way ANOVA (Sidak) | Yes | **** | <0.000001 |
|  | 25 µM vs. 400 µM | One-way ANOVA (Sidak) | Yes | **** | <0.000001 |
|  | 25 µM vs. 800 µM | One-way ANOVA (Sidak) | Yes | **** | <0.000001 |
|  | 50 µM vs. 100 µM | One-way ANOVA (Sidak) | Yes | **** | <0.000001 |
|  | 50 µM vs. 200 µM | One-way ANOVA (Sidak) | Yes | **** | <0.000001 |
|  | 50 µM vs. 400 µM | One-way ANOVA (Sidak) | Yes | **** | <0.000001 |
|  | 50 µM vs. 800 µM | One-way ANOVA (Sidak) | Yes | **** | <0.000001 |
|  | 100 µM vs. 200 µM | One-way ANOVA (Sidak) | Yes | **** | <0.000001 |
|  | 100 µM vs. 400 µM | One-way ANOVA (Sidak) | Yes | **** | <0.000001 |
|  | 100 µM vs. 800 µM | One-way ANOVA (Sidak) | Yes | **** | <0.000001 |
|  | 200 µM vs. 400 µM | One-way ANOVA (Sidak) | Yes | **** | <0.000001 |
|  | 200 µM vs. 800 µM | One-way ANOVA (Sidak) | Yes | **** | <0.000001 |
|  | 400 µM vs. 800 µM | One-way ANOVA (Sidak) | Yes | *** | 0.000819 |
| **Figure 5I** | 0 µM vs. 50 µM | One-way ANOVA (Tukey) | Yes | * | 0.023370 |
|  | 0 µM vs. 100 µM | One-way ANOVA (Tukey) | Yes | *** | 0.000150 |
|  | 0 µM vs. 200 µM | One-way ANOVA (Tukey) | Yes | **** | <0.000001 |
|  | 50 µM vs. 100 µM | One-way ANOVA (Tukey) | Yes | **** | <0.000001 |
|  | 50 µM vs. 200 µM | One-way ANOVA (Tukey) | Yes | **** | <0.000001 |
|  | 100 µM vs. 200 µM | One-way ANOVA (Tukey) | Yes | *** | 0.000227 |
| **Figure 5J** | 0.025 | One-way ANOVA (Tukey) | No | ns | 0.904504 |
|  | 0.05 | One-way ANOVA (Tukey) | No | ns | 0.972360 |
|  | 0.1 | One-way ANOVA (Tukey) | Yes | ** | 0.001112 |
|  | 0.2 | One-way ANOVA (Tukey) | Yes | **** | 0.000003 |
|  | 0.4 | One-way ANOVA (Tukey) | Yes | **** | 0.000001 |
|  | 0.8 | One-way ANOVA (Tukey) | No | ns | 0.055010 |
| **Figure 6A** | 0 µM vs. 25 µM | One-way ANOVA (Sidak) | No | ns | 0.736371 |
|  | 0 µM vs. 50 µM | One-way ANOVA (Sidak) | Yes | *** | 0.000376 |
|  | 0 µM vs. 100 µM | One-way ANOVA (Sidak) | Yes | **** | <0.000001 |
|  | 0 µM vs. 200 µM | One-way ANOVA (Sidak) | Yes | **** | <0.000001 |
|  | 0 µM vs. 400 µM | One-way ANOVA (Sidak) | Yes | **** | <0.000001 |
|  | 0 µM vs. 800 µM | One-way ANOVA (Sidak) | Yes | **** | <0.000001 |
|  | 25 µM vs. 50 µM | One-way ANOVA (Sidak) | Yes | **** | 0.000003 |
|  | 25 µM vs. 100 µM | One-way ANOVA (Sidak) | Yes | **** | <0.000001 |
|  | 25 µM vs. 200 µM | One-way ANOVA (Sidak) | Yes | **** | <0.000001 |
|  | 25 µM vs. 400 µM | One-way ANOVA (Sidak) | Yes | **** | <0.000001 |
|  | 25 µM vs. 800 µM | One-way ANOVA (Sidak) | Yes | **** | <0.000001 |
|  | 50 µM vs. 100 µM | One-way ANOVA (Sidak) | Yes | **** | <0.000001 |
|  | 50 µM vs. 200 µM | One-way ANOVA (Sidak) | Yes | **** | <0.000001 |
|  | 50 µM vs. 400 µM | One-way ANOVA (Sidak) | Yes | **** | <0.000001 |
|  | 50 µM vs. 800 µM | One-way ANOVA (Sidak) | Yes | **** | <0.000001 |
|  | 100 µM vs. 200 µM | One-way ANOVA (Sidak) | Yes | * | 0.014955 |
|  | 100 µM vs. 400 µM | One-way ANOVA (Sidak) | Yes | **** | 0.000001 |
|  | 100 µM vs. 800 µM | One-way ANOVA (Sidak) | Yes | **** | <0.000001 |
|  | 200 µM vs. 400 µM | One-way ANOVA (Sidak) | No | ns | 0.060504 |
|  | 200 µM vs. 800 µM | One-way ANOVA (Sidak) | Yes | **** | <0.000001 |
|  | 400 µM vs. 800 µM | One-way ANOVA (Sidak) | Yes | **** | 0.000021 |
| **Figure 6B** | 0 µM vs. 50 µM | One-way ANOVA (Tukey) | No | ns | 0.088990 |
|  | 0 µM vs. 100 µM | One-way ANOVA (Tukey) | Yes | **** | <0.000001 |
|  | 0 µM vs. 200 µM | One-way ANOVA (Tukey) | Yes | **** | <0.000001 |
|  | 50 µM vs. 100 µM | One-way ANOVA (Tukey) | Yes | **** | <0.000001 |
|  | 50 µM vs. 200 µM | One-way ANOVA (Tukey) | Yes | **** | <0.000001 |
|  | 100 µM vs. 200 µM | One-way ANOVA (Tukey) | Yes | **** | <0.000001 |
| **Figure 6C** | 0.025 | One-way ANOVA (Tukey) | Yes | **** | 0.000007 |
|  | 0.05 | One-way ANOVA (Tukey) | No | ns | 0.999988 |
|  | 0.1 | One-way ANOVA (Tukey) | Yes | **** | <0.000001 |
|  | 0.2 | One-way ANOVA (Tukey) | Yes | **** | <0.000001 |
|  | 0.4 | One-way ANOVA (Tukey) | Yes | **** | <0.000001 |
|  | 0.8 | One-way ANOVA (Tukey) | No | ns | 0.999997 |
| **Figure 6D (middle panel)** | Ctrl vs. 50 µM | One-way ANOVA (Tukey) | No | ns | 0.898871 |
|  | Ctrl vs. 100 µM | One-way ANOVA (Tukey) | No | ns | 0.995732 |
|  | 50 µM vs. 100 µM | One-way ANOVA (Tukey) | No | ns | 0.857861 |
| **Figure 6D (right panel)** | Ctrl vs. 50 µM | One-way ANOVA (Dunn) | No | ns | >0.999999 |
|  | Ctrl vs. 100 µM | One-way ANOVA (Dunn) | No | ns | >0.999999 |
|  | 50 µM vs. 100 µM | One-way ANOVA (Dunn) | No | ns | >0.999999 |
| **Figure 6E** | 0 µM vs. 25 µM | One-way ANOVA (Sidak) | No | ns | 0.682499 |
|  | 0 µM vs. 50 µM | One-way ANOVA (Sidak) | Yes | * | 0.012770 |
|  | 0 µM vs. 100 µM | One-way ANOVA (Sidak) | Yes | **** | <0.000001 |
|  | 0 µM vs. 200 µM | One-way ANOVA (Sidak) | Yes | **** | <0.000001 |
|  | 0 µM vs. 400 µM | One-way ANOVA (Sidak) | Yes | **** | <0.000001 |
|  | 0 µM vs. 800 µM | One-way ANOVA (Sidak) | Yes | **** | <0.000001 |
|  | 25 µM vs. 50 µM | One-way ANOVA (Sidak) | No | ns | 0.417749 |
|  | 25 µM vs. 100 µM | One-way ANOVA (Sidak) | Yes | **** | <0.000001 |
|  | 25 µM vs. 200 µM | One-way ANOVA (Sidak) | Yes | **** | <0.000001 |
|  | 25 µM vs. 400 µM | One-way ANOVA (Sidak) | Yes | **** | <0.000001 |
|  | 25 µM vs. 800 µM | One-way ANOVA (Sidak) | Yes | **** | <0.000001 |
|  | 50 µM vs. 100 µM | One-way ANOVA (Sidak) | Yes | **** | 0.000055 |
|  | 50 µM vs. 200 µM | One-way ANOVA (Sidak) | Yes | **** | <0.000001 |
|  | 50 µM vs. 400 µM | One-way ANOVA (Sidak) | Yes | **** | <0.000001 |
|  | 50 µM vs. 800 µM | One-way ANOVA (Sidak) | Yes | **** | <0.000001 |
|  | 100 µM vs. 200 µM | One-way ANOVA (Sidak) | Yes | * | 0.029668 |
|  | 100 µM vs. 400 µM | One-way ANOVA (Sidak) | No | ns | 0.051992 |
|  | 100 µM vs. 800 µM | One-way ANOVA (Sidak) | Yes | * | 0.029503 |
|  | 200 µM vs. 400 µM | One-way ANOVA (Sidak) | No | ns | 0.999988 |
|  | 200 µM vs. 800 µM | One-way ANOVA (Sidak) | No | ns | >0.999999 |
|  | 400 µM vs. 800 µM | One-way ANOVA (Sidak) | No | ns | 0.999987 |
| **Figure 6G** | U87 T-f L vs. U87 L (0 µM ) | One-way ANOVA (Sidak) | No | ns | 0.944517 |
|  | U87 T-f L vs. U87 L (50 µM) | One-way ANOVA (Sidak) | No | ns | 0.790840 |
|  | U87 T-f L vs. U87 L (100 µM) | One-way ANOVA (Sidak) | No | ns | 0.998589 |
|  | U87 T-f L vs. U87 L (200 µM) | One-way ANOVA (Sidak) | No | ns | 0.999959 |
| **Figure 6H** | HMC3 T-f L vs. HMC3 L (0 µM ) | One-way ANOVA (Sidak) | No | ns | 0.854129 |
|  | HMC3 T-f L vs. HMC3 L (50 µM) | One-way ANOVA (Sidak) | No | ns | 0.052411 |
|  | HMC3 T-f L vs. HMC3 L (100 µM) | One-way ANOVA (Sidak) | Yes | **** | 0.000003 |
|  | HMC3 T-f L vs. HMC3 L (200 µM) | One-way ANOVA (Sidak) | Yes | **** | <0.000001 |
| **Figure 6I (middle panel) SN6 192** | 0 µM vs. 1.25 µM | One-way ANOVA (Sidak) | Yes | **** | <0.000001 |
|  | 0 µM vs. 2.5 µM | One-way ANOVA (Sidak) | Yes | **** | 0.000002 |
|  | 0 µM vs. 5 µM | One-way ANOVA (Sidak) | No | ns | 0.165091 |
|  | 0 µM vs. 10 µM | One-way ANOVA (Sidak) | No | ns | 0.999999 |
|  | 0 µM vs. 20 µM | One-way ANOVA (Sidak) | No | ns | 0.481589 |
|  | 0 µM vs. 50 µM | One-way ANOVA (Sidak) | Yes | **** | <0.000001 |
|  | 1.25 µM vs. 2.5 µM | One-way ANOVA (Sidak) | No | ns | 0.196888 |
|  | 1.25 µM vs. 5 µM | One-way ANOVA (Sidak) | Yes | **** | 0.000003 |
|  | 1.25 µM vs. 10 µM | One-way ANOVA (Sidak) | Yes | **** | <0.000001 |
|  | 1.25 µM vs. 20 µM | One-way ANOVA (Sidak) | Yes | **** | <0.000001 |
|  | 1.25 µM vs. 50 µM | One-way ANOVA (Sidak) | Yes | **** | <0.000001 |
|  | 2.5 µM vs. 5 µM | One-way ANOVA (Sidak) | Yes | ** | 0.006982 |
|  | 2.5 µM vs. 10 µM | One-way ANOVA (Sidak) | Yes | **** | 0.000001 |
|  | 2.5 µM vs. 20 µM | One-way ANOVA (Sidak) | Yes | **** | <0.000001 |
|  | 2.5 µM vs. 50 µM | One-way ANOVA (Sidak) | Yes | **** | <0.000001 |
|  | 5 µM vs. 10 µM | One-way ANOVA (Sidak) | No | ns | 0.119381 |
|  | 5 µM vs. 20 µM | One-way ANOVA (Sidak) | Yes | ** | 0.001051 |
|  | 5 µM vs. 50 µM | One-way ANOVA (Sidak) | Yes | **** | <0.000001 |
|  | 10 µM vs. 20 µM | One-way ANOVA (Sidak) | No | ns | 0.581710 |
|  | 10 µM vs. 50 µM | One-way ANOVA (Sidak) | Yes | **** | <0.000001 |
|  | 20 µM vs. 50 µM | One-way ANOVA (Sidak) | Yes | **** | <0.000001 |
| **Figure 6I (right panel)** | 0 µM vs. 0.25 µM | One-way ANOVA (Sidak) | No | ns | 0.321169 |
|  | 0 µM vs. 0.5 µM | One-way ANOVA (Sidak) | Yes | **** | 0.000014 |
|  | 0 µM vs. 1 µM | One-way ANOVA (Sidak) | Yes | **** | <0.000001 |
|  | 0 µM vs. 2 µM | One-way ANOVA (Sidak) | Yes | **** | <0.000001 |
|  | 0 µM vs. 4 µM | One-way ANOVA (Sidak) | Yes | **** | <0.000001 |
|  | 0 µM vs. 8 µM | One-way ANOVA (Sidak) | Yes | **** | <0.000001 |
|  | 0.25 µM vs. 0.5 µM | One-way ANOVA (Sidak) | Yes | * | 0.012013 |
|  | 0.25 µM vs. 1µM | One-way ANOVA (Sidak) | Yes | **** | <0.000001 |
|  | 0.25 µM vs. 2 µM | One-way ANOVA (Sidak) | Yes | **** | <0.000001 |
|  | 0.25 µM vs. 4 µM | One-way ANOVA (Sidak) | Yes | **** | <0.000001 |
|  | 0.25 µM vs. 8 µM | One-way ANOVA (Sidak) | Yes | **** | <0.000001 |
|  | 0.5 µM vs. 1µM | One-way ANOVA (Sidak) | Yes | **** | <0.000001 |
|  | 0.5 µM vs. 2 µM | One-way ANOVA (Sidak) | Yes | **** | <0.000001 |
|  | 0.5 µM vs. 4 µM | One-way ANOVA (Sidak) | Yes | **** | <0.000001 |
|  | 0.5 µM vs. 8 µM | One-way ANOVA (Sidak) | Yes | **** | <0.000001 |
|  | 1 µM vs. 2 µM | One-way ANOVA (Sidak) | Yes | **** | 0.000001 |
|  | 1 µM vs. 4 µM | One-way ANOVA (Sidak) | Yes | **** | <0.000001 |
|  | 1 µM vs. 8 µM | One-way ANOVA (Sidak) | Yes | **** | <0.000001 |
|  | 2 µM vs. 4 µM | One-way ANOVA (Sidak) | Yes | **** | <0.000001 |
|  | 2 µM vs. 8 µM | One-way ANOVA (Sidak) | Yes | **** | <0.000001 |
|  | 4µM vs. 8 µM | One-way ANOVA (Sidak) | No | ns | 0.736214 |
| **Figure 6J** | 0 µM vs. 1.25 µM | One-way ANOVA (Sidak) | No | ns | 0.448682 |
|  | 0 µM vs. 2.5 µM | One-way ANOVA (Sidak) | Yes | **** | 0.000095 |
|  | 0 µM vs. 5 µM | One-way ANOVA (Sidak) | Yes | **** | <0.000001 |
|  | 0 µM vs. 10 µM | One-way ANOVA (Sidak) | Yes | **** | <0.000001 |
|  | 0 µM vs. 20 µM | One-way ANOVA (Sidak) | Yes | **** | <0.000001 |
|  | 0 µM vs. 50 µM | One-way ANOVA (Sidak) | Yes | **** | <0.000001 |
|  | 1.25 µM vs. 2.5 µM | One-way ANOVA (Sidak) | Yes | * | 0.032573 |
|  | 1.25 µM vs. 5 µM | One-way ANOVA (Sidak) | Yes | **** | <0.000001 |
|  | 1.25 µM vs. 10 µM | One-way ANOVA (Sidak) | Yes | **** | <0.000001 |
|  | 1.25 µM vs. 20 µM | One-way ANOVA (Sidak) | Yes | **** | <0.000001 |
|  | 1.25 µM vs. 50 µM | One-way ANOVA (Sidak) | Yes | **** | <0.000001 |
|  | 2.5 µM vs. 5 µM | One-way ANOVA (Sidak) | Yes | **** | <0.000001 |
|  | 2.5 µM vs. 10 µM | One-way ANOVA (Sidak) | Yes | **** | <0.000001 |
|  | 2.5 µM vs. 20 µM | One-way ANOVA (Sidak) | Yes | **** | <0.000001 |
|  | 2.5 µM vs. 50 µM | One-way ANOVA (Sidak) | Yes | **** | <0.000001 |
|  | 5 µM vs. 10 µM | One-way ANOVA (Sidak) | No | ns | 0.380101 |
|  | 5 µM vs. 20 µM | One-way ANOVA (Sidak) | Yes | ** | 0.002962 |
|  | 5 µM vs. 50 µM | One-way ANOVA (Sidak) | Yes | **** | <0.000001 |
|  | 10 µM vs. 20 µM | One-way ANOVA (Sidak) | No | ns | 0.401834 |
|  | 10 µM vs. 50 µM | One-way ANOVA (Sidak) | Yes | **** | 0.000033 |
|  | 20 µM vs. 50 µM | One-way ANOVA (Sidak) | Yes | * | 0.017130 |
| **Suppl. Fig. 2** | LRRC8A |  |  |  |  |
|  | S557 vs. S565 | One-way ANOVA (Sidak) | No | ns | 0.951657 |
|  | S557 vs. S566 | One-way ANOVA (Sidak) | Yes | **** | <0.000001 |
|  | S557 vs. S610 | One-way ANOVA (Sidak) | Yes | **** | <0.000001 |
|  | S557 vs. S567 | One-way ANOVA (Sidak) | Yes | **** | <0.000001 |
|  | S557 vs. S568 | One-way ANOVA (Sidak) | Yes | **** | <0.000001 |
|  | S565 vs. S566 | One-way ANOVA (Sidak) | Yes | **** | <0.000001 |
|  | S565 vs. S610 | One-way ANOVA (Sidak) | Yes | **** | <0.000001 |
|  | S565 vs. S567 | One-way ANOVA (Sidak) | Yes | **** | <0.000001 |
|  | S565 vs. S568 | One-way ANOVA (Sidak) | Yes | **** | <0.000001 |
|  | S566 vs. S610 | One-way ANOVA (Sidak) | Yes | **** | <0.000001 |
|  | S566 vs. S567 | One-way ANOVA (Sidak) | No | ns | 0.161094 |
|  | S566 vs. S568 | One-way ANOVA (Sidak) | Yes | **** | <0.000001 |
|  | S610 vs. S567 | One-way ANOVA (Sidak) | Yes | **** | <0.000001 |
|  | S610 vs. S568 | One-way ANOVA (Sidak) | Yes | **** | <0.000001 |
|  | S567 vs. S568 | One-way ANOVA (Sidak) | Yes | **** | <0.000001 |
|  | LRRC8B |  |  |  |  |
|  | S557 vs. S565 | One-way ANOVA (Sidak) | Yes | **** | <0.000001 |
|  | S557 vs. S566 | One-way ANOVA (Sidak) | No | ns | 0.157404 |
|  | S557 vs. S610 | One-way ANOVA (Sidak) | Yes | **** | <0.000001 |
|  | S557 vs. S567 | One-way ANOVA (Sidak) | Yes | **** | <0.000001 |
|  | S557 vs. S568 | One-way ANOVA (Sidak) | Yes | **** | <0.000001 |
|  | S565 vs. S566 | One-way ANOVA (Sidak) | Yes | **** | <0.000001 |
|  | S565 vs. S610 | One-way ANOVA (Sidak) | Yes | * | 0.283520 |
|  | S565 vs. S567 | One-way ANOVA (Sidak) | No | ns | 559.265.000000 |
|  | S565 vs. S568 | One-way ANOVA (Sidak) | Yes | **** | <0.000001 |
|  | S566 vs. S610 | One-way ANOVA (Sidak) | Yes | **** | <0.000001 |
|  | S566 vs. S567 | One-way ANOVA (Sidak) | Yes | **** | 0.000010 |
|  | S566 vs. S568 | One-way ANOVA (Sidak) | Yes | **** | <0.000001 |
|  | S610 vs. S567 | One-way ANOVA (Sidak) | Yes | *** | 0.001400 |
|  | S610 vs. S568 | One-way ANOVA (Sidak) | Yes | **** | <0.000001 |
|  | S567 vs. S568 | One-way ANOVA (Sidak) | Yes | **** | <0.000001 |
|  | LRRC8C |  |  |  |  |
|  | S557 vs. S565 | One-way ANOVA (Sidak) | Yes | **** | 0.000040 |
|  | S557 vs. S566 | One-way ANOVA (Sidak) | No | ns | 0.997458 |
|  | S557 vs. S610 | One-way ANOVA (Sidak) | Yes | * | 0.012180 |
|  | S557 vs. S567 | One-way ANOVA (Sidak) | No | ns | 0.642381 |
|  | S557 vs. S568 | One-way ANOVA (Sidak) | Yes | **** | <0.000001 |
|  | S565 vs. S566 | One-way ANOVA (Sidak) | Yes | **** | 0.000220 |
|  | S565 vs. S610 | One-way ANOVA (Sidak) | No | ns | 0.197486 |
|  | S565 vs. S567 | One-way ANOVA (Sidak) | Yes | ** | 0.001001 |
|  | S565 vs. S568 | One-way ANOVA (Sidak) | No | ns | 0.997218 |
|  | S566 vs. S610 | One-way ANOVA (Sidak) | Yes | * | 0.042444 |
|  | S566 vs. S567 | One-way ANOVA (Sidak) | No | ns | 0.886413 |
|  | S566 vs. S568 | One-way ANOVA (Sidak) | Yes | **** | 0.000004 |
|  | S610 vs. S567 | One-way ANOVA (Sidak) | No | ns | 0.397175 |
|  | S610 vs. S568 | One-way ANOVA (Sidak) | No | ns | 0.720740 |
|  | S567 vs. S568 | One-way ANOVA (Sidak) | Yes | *** | 0.000204 |
|  | LRRC8D |  |  |  |  |
|  | S557 vs. S565 | One-way ANOVA (Sidak) | Yes | **** | <0.000001 |
|  | S557 vs. S566 | One-way ANOVA (Sidak) | Yes | **** | <0.000001 |
|  | S557 vs. S610 | One-way ANOVA (Sidak) | Yes | **** | <0.000001 |
|  | S557 vs. S567 | One-way ANOVA (Sidak) | Yes | **** | <0.000001 |
|  | S557 vs. S568 | One-way ANOVA (Sidak) | Yes | **** | <0.000001 |
|  | S565 vs. S566 | One-way ANOVA (Sidak) | Yes | *** | 0.000175 |
|  | S565 vs. S610 | One-way ANOVA (Sidak) | Yes | **** | 0.000075 |
|  | S565 vs. S567 | One-way ANOVA (Sidak) | No | ns | 0.326078 |
|  | S565 vs. S568 | One-way ANOVA (Sidak) | Yes | **** | <0.000001 |
|  | S566 vs. S610 | One-way ANOVA (Sidak) | Yes | **** | <0.000001 |
|  | S566 vs. S567 | One-way ANOVA (Sidak) | Yes | **** | <0.000001 |
|  | S566 vs. S568 | One-way ANOVA (Sidak) | Yes | **** | <0.000001 |
|  | S610 vs. S567 | One-way ANOVA (Sidak) | Yes | * | 0.494300 |
|  | S610 vs. S568 | One-way ANOVA (Sidak) | Yes | **** | <0.000001 |
|  | S567 vs. S568 | One-way ANOVA (Sidak) | Yes | **** | <0.000001 |
|  | LRRC8E |  |  |  |  |
|  | S557 vs. S565 | One-way ANOVA (Sidak) | No | ns | 0.999906 |
|  | S557 vs. S566 | One-way ANOVA (Sidak) | No | ns | 0.999983 |
|  | S557 vs. S610 | One-way ANOVA (Sidak) | No | ns | 0.999478 |
|  | S557 vs. S567 | One-way ANOVA (Sidak) | No | ns | 0.854385 |
|  | S557 vs. S568 | One-way ANOVA (Sidak) | No | ns | 0.999960 |
|  | S565 vs. S566 | One-way ANOVA (Sidak) | No | ns | >0.999999 |
|  | S565 vs. S610 | One-way ANOVA (Sidak) | No | ns | 0.999999 |
|  | S565 vs. S567 | One-way ANOVA (Sidak) | No | ns | 0.937743 |
|  | S565 vs. S568 | One-way ANOVA (Sidak) | No | ns | >0.999999 |
|  | S566 vs. S610 | One-way ANOVA (Sidak) | No | ns | 0.999983 |
|  | S566 vs. S567 | One-way ANOVA (Sidak) | No | ns | 0.917759 |
|  | S566 vs. S568 | One-way ANOVA (Sidak) | No | ns | >0.999999 |
|  | S610 vs. S567 | One-way ANOVA (Sidak) | No | ns | 0.960431 |
|  | S610 vs. S568 | One-way ANOVA (Sidak) | No | ns | 0.999994 |
|  | S567 vs. S568 | One-way ANOVA (Sidak) | No | ns | 0.927242 |
| **Suppl. Fig. 11A** | 0 µM vs. 0.25 µM | One-way ANOVA (Sidak) | No | ns | 0.999983 |
|  | 0 µM vs. 0.5 µM | One-way ANOVA (Sidak) | No | ns | 0.998190 |
|  | 0 µM vs. 1 µM | One-way ANOVA (Sidak) | No | ns | 0.400678 |
|  | 0 µM vs. 2 µM | One-way ANOVA (Sidak) | No | ns | 0.052682 |
|  | 0 µM vs. 4 µM | One-way ANOVA (Sidak) | Yes | **** | 0.000004 |
|  | 0 µM vs. 8 µM | One-way ANOVA (Sidak) | Yes | **** | <0.000001 |
|  | 0.25 µM vs. 0.5 µM | One-way ANOVA (Sidak) | No | ns | 0.986597 |
|  | 0.25 µM vs. 1µM | One-way ANOVA (Sidak) | No | ns | 0.274032 |
|  | 0.25 µM vs. 2 µM | One-way ANOVA (Sidak) | Yes | * | 0.029060 |
|  | 0.25 µM vs. 4 µM | One-way ANOVA (Sidak) | Yes | **** | 0.000002 |
|  | 0.25 µM vs. 8 µM | One-way ANOVA (Sidak) | Yes | **** | <0.000001 |
|  | 0.5 µM vs. 1µM | One-way ANOVA (Sidak) | No | ns | 0.733244 |
|  | 0.5 µM vs. 2 µM | One-way ANOVA (Sidak) | No | ns | 0.170779 |
|  | 0.5 µM vs. 4 µM | One-way ANOVA (Sidak) | Yes | **** | 0.000020 |
|  | 0.5 µM vs. 8 µM | One-way ANOVA (Sidak) | Yes | **** | <0.000001 |
|  | 1 µM vs. 2 µM | One-way ANOVA (Sidak) | No | ns | 0.945264 |
|  | 1 µM vs. 4 µM | One-way ANOVA (Sidak) | Yes | ** | 0.002441 |
|  | 1 µM vs. 8 µM | One-way ANOVA (Sidak) | Yes | **** | 0.000017 |
|  | 2 µM vs. 4 µM | One-way ANOVA (Sidak) | Yes | * | 0.041444 |
|  | 2 µM vs. 8 µM | One-way ANOVA (Sidak) | Yes | *** | 0.000461 |
|  | 4µM vs. 8 µM | One-way ANOVA (Sidak) | No | ns | 0.702177 |
| **Suppl. Fig. 11B** | 0 µM vs. 0.25 µM | One-way ANOVA (Sidak) | No | ns | 0.158010 |
|  | 0 µM vs. 0.5 µM | One-way ANOVA (Sidak) | Yes | **** | <0.000001 |
|  | 0 µM vs. 1 µM | One-way ANOVA (Sidak) | Yes | **** | <0.000001 |
|  | 0 µM vs. 2 µM | One-way ANOVA (Sidak) | Yes | **** | <0.000001 |
|  | 0 µM vs. 4 µM | One-way ANOVA (Sidak) | Yes | **** | <0.000001 |
|  | 0 µM vs. 8 µM | One-way ANOVA (Sidak) | Yes | **** | <0.000001 |
|  | 0.25 µM vs. 0.5 µM | One-way ANOVA (Sidak) | Yes | **** | <0.000001 |
|  | 0.25 µM vs. 1µM | One-way ANOVA (Sidak) | Yes | **** | <0.000001 |
|  | 0.25 µM vs. 2 µM | One-way ANOVA (Sidak) | Yes | **** | <0.000001 |
|  | 0.25 µM vs. 4 µM | One-way ANOVA (Sidak) | Yes | **** | <0.000001 |
|  | 0.25 µM vs. 8 µM | One-way ANOVA (Sidak) | Yes | **** | <0.000001 |
|  | 0.5 µM vs. 1µM | One-way ANOVA (Sidak) | Yes | **** | <0.000001 |
|  | 0.5 µM vs. 2 µM | One-way ANOVA (Sidak) | Yes | **** | <0.000001 |
|  | 0.5 µM vs. 4 µM | One-way ANOVA (Sidak) | Yes | **** | <0.000001 |
|  | 0.5 µM vs. 8 µM | One-way ANOVA (Sidak) | Yes | **** | <0.000001 |
|  | 1 µM vs. 2 µM | One-way ANOVA (Sidak) | Yes | *** | 0.000345 |
|  | 1 µM vs. 4 µM | One-way ANOVA (Sidak) | Yes | **** | <0.000001 |
|  | 1 µM vs. 8 µM | One-way ANOVA (Sidak) | Yes | **** | <0.000001 |
|  | 2 µM vs. 4 µM | One-way ANOVA (Sidak) | Yes | *** | 0.000702 |
|  | 2 µM vs. 8 µM | One-way ANOVA (Sidak) | Yes | **** | <0.000001 |
|  | 4µM vs. 8 µM | One-way ANOVA (Sidak) | Yes | *** | 0.000369 |
| **Suppl. Fig. 11C** | 0 µM vs. 0.25 µM | One-way ANOVA (Sidak) | No | ns | 0.999854 |
|  | 0 µM vs. 0.5 µM | One-way ANOVA (Sidak) | No | ns | >0.999999 |
|  | 0 µM vs. 1 µM | One-way ANOVA (Sidak) | No | ns | >0.999999 |
|  | 0 µM vs. 2 µM | One-way ANOVA (Sidak) | No | ns | 0.997477 |
|  | 0 µM vs. 4 µM | One-way ANOVA (Sidak) | Yes | **** | <0.000001 |
|  | 0 µM vs. 8 µM | One-way ANOVA (Sidak) | Yes | **** | <0.000001 |
|  | 0.25 µM vs. 0.5 µM | One-way ANOVA (Sidak) | No | ns | 0.999854 |
|  | 0.25 µM vs. 1µM | One-way ANOVA (Sidak) | No | ns | 0.999927 |
|  | 0.25 µM vs. 2 µM | One-way ANOVA (Sidak) | No | ns | 0.969353 |
|  | 0.25 µM vs. 4 µM | One-way ANOVA (Sidak) | Yes | **** | <0.000001 |
|  | 0.25 µM vs. 8 µM | One-way ANOVA (Sidak) | Yes | **** | <0.000001 |
|  | 0.5 µM vs. 1µM | One-way ANOVA (Sidak) | No | ns | >0.999999 |
|  | 0.5 µM vs. 2 µM | One-way ANOVA (Sidak) | No | ns | 0.997480 |
|  | 0.5 µM vs. 4 µM | One-way ANOVA (Sidak) | Yes | **** | <0.000001 |
|  | 0.5 µM vs. 8 µM | One-way ANOVA (Sidak) | Yes | **** | <0.000001 |
|  | 1 µM vs. 2 µM | One-way ANOVA (Sidak) | No | ns | 0.996377 |
|  | 1 µM vs. 4 µM | One-way ANOVA (Sidak) | Yes | **** | <0.000001 |
|  | 1 µM vs. 8 µM | One-way ANOVA (Sidak) | Yes | **** | <0.000001 |
|  | 2 µM vs. 4 µM | One-way ANOVA (Sidak) | Yes | **** | <0.000001 |
|  | 2 µM vs. 8 µM | One-way ANOVA (Sidak) | Yes | **** | <0.000001 |
|  | 4µM vs. 8 µM | One-way ANOVA (Sidak) | Yes | **** | 0.000040 |

We subsequently evaluated the biomechanical response of S82 cells to the cocktail. Treatment with 100 and 200 µM reduced the apparent Young’s modulus from 2.03 ± 0.03 kPa to 1.00 ± 0.04 kPa and to 0.13 ± 0.03 kPa, respectively, indicating a pronounced reduction in cellular stiffness (Fig. 6B). This response was consistent with the biomechanical changes observed in U87 cells, suggesting that the cocktail-induced softening extends to patient-derived GSCs. Finally, we applied the Bliss independence model to evaluate the cytotoxic combined pharmacological effect in S82 cells. The cocktail exhibited greater-than-expected effects within the same concentration range identified in U87 cells, with an even more pronounced effect in S82 cells (Fig. 6C). These findings indicate that the enhanced response to the combined treatment is not restricted to the established U87 cell line and can also be observed in a patient-derived GSC model.

Together, these findings show that the cocktail affects GSC viability and biomechanics, with heterogeneous patient responses. The association between VRAC-like currents and treatment sensitivity suggests that electrophysiological profiling may help identify responsive GSC populations.

### Multi-target strategy cocktail has a specific effect on GSCs

Standard glioma therapies can also affect non-malignant neural tissue. For example, temozolomide has been associated with mitochondrial dysfunction and morphological alterations in hippocampal neurons (39), whereas radiotherapy has been linked to cognitive impairment in patients (40). Given these potential adverse effects, we investigated whether the proposed multi-target pharmacological cocktail altered neuronal activity in primary mouse hippocampal cultures. To this end, we performed calcium imaging experiments following a pre-treatment for 72 hours with cocktail concentrations ranging from 50 to 100 µM (Fig. 6D). We first quantified the amount of spontaneous calcium transients occurring during 10 min of recording as a functional readout of neuronal activity. At 100 µM, the cocktail-treated neurons exhibited a mean calcium-spike of 41.5 ± 16.84 events, compared with 43.64 ± 16.88 events in vehicle-treated controls (Fig. 6D, middle panel). We then quantified the proportion of neurons exhibiting spontaneous calcium activity. Under treatment with 100 µM of the cocktail, 70 ± 0.08 % of neurons displayed spontaneous calcium spiking, with no statistically significant difference compared with vehicle-treated controls (Fig. 6D, Right panel). These findings indicate that, under the conditions tested, cocktail treatment did not significantly alter the proportion of spontaneously active neurons.

To further assess the selectivity of the proposed pharmacological strategy, we investigated its effects on the HMC3 cell line as a immortalized model of immune cells within the central nervous system. These cells were exposed to the cocktail at concentrations comparable to those used in GSC experiments. Notably, HMC3 cells exhibited substantial sensitivity to the treatment, with a cocktail concentration of 100 µM reducing cell viability to below 50% after 72 h of exposure (Fig. 6E). This finding highlighted the need to improve the selectivity of drug delivery to minimize potential off-target effects on non-malignant neural cells. We therefore explored liposomal encapsulation combined with receptor-mediated targeting as a strategy to improve the delivery of the cocktail to glioma cells.

As the cocktail comprises compounds with different physicochemical properties, we first evaluated two liposomal formulations to identify a suitable carrier for drug encapsulation. Conventional liposomes were composed of dipalmitoylphosphatidylcholine (DPPC, 60%), cholesterol (30%), and dipalmitoylphosphatidylglycerol (DPPG, 10%). A second formulation, referred to as PEGylated liposomes, contained DPPC (50%), cholesterol (30%), DPPG (10%), and DSPE-PEG2000 -COOH (10%), with the latter incorporated to provide a polyethylene glycol (PEG) coating. Empty and cocktail-loaded formulations were characterized using dynamic light scattering (DLS), nanoparticle tracking analysis (NTA) and atomic force microscopy (AFM) to assess their physicochemical properties, including particle size, concentration, morphology, and colloidal stability. The characterization revealed no significant differences in particle size or morphology between conventional and PEGylated liposomes, irrespective of cocktail loading. Furthermore, empty liposomes at a concentration of 20 µg/mL did not exert detectable cytotoxic effects under the conditions tested (Supplementary Fig. 9), supporting their suitability for subsequent biological evaluation.

We next quantified drug encapsulation efficiency following ultracentrifugation to separate liposome-associated compounds from unencapsulated drugs. The liposomes were subsequently disrupted to release their encapsulated contents, which were quantified by spectrophotometry. Across the tested conditions, the measured encapsulation efficiencies ranged from 75% to 98% (Fig. 6F), indicating efficient incorporation of the cocktail into the liposomal formulations.

As a molecular target for liposome functionalization, we select the transferrin receptor (TfR). which have been described as a target for glioma-directed delivery including the U87 cell model (41). The resulting transferrin-functionalized liposomes were subsequently characterized and exhibited physicochemical properties comparable to those of the corresponding non-functionalized formulations (Supplementary Fig. 9).

We evaluated the biological effects of cocktail-loaded non-functionalized and transferrin-functionalized liposomes (Fig. 6 G–H) in U87 glioblastoma cells and HMC3 cells. In U87 cells, both liposome formulations significantly decreased cell viability across the tested concentration range (50–200 µM; Fig. 6G), indicating that liposomal encapsulation preserved the cytotoxic activity of the cocktail against glioblastoma cells, consistent with the results shown in Fig. 5J. In HMC3 cells, transferrin-functionalized liposomes significantly reduced cytotoxicity compared to non-functionalized liposomes across the same concentration range (50–200 µM; Fig. 6H). These findings suggest that transferrin-functionalized liposomal delivery preserves the antitumor activity of the cocktail while attenuating its toxicity toward HMC3 cells, thereby enhancing the *in vitro* selectivity of the treatment.

These findings support combining multitarget pharmacology with transferrin-functionalized liposomes to improve drug-delivery selectivity and potentially reduce off-target toxicity in non-malignant neural cells.

### Na^+^/Ca^2+^ exchanger as an alternative therapy for unresponsive GSCs

Given that the multitarget cocktail elicited a significant response in only a subset of patient-derived GSC cultures, we investigated whether the NCX, a second electrophysiological target identified in these cells, could represent an alternative pharmacological target. We selected the patient-derived GSC culture S192, which exhibited an inward current sensitive to the NCX inhibitor SN-6 (Supplementary Fig. 10A). We then evaluated the effects of SN-6 and doxorubicin on cell viability. Doxorubicin, an FDA-approved chemotherapeutic agent, was included because it partially inhibited the NCX-associated current at the concentration used in our experiments (Fig. 3F).

SN-6 exerted a concentration-dependent cytotoxic effect on S192-derived GSCs across the tested concentration range of 0–50 µM (Fig. 6I, left panel). Doxorubicin also reduced cell viability at concentrations up to 8 µM (Fig. 6I, right panel); however, its effects were markedly variable. In a subsequent independent experiment, S192-derived GSCs displayed a substantially different sensitivity profile compared with the initial assay (Supplementary Fig. 11A).

To further investigate whether doxorubicin sensitivity was associated with the electrophysiological profile of the cells, we evaluated its effects on GSC cultures derived from patients S82 and S197 (Supplementary Fig. 8 and 10B), characterized by prominent VRAC-like and NCX-associated currents, respectively. Unexpectedly, S82-derived GSCs were more sensitive to doxorubicin at lower concentrations than S197-derived GSCs (Supplementary Fig. 11B-C). Thus, the observed cytotoxic response did not correspond directly to the electrophysiological profile of the cultures.

We also assessed the effects of SN-6 on HMC3 cell viability, SN-6 reduced its viability to below 50% at concentrations up to 10 µM (Fig. 6J), indicating that its cytotoxic effects were not restricted to GSCs.

Together, these findings identify NCX-associated activity as a potential vulnerability in patient-derived GSCs. However, SN-6 toxicity in HMC3 cells raises concerns about selectivity, while the variable effects of doxorubicin leave the relationship between NCX inhibition and cytotoxicity unclear. Further studies with more selective NCX inhibitors are needed to clarify its role and assess the value of electrophysiological profiling in identifying patient-specific pharmacological vulnerabilities.

## Discussion

In this study, we characterize the electrophysiological profiling of patient-derived GSCs identifies heterogeneous calcium-dependent ionic currents as candidate therapeutic targets, providing a rationale for developing multitarget strategies against glioblastoma.

### Electrophysiological screening of calcium-dependent currents on GSCs

The first candidate current identified in patient-derived GSCs was a calcium-dependent anion conductance with properties consistent with VRAC. Its amplitude increased with intracellular Ca²⁺ concentration, consistent with previous studies reporting Ca²⁺-dependent regulation of LRRC8-containing channels in heterologous expression systems (42–44). Furthermore, the current exhibited kinetic, permeability, and pharmacological properties consistent with those previously described for VRACs (45, 46). Taken together, these findings support the functional identification of a VRAC-like conductance in the GSCs examined in this study.

Interestingly, we also observed a strong correlation between LRRC8C expression, one of the variable subunits of the LRRC8 family, and the mean maximal current amplitude across patient-derived cultures. This association raises the possibility that LRRC8C contributes to the functional properties of the current in these cells, although a causal relationship cannot be established from the present data. Functional VRACs are heteromeric complexes comprising the essential LRRC8A subunit in combination with one or more of the other LRRC8 family members (LRRC8B–E). The composition of these complexes can influence their biophysical properties and substrate permeability (47,48). Furthermore, electrophysiological and pharmacological characterization alone cannot establish the molecular identity or subunit composition of the current recorded in our cells. Further molecular and functional studies, including targeted manipulation of individual LRRC8 subunits, will therefore be required to determine whether LRRC8C contributes to the observed current and to clarify the molecular composition of the VRAC-like conductance in patient-derived GSCs.

Notably, we did not detect currents consistent with other calcium-activated chloride conductances previously implicated in GBM, including those associated with TMEM16A and TMEM16B (49, 50). his finding contrasts with previous reports describing calcium-activated chloride currents in glioma models and suggests that the contribution of these conductances may vary according to cellular context. One possible explanation is the cellular heterogeneity of GBM, in which GSCs represent only one of several tumor-associated cell populations. For example, glioma-associated stem-like cell populations, including those described as glioma-associated stem cells (GASCs), have been reported to express a TMEM-like current sensitive to niflumic acid (NFA) (8). Differences in the cellular populations investigated could therefore contribute to the distinct electrophysiological profiles reported across studies (51).

An additional consideration concerns the calcium sensitivity of TMEM-family channels. Where some members requiring relatively high calcium levels to produce substantial currents (52, 53). Consequently, the absence of detectable TMEM16A/B-like currents in our recordings could also reflect differences in intracellular calcium conditions, channel expression, or experimental configuration. Thus, our findings do not exclude the presence of these channels in GSCs but instead suggest that the VRAC-like conductance was the predominant calcium-dependent anion current detected under the experimental conditions used in this study.

The second candidate, NCX-associated current, exhibited biophysical and pharmacological properties consistent with those reported for this transporter. NCX activity is electrogenic, and both the direction and amplitude of the resulting current depend on the transmembrane Na⁺ and Ca²⁺ gradients, as well as on membrane potential (54). In our recordings, the inward current is observed with 0.5 µM Ca²⁺ in the patch pipette and was inhibited by 5 mM Ni²⁺, a classical but non-selective NCX inhibitor, and by 20 µM SN-6, a pharmacological inhibitor of NCX (55). Although SN-6 preferentially inhibits the outward component of NCX current, this component was relatively small under our experimental conditions but remained sensitive to the inhibitor. These findings support the functional presence of an NCX-associated conductance in the patient-derived GSCs examined in this study. This observation is consistent with previous reports of NCX expression and function in glioma models, including the U251 cell line (26, 55).

### From electrophysiological target identification to a multitarget strategy

Electrophysiological screening provides the starting point for identifying therapeutic vulnerabilities, which then informs the development of a multitarget pharmacological cocktail designed to interfere with complementary mechanisms involved in glioma cell function (10, 56, 57). Rather than restricting the intervention to a single ion channel or transporter, we investigated whether combining compounds with distinct pharmacological activities could produce a more pronounced effect on tumor-cell viability and functional properties. Accordingly, the FDA approved drugs: niflumic acid (NFA), bumetanide and R-ketorolac were selected to target different, potentially complementary mechanisms: chloride conductances (57), cation-chloride cotransport (58, 59) and cytoskeleton-associated signaling (18, 60), respectively. This approach was intended to broaden the range of cellular processes affected by treatment and potentially overcome the limitations associated with targeting a single mechanism in a heterogeneous tumor population.

Nevertheless, the compounds included in the cocktail are not selective for their intended targets and may exert additional pharmacological effects.

In particular, niflumic acid (NFA) and R-Ketorolac (R-K) have been reported to inhibit cyclooxygenase (COX) enzymes at micromolar concentrations (61), although the R-enantiomer of ketorolac is largely devoid of COX-inhibitory activity and primarily targets Rho-family GTPases (62). However, the cytotoxic effects observed in GSCs required concentrations in the high micromolar range, which overlap with concentrations reported to inhibit VRAC-mediated currents and TMEM16A-associated chloride currents (52, 63, 64).

This concentration overlap is consistent with the possibility that modulation of chloride conductances contributes to the observed effects. Nevertheless, given the pharmacological profiles of these compounds, contributions from other molecular targets cannot be excluded, and the relative involvement of each mechanism remains to be established.

The multitarget cocktail reduced GSC viability more effectively than individual compounds and decreased the apparent Young’s modulus. Bliss independence analysis also indicated synergistic effects at selected concentrations. These findings support combining compounds that modulate complementary processes, potentially affecting ion homeostasis, cell-volume regulation, and cytoskeleton-associated signaling. However, the mechanisms underlying this interaction remain to be established through further studies of target engagement and the contribution of individual compounds.

Regarding the NCX-associated current, its partial inhibition by doxorubicin highlights the potential for pharmacological effects beyond those directly related to the exchanger. Doxorubicin exerts cytotoxic effects through multiple mechanisms, including DNA damage, which may help explain the lack of a clear association between the electrophysiological profiles of patient-derived GSC cultures and their sensitivity to this drug. Moreover, the role of NCX in glioma biology remains unclear, with studies reporting different outcomes depending on the experimental context and pharmacological approach. For example, NCX inhibition has been associated with impaired glioma cell migration without a significant reduction in cell viability (65), whereas other studies have reported cytotoxic effects following pharmacological inhibition of the exchanger’s forward mode (26). In our study, SN-6 reduced the viability of patient-derived GSCs at concentrations between 10 and 50 µM, suggesting that pharmacological inhibition of NCX-associated activity may be associated with cytotoxic effects in these cells. Nevertheless, given the concentration range used and the potential for off-target effects, the contribution of NCX inhibition to the observed cytotoxicity remains to be confirmed. Furthermore, studies in cardiomyocytes have highlighted a role for reverse-mode NCX activity in regulating intracellular ion homeostasis and cell-volume changes under specific conditions (66–68).

Whether similar mechanisms contribute to volume regulation and survival in GSCs remain unknown. Further studies using more selective pharmacological tools and approaches capable of distinguishing the forward and reverse modes of NCX are therefore needed to clarify its contribution to GSC physiology and assess its potential as a therapeutic target.

### Specificity of the therapy

Our results showed that cocktail treatment did not measurably alter the spontaneous activity of cultured hippocampal neurons, suggesting that neuronal activity was preserved under the experimental conditions tested. This finding is encouraging, particularly given the functional expression of calcium-activated chloride conductances in several brain regions (69–71) and the involvement of Rac1 signaling in neuronal functions such as synaptic plasticity (72). Nevertheless, the absence of detectable changes in spontaneous neuronal activity does not exclude other potential neurotoxic effects or adverse effects on additional cell types within the central nervous system.

In this regard, our finding that the cocktail exerted substantial cytotoxic effects on a HMC3 cell line highlights a potential limitation of the free-drug formulation. Importantly, encapsulation of the cocktail in transferrin-functionalized liposomes attenuated its cytotoxic effects on HMC3 cells while preserving its activity against glioblastoma cells. These findings support the potential of targeted drug-delivery strategies to improve the in vitro selectivity of the cocktail and reduce its impact on non-malignant cell populations. Although the individual compounds have established clinical uses, with characterized safety profiles in their respective therapeutic contexts (73–77), these profiles cannot be directly extrapolated to their combined administration. The safety and tolerability of this specific three-drug combination therefore remain to be determined.

Overall, although cultured hippocampal neurons and HMC3 cell lines provide useful in vitro models for evaluating potential off-target effects, further studies in appropriate animal models will be required to assess the systemic and neurological safety, tolerability, biodistribution, and therapeutic efficacy of the cocktail, both in its free form and following encapsulation in transferrin-functionalized liposomes.

## Materials and Methods

### Cell culture

Human U87-MG glioblastoma cells (RRID: CVCL_0022) and HMC3 microglial cells were obtained from ATCC and maintained in DMEM-HG and MEM, respectively, supplemented with 10% FBS, L-glutamine, penicillin, and streptomycin. Both cell lines were cultured at 37°C in a humidified atmosphere containing 5% CO₂ and routinely tested negative for mycoplasma contamination.

Patient-derived glioma stem cells (GSCs) were isolated from human glioblastoma (GBM) surgical specimens collected by the Department of Neurosurgery of the Azienda Ospedaliero-Universitaria di Udine and, where applicable, the Padova University-Hospital, Italy. Following previously described protocols (78, 79). Human tissue collection and use were approved by the relevant ethics committees, including the Independent Ethics Committee of the University Hospital of Udine (approval no. 196/2014Em) and the Padova University-Hospital (2462P), and conducted in accordance with the Declaration of Helsinki. Informed consent was obtained as required. GSCs were maintained under serum-free, adherent conditions.

Primary hippocampal neurons were obtained from P2–P3 Wistar rats (Charles River) and maintained in Neurobasal-A medium supplemented with GlutaMAX and B27 at 37°C in a humidified atmosphere containing 5% CO₂.

Further details of cell isolation, culture conditions, media composition, and neuronal preparation are provided in the Appendices.

### Electrophysiological recordings

GSCs and U87 cells were subjected to whole-cell voltage-clamp recordings using an inverted microscope and a MultiClamp 700B amplifier. Different intracellular and extracellular solutions were used to characterize membrane currents, assess ionic permeability, and investigate the effects of pharmacological inhibitors and osmotic stimulation. Recordings were acquired using Clampex software and analyzed with IGOR Pro. Further details of the recording procedures, solution compositions, and experimental conditions are provided in the Appendices.

### Calcium imaging experiments

Primary hippocampal neurons were pretreated with vehicle (DMSO) or the drug cocktail for 72 h before calcium imaging. Intracellular Ca²⁺ signals were monitored using Fluo-4 AM and epifluorescence microscopy during continuous perfusion with oxygenated artificial cerebrospinal fluid (ACSF). Fluorescence was acquired at 5 Hz, quantified in individual neuronal regions of interest, and expressed as ΔF/F₀. Neuronal responsiveness was evaluated following 25 mM K⁺ stimulation. Further details of dye loading, image acquisition, fluorescence analysis, and response criteria are provided in the Appendices.

### RT-PCR

Quantitative real-time PCR was used to evaluate the expression of VRAC subunits: LRCC8A, LRCC8B, LRCC8C, LRCC8D and LRRC8E. The RNeasy Mini Kit (Qiagen, cat.74104,) was used for isolation of total RNA from pellet of 5,0 × 10⁵ cells. After RNA extraction, a TURBO DNA-free Kit (MilliporeSigma, AM1907) was used to remove traces of genomic DNA. Then, cDNA synthesis was performed by Maxima First Strand cDNA Synthesis Kit (Thermo Fisher Scientific, K1641) using MiniAmp Thermal Cycler (Thermo Fisher Scientific); then used for quantitative real-time PCR using SYBR Green Supermix (Bio-Rad, 1725270) on a CFX Duet Real-Time PCR System (CFX Maestro Software) (Bio-Rad Laboratories). Expression levels were normalized to actin and predesigned primers were used (Sigma-Aldrich, St. Louis, MO, United States), their sequences are reported in Table 2, and fold change was determined by application of the 2^−ΔΔCt^ method.

**Table 2:**
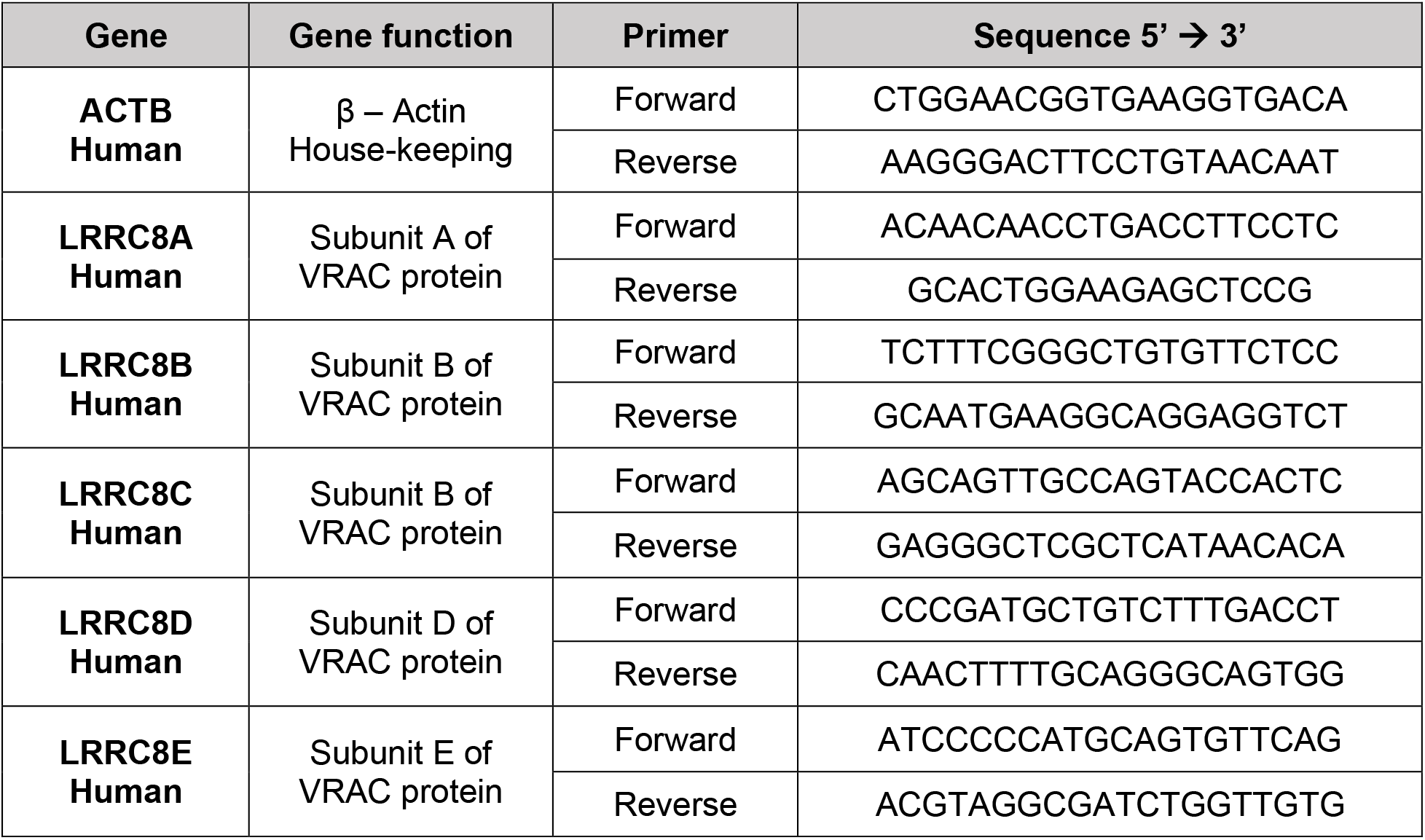
Primer sequences.

### Immunostaining

Patient-derived GSCs (S670) were fixed with 4% paraformaldehyde, permeabilized with Triton X-100, and blocked with bovine serum albumin. Cells were incubated with a mouse monoclonal anti-Na⁺/Ca²⁺ exchanger (NCX) antibody (C2C12, 1:200), followed by an Alexa Fluor 488-conjugated goat anti-mouse secondary antibody (1:600). Nuclei were counterstained with DAPI, and coverslips were mounted using VECTASHIELD Vibrance Antifade Mounting Medium. Images were acquired using a Nikon A1R confocal microscope with a 40×/0.95 NA objective. Further details of the staining procedure and image acquisition are provided in the Appendices.

### Survival assays

Cells (5,000 cells/well for the U87 cell line and 10,000 cells/well for GSCs cells) were seeded in 96-well plates and, after 48 h, treated for 24, 48, and 72 h with different concentrations of NFA (83), BTN (84), R-K (85) or cocktail (from 25 to 800 µM), SN6 (from 1.25 to 50 µM) (86), or Doxo (0.25 to 8 µM) (87). To evaluate the effects of liposomes, U87 cells were treated with empty liposomes (from 1 to 80 µg/mL) and then with 20 µg/mL of liposomes loaded with 50, 100, and 200 µM of cocktail. Two hours before the end of the incubation, Resazurin sodium salt (Merck, Darmstadt, Germany, 199303-5G) was added to each well at a final concentration of 350 µg/mL. Absorbance was detected at 560/590 nm using a Synergy H1 Hybrid Reader (BioTek, Agilent Technologies, Santa Clara, CA, USA).

### Biomechanical measurements

AFM force spectroscopy was used to quantify cellular stiffness, expressed as the Young’s modulus (88, 89). U87 cells and patient-derived GSCs were seeded onto glass coverslips at 0.3 × 10⁶ cells/coverslip, fixed with 4% paraformaldehyde for 20 min at room temperature, and stained with DAPI for cell identification. Measurements were performed using an NT-MDT Smena AFM equipped with 20 µm silicon spherical probes (CSG01 cantilever; k = 0.006–0.007 N/m). For each cell, force– distance curves were acquired over the nuclear region using an applied force of 0.7 nN, an indentation rate of 2 µm/s, and a maximum indentation depth of 500 nm. Elastic modulus values (E, kPa) were obtained by fitting the curves with a Hertz contact model for a spherical indenter (90) using AtomicJ software. Three biological replicates were performed, with four coverslips per condition per replicate and 100 cells analyzed per coverslip, corresponding to 1,200 cells analyzed per condition overall. Data are presented as the mean of the three biological replicates.

### Liposome Formulation and Functionalization

Liposomes were prepared using DPPC, cholesterol, DPPG, and DSPE-PEG2000-COOH at a molar ratio of 50:30:10:10, respectively. The lipid components were mixed with the individual cocktail compounds at the indicated concentrations (50, 100, and 200 µM), and a thin lipid film was formed by solvent evaporation under nitrogen. The film was hydrated with MES buffer and subjected to five freeze–thaw cycles to obtain multilamellar vesicles, followed by extrusion through 100 nm polycarbonate membranes to produce unilamellar liposomes. For transferrin functionalization, surface carboxyl groups were activated using EDC/NHS chemistry and conjugated to human holo-transferrin. Unbound transferrin and residual reagents were removed by centrifugal filtration. Further details of liposome preparation and functionalization are provided in the Appendices.

### Dynamic Light Scattering (DLS)

DLS was used to determine the size distribution of liposomes prepared. Following preparation, each liposome suspension was diluted to 0.1 μg/mL total lipid concentration using MilliQ water. and each sample was analyzed using a DynaPro NanoStar DLS (Wyatt Technologies, Santa Barbara, CA) with temperature control set at 20°C. 5 acquisitions were averaged and analyzed using the Hollow Sphere model in the DYNAMICS software package (Wyatt Technologies).

### Vesicles characterization

Liposomes were characterized by Nanoparticle Tracking Analysis (NTA) to determine their hydrodynamic size and particle concentration, and by Atomic Force Microscopy (AFM) to assess their morphology and dimensions. AFM was also used to investigate supported lipid bilayer formation and temperature-dependent membrane organization. For transferrin-functionalized liposomes, the associated protein content was quantified using the Bradford assay. Drug encapsulation efficiency for niflumic acid, bumetanide, and (R)-ketorolac was determined by UV–Vis spectrophotometry following liposome disruption. Further details of the experimental procedures and analytical conditions are provided in the Appendices.

### Chemicals and Pharmacology

(R)-Ketorolac (SML1654-5MG), purity ≥ 95% (HPLC), Bumetanide (B3023-250MG), purity ≥ 98% (TLC), Niflumic Acid (Y0000824), Doxorubicin Hydrochloride (D1515-10MG) (Merck, Darmstadt, Germany), SN6 (2184/50 Bio-Techne, Minneapolis, MN, United States) purity ≥98% (HPLC) were dissolved in dimethyl sulfoxide (DMSO, 34869 Sigma-Aldrich, St. Louis, MO, United States) at a 50 mM stock solution, divided into aliquots and stored at −20 °C protected from light. Working solutions were freshly prepared before each experiment by diluting the stock solution in the appropriate assay buffer.

### Statistics

All data are presented as mean ± standard deviation (SD) or mean ± standard error of the mean (SEM), as indicated in the respective figure legends. Statistical analyses were performed using GraphPad Prism (versions 7 and 8.0; GraphPad Software, San Diego, CA, USA) and RStudio (version 4.2.2). Comparisons between two independent groups were performed using a two-tailed unpaired Student’s *t*-test or the Mann–Whitney U test, as appropriate. For paired comparisons, the Wilcoxon signed-rank test was used when a non-parametric approach was appropriate. Comparisons involving three or more groups were performed using one-way or two-way analysis of variance (ANOVA), as appropriate, followed by Bonferroni’s or Tukey’s post hoc multiple-comparison tests. For non-normally distributed data involving three or more independent groups, the Kruskal– Wallis test was used. Homogeneity of variances was assessed using the F-test or the Brown– Forsythe and Bartlett’s tests, as appropriate. A *p*-value < 0.05 was considered statistically significant.

### Sex as a biological variable

Sex was not considered a biological variable in this study. Further information on the patients from whom GSCs were derived is provided in the Appendices, in accordance with the privacy policies of the institutions involved.

## Data availability

All materials and data from this study are available to researchers upon reasonable request and after issuance of relevant institutional data sharing agreements by contacting the corresponding authors. Human data with identifiers are restricted owing to patient confidentiality.

## Study approval

The protocol for human samples was approved by the local Ethics Committee, Comitato Etico Unico Regionale del Friuli Venezia Giulia (protocol 718345, opinion 196/2014/Em) and the Padova University-Hospital (2462P)

The use of primary neurons was approved by the Local Veterinary Service (Trieste, Italy), by the SISSA Ethics Committee board, and by the National Ministry of Health. All the procedures were conducted according to the guidelines of the Italian Animal Welfare Act and to the European Union guidelines for animal care (d.1.116/92; 86/609/C.E.).

## Manuscript Information

### Author contributions

C.A.S.T., A.N., and A.S. contributed equally to this work and are co–first authors. R.S., C.A.S.T., and V.T. conceived and designed the research. C.A.S.T. performed and analyzed the electrophysiological experiments, calcium imaging, and immunohistochemical analyses. A.N. and A.S. performed and analyzed the biochemical and molecular experiments. A.N. also performed the biomechanical experiments and formulated and functionalized the liposomes. C.A.S.T., A.N., A.S., and R.S. performed experiments and acquired data. C.A.S.T., A.N., and A.S. analyzed and interpreted the results. D.C., I.M., T.I., E.R., L.P., and M.S. provided essential GSCs and scientific advice. P.P., L.C. (Collavin), L.C. (Casalis), A.M., and R.S. provided technical and scientific support. P.P. and L.C. (Casalis) characterized the biophysical properties of the vesicles. A.M. supervised the electrophysiological experiments. C.A.S.T., A.N., A.S. and V.T. drafted the manuscript. C.A.S.T. and V.T. supervised the study and critically revised the manuscript. R.S. and V.T. provided financial support. All authors reviewed and approved the final version of the manuscript.

### AI disclosure

ChatGPT (OpenAI) was used to assist with English language editing and improve the clarity, readability, and grammatical accuracy of the manuscript. The authors reviewed and approved all suggested revisions and take full responsibility for the manuscript’s content.

### Funding

This work was supported by the *Booster for Life Science FVG – TRL Advancement* programme, which supports projects aimed at validating innovative ideas and technologies and advancing them to Technology Readiness Levels (TRL) 6, 7, or 8.

## Supplementary Captions

**Supplementary Figure 1.**
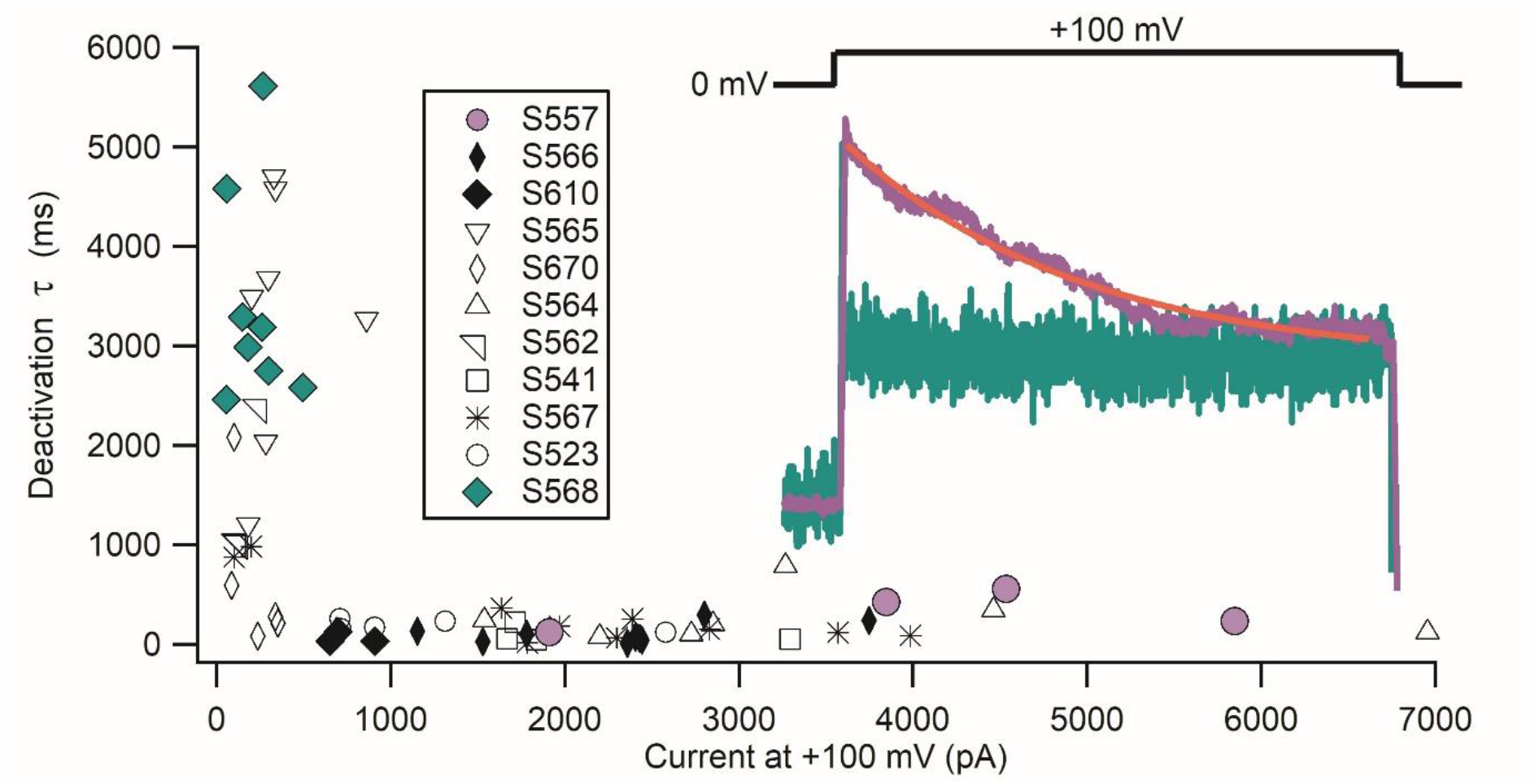
Deactivation time constants as a function of the maximal current amplitude of VRAC-like current measured at +100 mV, showing the heterogeneity of the VRAC current activated by calcium in GSCs.

**Supplementary Figure 2.**
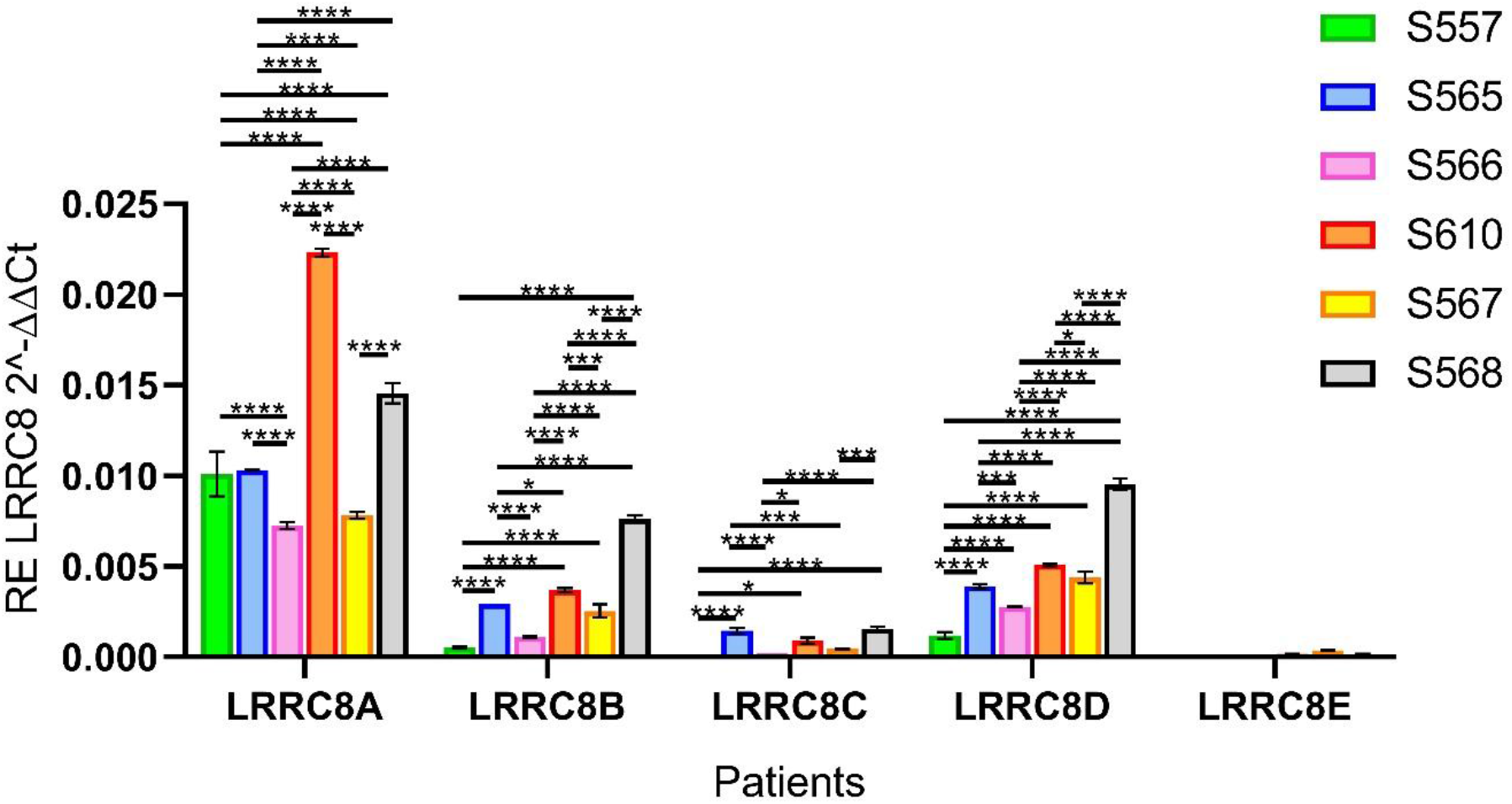
Differential expression of the five VRAC subunits across six patients, evaluated by real-time PCR, showing high heterogeneity in LRRC8A, LRRC8B and LRRC8D expression (One-Way ANOVA, Sidak’s post-test).

**Supplementary Figure 3.**
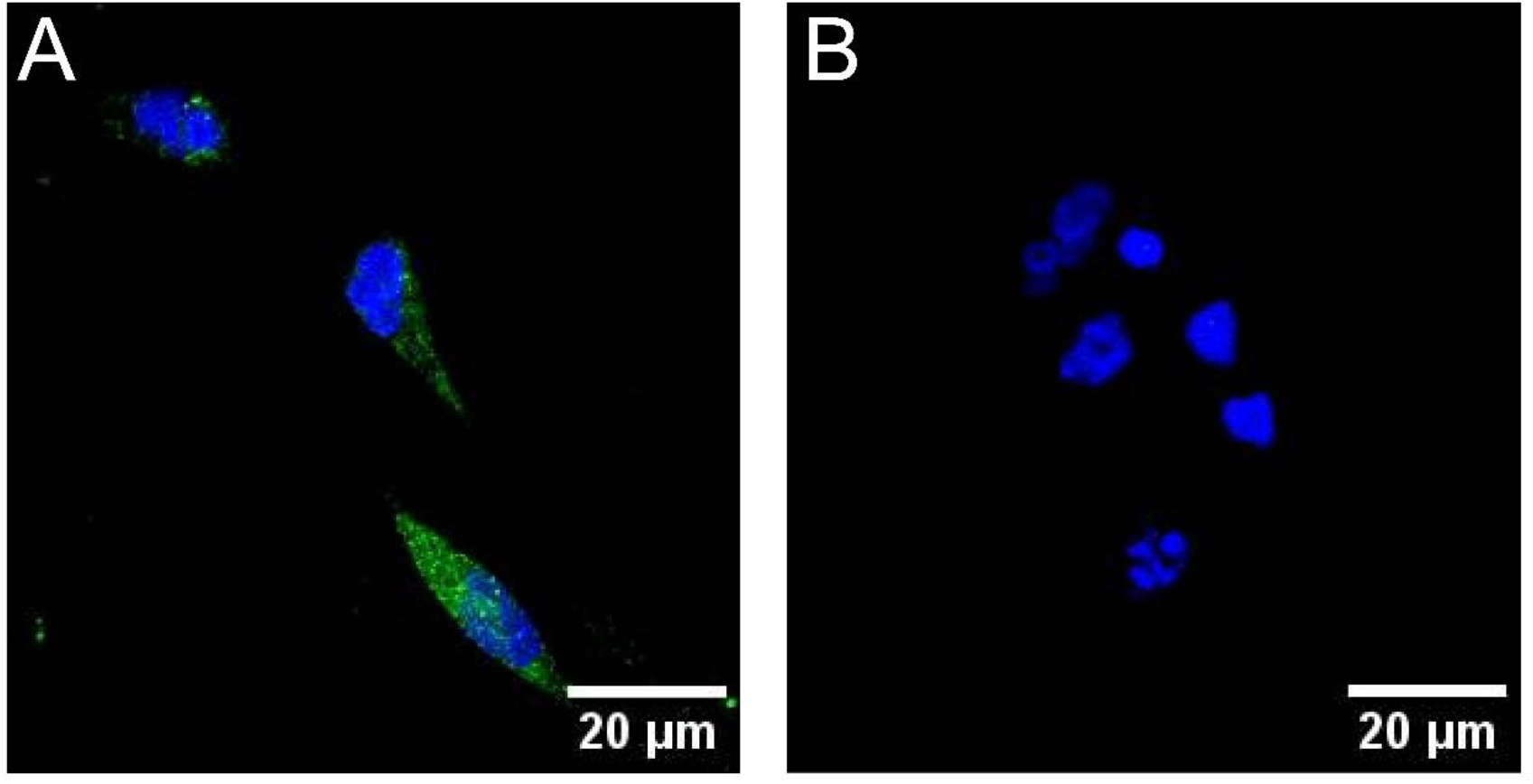
Immunostaining of patient-derived GSCs from patient 670 for the Na⁺/Ca²⁺ exchanger (NCX). **(A)** GSCs immunostained with an anti-NCX primary antibody, showing green fluorescence consistent with membrane-associated NCX expression. **(B)** Negative control processed with the Alexa Fluor 488-conjugated secondary antibody alone.

**Supplementary Figure 4.**
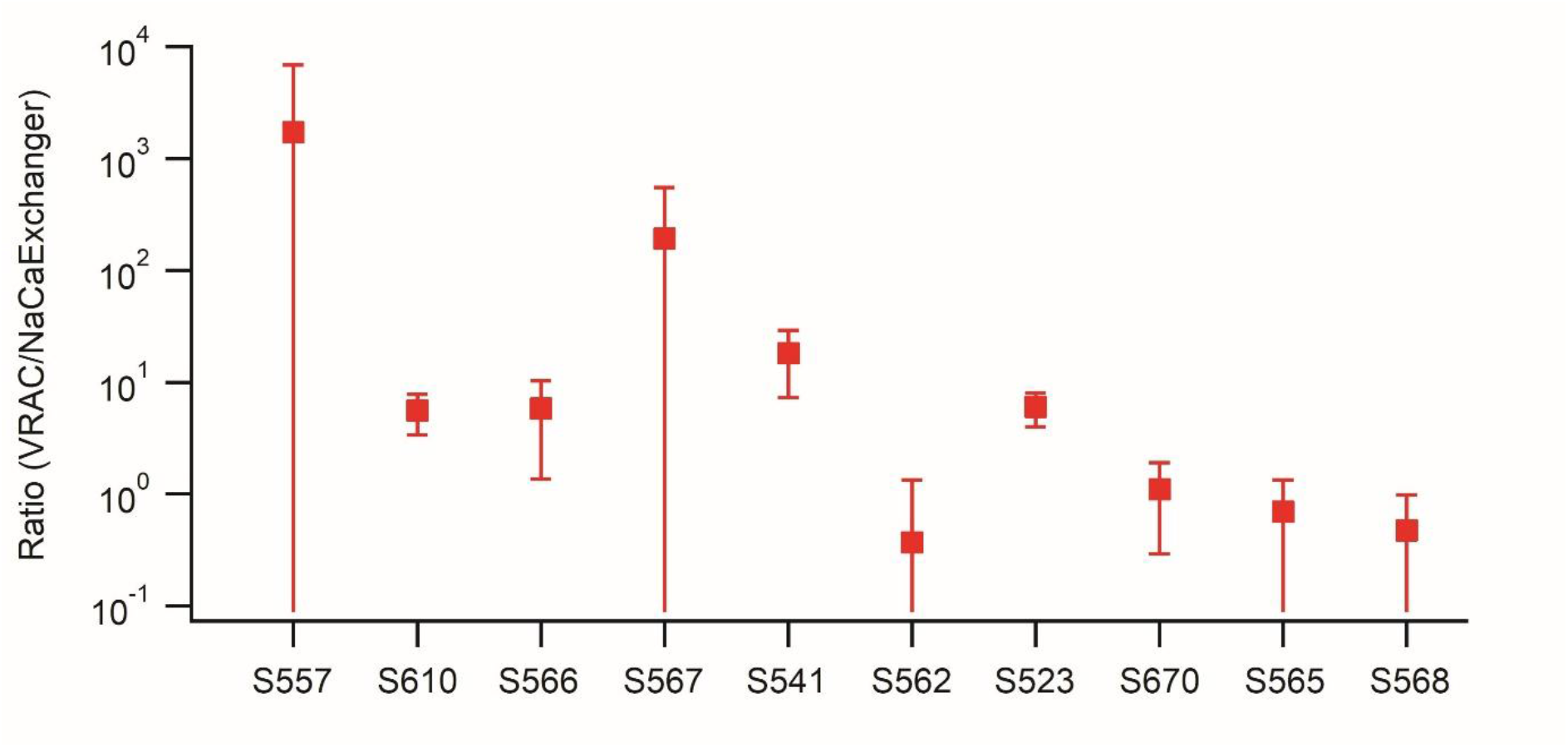
Ratio of VRAC-like current amplitude to NCX-associated current amplitude across GSC cultures derived from 10 patients. Ratios greater than 1 indicate a larger VRAC-like current than NCX-associated current. Patients S557, S610, S566, S567, S541, and S523 exhibited ratios greater than 1, indicating a marked predominance of the VRAC-like current.

**Supplementary Figure 5.**
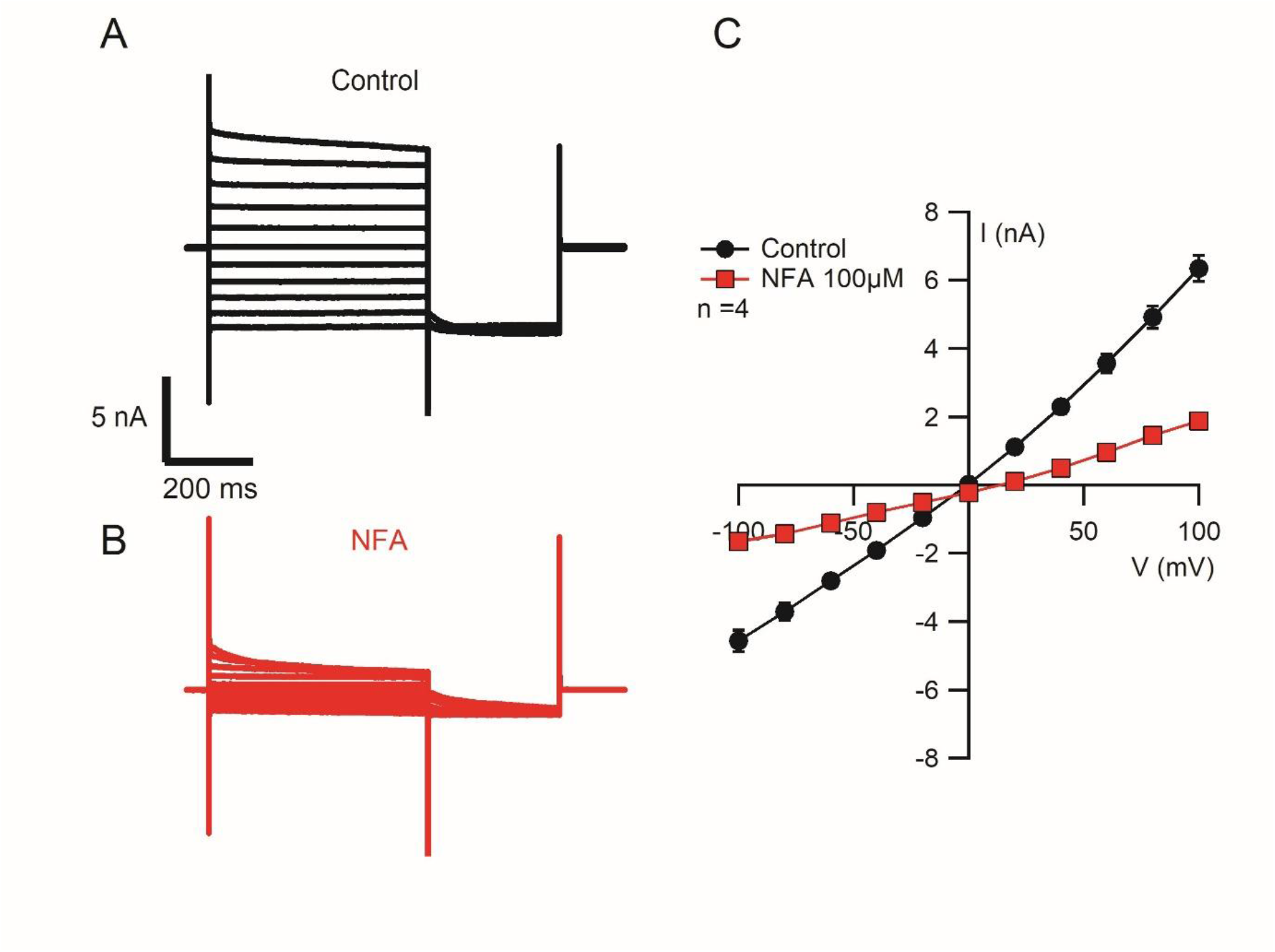
U87 cells exhibit a calcium-dependent VRAC-like current that is inhibited by niflumic acid (NFA). **(A)** Representative current traces recorded from a U87 cell held at 0 mV and subjected to voltage steps from −100 to +100 mV, showing the characteristic kinetics of the VRAC-like current when the pipette solution contained 0.5 µM Ca²⁺. **(B)** Current traces recorded from the same cell following treatment with 100 µM NFA, showing marked current inhibition at both positive and negative membrane potentials. **(C)** Current–voltage (I– V) relationship recorded before (black) and during (red) application of 100 µM NFA.

**Supplementary Figure 6.**
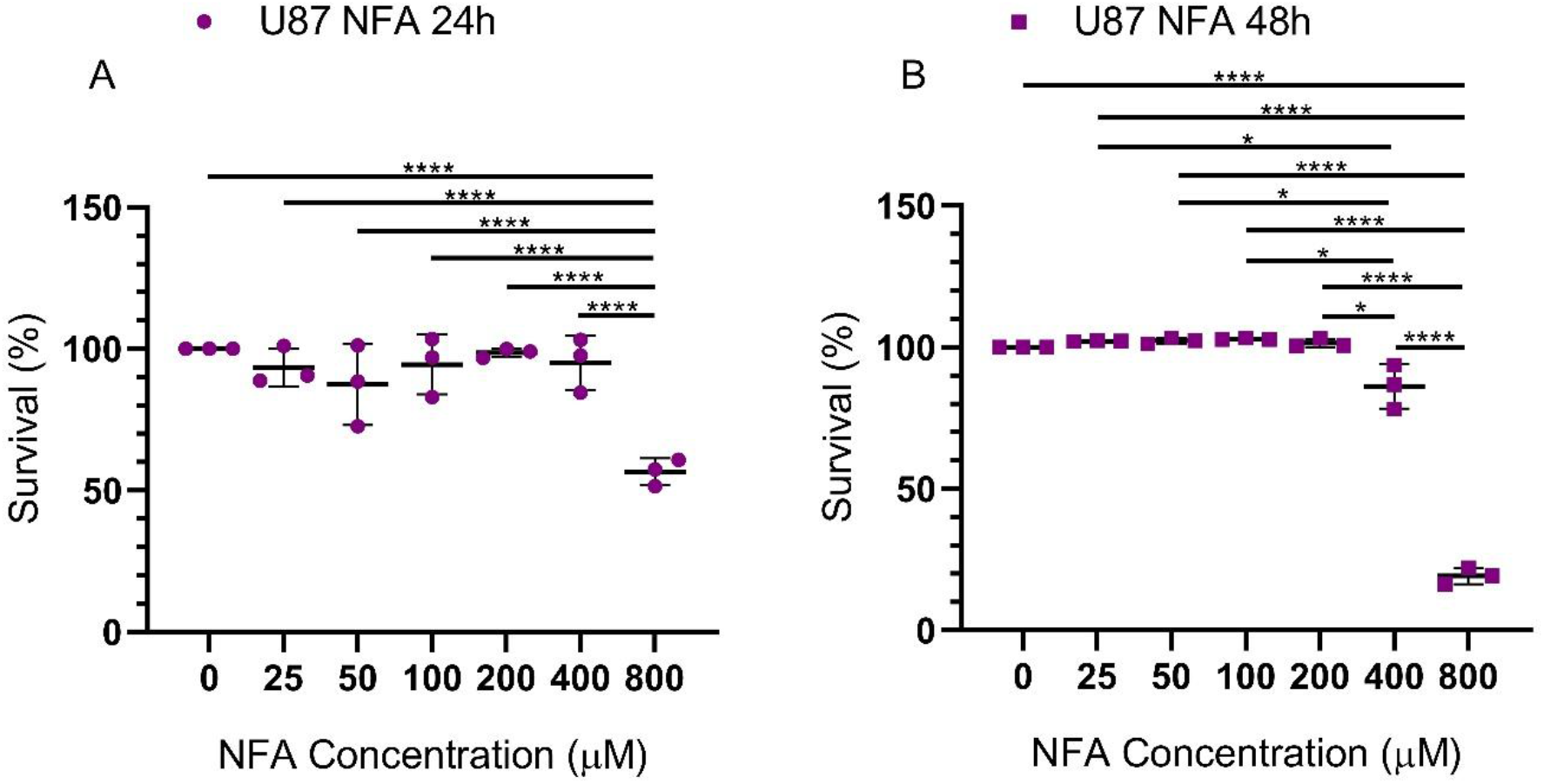
Cell viability following 24 h (**A**) and 48 h (**B**) of NFA treatment. At 24 h, a significant reduction is observed only at 800 *μ*M compared to all other concentrations (p < 0.000001; one-way ANOVA, Sidak’s post-test), whereas at 48 h, a significant reduction is evident at 400 and 800 *μ*M compared with all other conditions (p < 0.05 and p < 0.00001, respectively; one-way ANOVA, Sidak’s post-test).

**Supplementary Figure 7.**
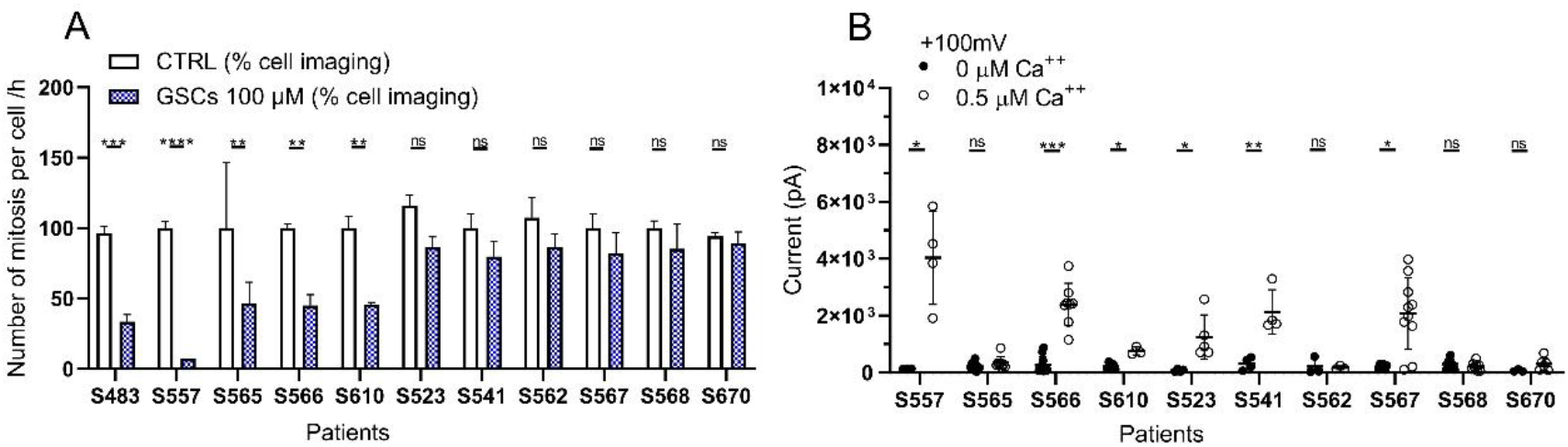
**(A)** Sensitivity of 11 different patient-derived GSCs to 100 µM cocktail, evaluated by measuring the number of mitoses per cell per hour via live-cell imaging. 45% of patients (5 out of 11) showed a significant response compared to non-responsive patients. (**B**) Measurement of VRAC-like currents in 10 different patient samples, showing a significant difference between the absence and presence of 0.5 µM Ca²⁺. Six out of the ten patients showed a significant VRAC like current.

**Supplementary Figure 8.**
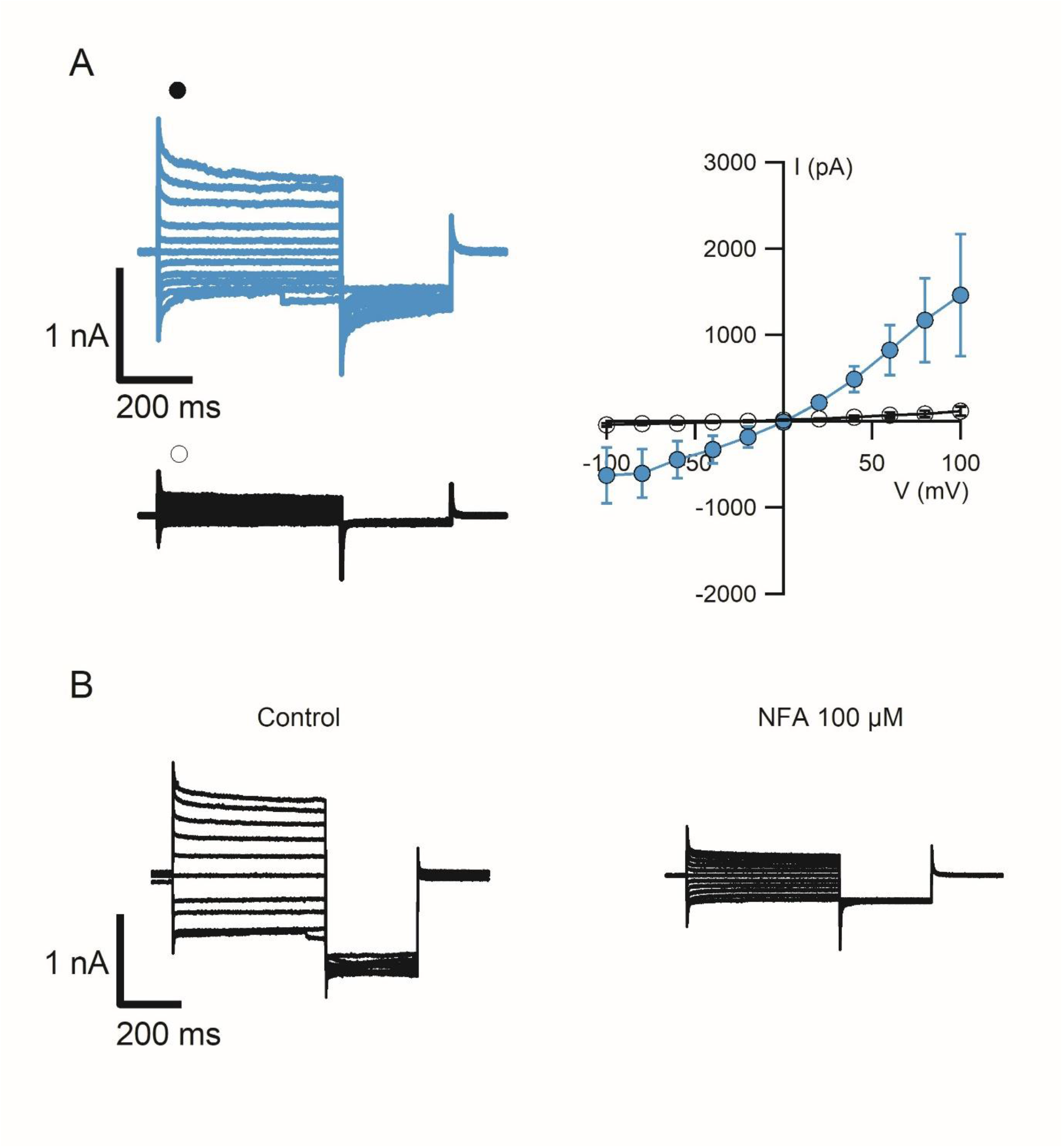
Patient-derived GSCs from patient S82 exhibit a calcium-dependent VRAC-like current that is inhibited by niflumic acid (NFA). **(A)** Representative current traces recorded with pipette solutions containing 0.5 µM Ca²⁺ (blue) or nominally zero Ca²⁺ (black) (left). The corresponding current–voltage (I–V) relationships for the two conditions are shown on the right (n = 3–4 cells). **(B)** Representative current traces recorded from a GSC before and after application of NFA, showing inhibition of the VRAC-like current at both positive and negative membrane potentials.

**Supplementary Figure 9:**
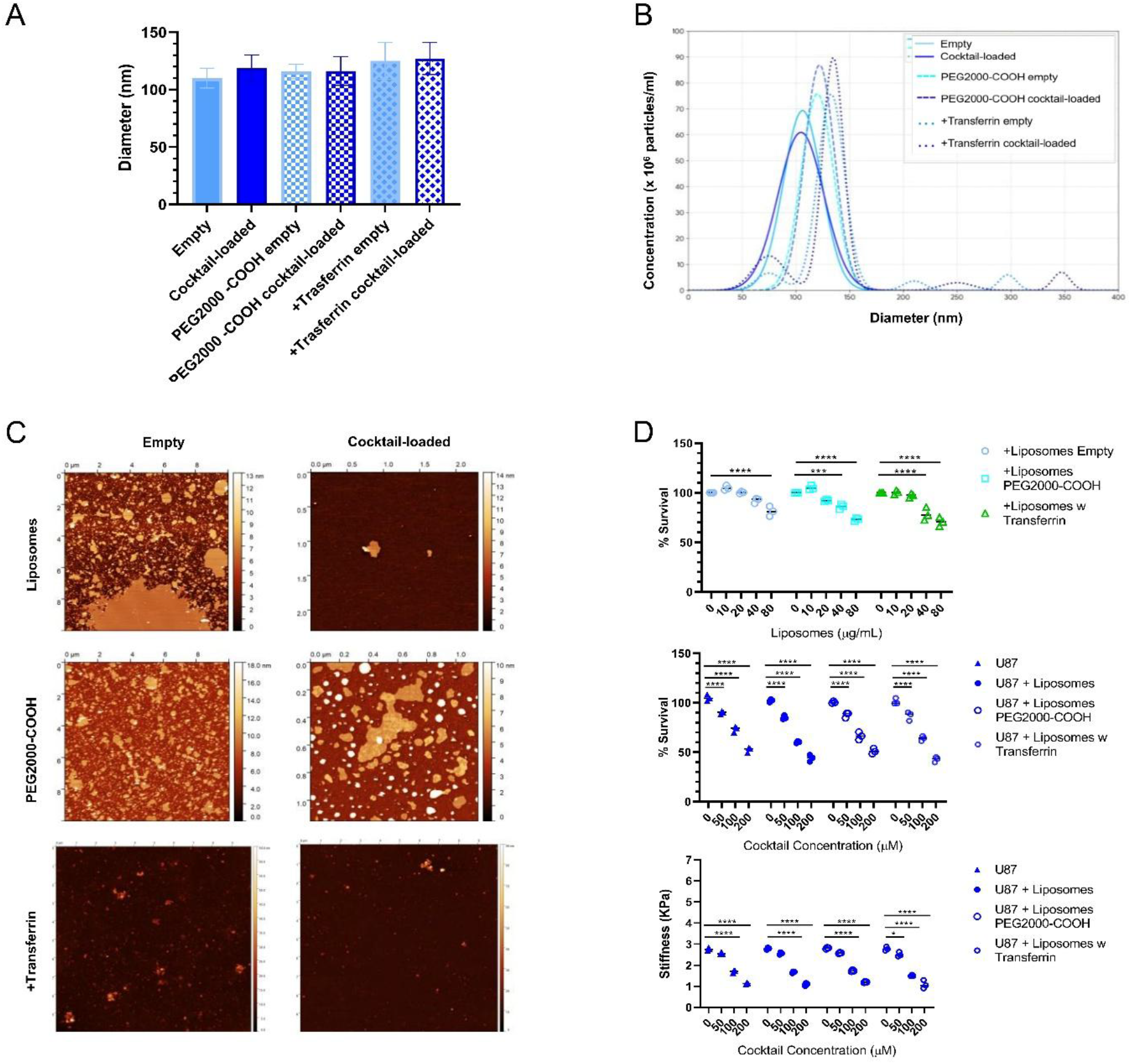
Characterization and effects of liposomes. (**A**) Evaluation of the sizes of liposomes, PEGylated liposomes, and transferrin-functionalized liposomes, comparing empty and cocktail-loaded nanoparticles by DLS. (**B**) Comparison of the size and concentration of liposomes, PEGylated liposomes and transferrin-functionalized liposomes, empty and cocktail-loaded by NTA. (**C**) Representative AFM images show the chemical and physical properties of the liposome and PEG-conjugated liposomes formulations and the morphology of transferrin-functionalized liposomes, comparing empty and cocktail-loaded nanoparticles. (**D**) Graphs show the effects of liposomes on the U87 cell line after 72 hours of treatment. Specifically, the first graph displays the cytotoxic effect of empty liposomes, PEGylated liposomes, and transferrin-functionalized liposomes, used to establish the maximum tolerated dose of nanoparticles formulations (20 µg/mL). The second graph compares the cytotoxic effect of the three different concentrations of the cocktail (50, 100 and 200 µM) and the 20 µg/mL of cocktail-loaded liposomes, PEGylated liposomes, and transferrin-functionalized liposomes. The third graph illustrates the effects of the cocktail (50, 100 and 200 µM) and the 20 µg/mL of cocktail-loaded liposomes, PEGylated liposomes, and transferrin-functionalized liposomes on cellular biomechanical properties.

**Supplementary Figure 10.**
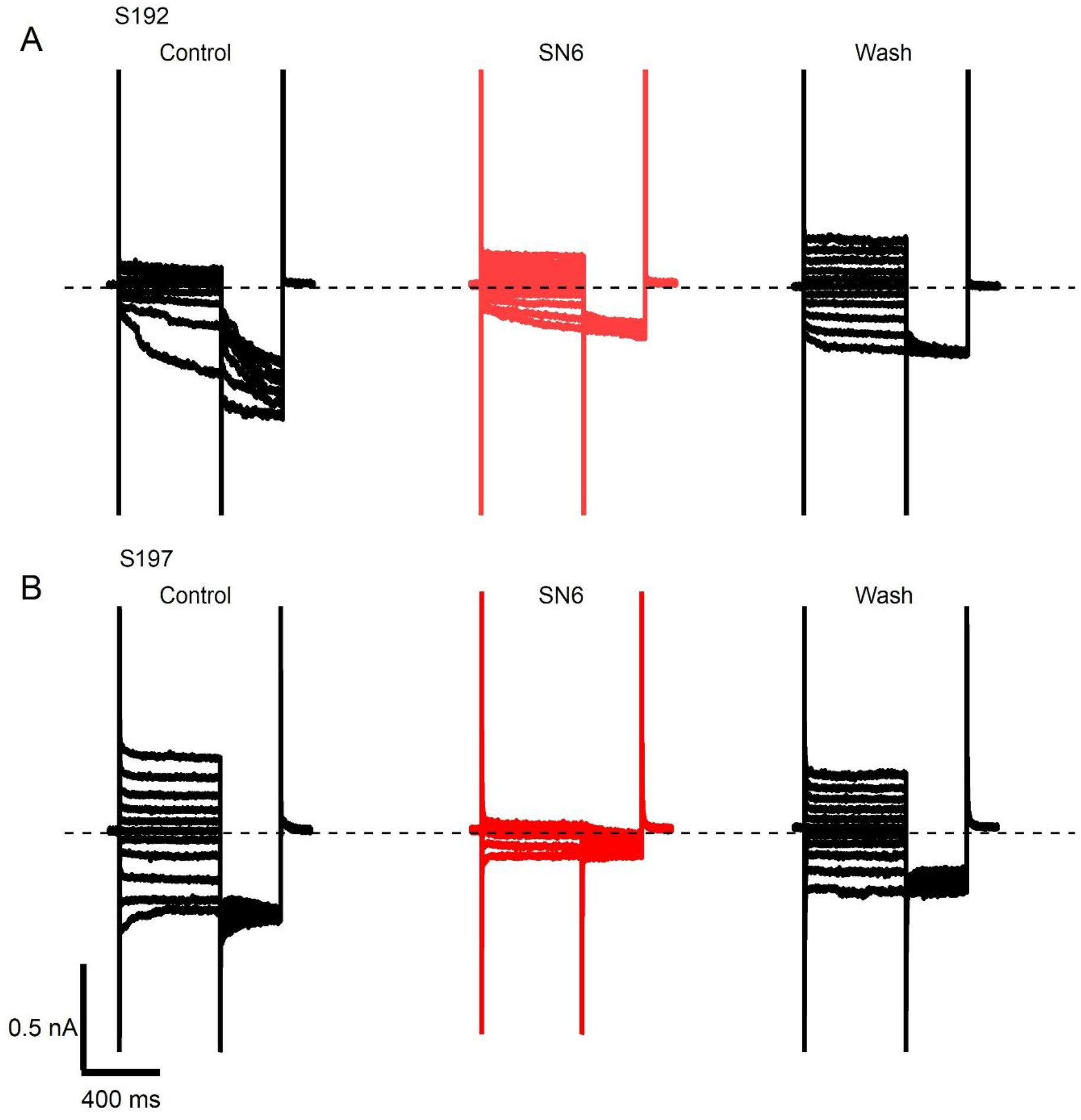
Patient-derived GSCs from patients 192 (**A**) and 197 (**B**) exhibit an inward current consistent with Na⁺/Ca²⁺ exchanger (NCX) activity, which is inhibited by the NCX inhibitor SN-6 at 20 uM.

**Supplementary Figure 11.**
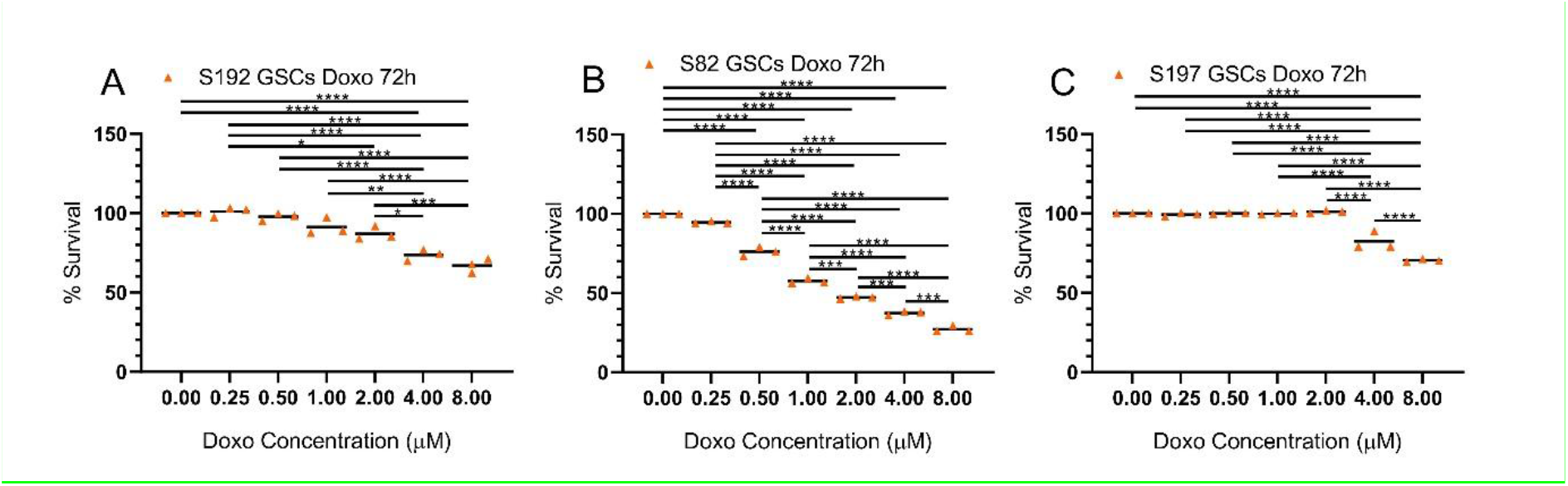
(**A**) Cell viability in S192 GSCs following 72 h of Doxorubicin treatment after 8 culture passages, showing reduced sensitivity to doxorubicin, with a significant difference compared with the untreated control at 4 µM and 8 µM (p = 0.000004 and p < 0.000001 respectively; one-way ANOVA, Sidak’s post-test). (**B**) Cell viability in S82 GSCs following 72 h of Doxorubicin treatment, showing reduced sensitivity to doxorubicin, with a significant difference compared with the untreated control from 0.5 µM to 8 µM (p = 0.000001 for all; one-way ANOVA, Sidak’s post-test). (**C**) Cell viability in S197 GSCs following 72 h of Doxorubicin treatment, showing reduced sensitivity to doxorubicin only at 4 µM and 8 µM, with a significant difference compared with concentrations ranging from 0 to 2 µM (p < 0.000001 for both; one-way ANOVA, Sidak’s post-test).

## Appendices

### Cell culture

Human U87-MG glioblastoma cells (RRID: CVCL_0022) and HMC3 human microglial cells were obtained from the American Type Culture Collection (ATCC, Manassas, VA, USA). U87-MG cells were cultured under adherent conditions in high-glucose Dulbecco’s modified Eagle’s medium (DMEM-HG; ECM0728L EuroClone, Milan, Italy), whereas HMC3 cells were maintained in Minimum essential medium (MEM; ECM0430L EuroClone, Milan, Italy). Both media were supplemented with 10% fetal bovine serum (FBS; F9665-500ML Sigma-Aldrich, Milan, Italy), 1% 200 mM L-glutamine (ECB3000D EuroClone, Milan, Italy), 1% penicillin (10,000 IU/mL; ECB3001D EuroClone, Milan, Italy), and 1% streptomycin (10 mg/mL; ECB3001D EuroClone, Milan, Italy). Cells were incubated at 37 °C in a humidified atmosphere containing 5% CO₂ and passaged when they reached approximately 80% confluence. Both cell lines were tested for mycoplasma contamination and confirmed to be negative.

Patient-derived glioma stem cells (GSCs) were isolated from human glioblastoma (GBM) surgical specimens collected by the Department of Neurosurgery of the Azienda Ospedaliero-Universitaria di Udine and, where applicable, the Department of Neurosurgery of the University Hospital of Padova, Italy. GSC isolation and expansion were performed according to previously described protocols (78, 79). The studies involving human specimens were conducted in accordance with the Declaration of Helsinki and were approved by the relevant ethics committees. The Udine study was approved by the Independent Ethics Committee of the University Hospital of Udine (approval no. 196/2014Em); the corresponding ethics approval details for specimens collected in Padova should be reported separately. All applicable informed-consent requirements were fulfilled.

GSCs were maintained under serum-free, adherent culture conditions using the established laboratory-specific culture protocols described below.

Primary hippocampal neuron cultures were prepared from postnatal day 2–3 (P2–P3) Wistar rats (Charles River). Hippocampal tissue was enzymatically dissociated using a solution containing 136.9 mM NaCl, 4.9 mM KCl, 7 mM Na₂HPO₄, 25.2 mM HEPES, 4.2 mM NaHCO₃, 200 μM kynurenic acid, and 25 μM D(-)-2-amino-5-phosphonopentanoic acid (all from MilliporeSigma). Following dissociation, the neuronal suspension was centrifuged at 100 × g for 5 min. The resulting cells were plated onto 15-mm glass coverslips (Menzel-Gläser, CB00150RA1) pre-coated with 0.5 μg/mL poly-L-ornithine (MilliporeSigma, P4957). Neurons were maintained in Neurobasal-A medium (Thermo Fisher Scientific, 21103049) supplemented with GlutaMAX Supplement (Thermo Fisher Scientific, 35050061), gentamicin solution (1 mg/mL; MilliporeSigma, G1272), and B27 Supplement (Thermo Fisher Scientific, 17504044). Cultures were maintained at 37°C in a humidified incubator with 5% CO₂ and 95% relative humidity.

#### Udine protocol

GSCs were isolated from dissociated GBM tissue and cultured under adherent, serum-free conditions on laminin-coated dishes. Cells were maintained in Neurobasal-A medium (Gibco, Invitrogen; cat. no. 10888) supplemented with 2 mM L-glutamine (Sigma-Aldrich; cat. no. G7513), B-27 supplement (Gibco; cat. no. 17504-044), N-2 supplement (Gibco; cat. no. 17502-001), 25 μg/mL insulin (Sigma-Aldrich; cat. no. I2643), 20 ng/mL human EGF (Peprotech; cat. no. AF-100-15), and 20 ng/mL human basic FGF (Peprotech; cat. no. 100-18B). This culture approach was based on the previously published protocol for the isolation and expansion of regionally defined human GBM cells (79).

GSC cultures were periodically tested for mycoplasma contamination. Their stem-like phenotype was routinely evaluated by immunostaining for Nestin and Vimentin, together with assessment of the intermediate-filament characteristics associated with an undifferentiated phenotype. Stemness was further assessed by examining the expression of OCT4, NANOG, and SOX2, as previously (80–82).

#### Padova protocol

Alternatively, GSCs were seeded at a density of 1.5–2.0 × 10⁴ cells/cm² onto fibronectin-coated culture plates and grown as adherent monolayers in DMEM/F12 medium (EuroClone, Milan, Italy) supplemented with 10% BIT9500 (STEMCELL Technologies, Vancouver, Canada), 20 ng/mL basic fibroblast growth factor (bFGF), and 20 ng/mL epidermal growth factor (EGF; both from Immunological Sciences, Rome, Italy). Cultures were maintained in a humidified incubator under hypoxic conditions consisting of 2% O₂ and 5% CO₂, with nitrogen as the balance gas.

### Electrophysiological recordings

GSCs and U87 cells were seeding (3,000 cells/well) on coverslips and viewed on an inverted microscope (Olympus, IX70), using a 40x objective, the cells were chosen randomly and recorded in voltage-clamp whole-cell configuration. Patch electrodes had a 3 to 5 MΩ resistance and were pulled from borosilicate capillaries (World Precision Instruments) with a PC-10 puller (Narishige); the electrodes were filled with an intracellular solution containing (in mM): 130 CsCl, 10 NaCl, 10 HEPES, 5 EGTA adjusted to pH 7.2 with CsOH. In some experiments to reverse the exchanger current CsCl was replaced by NaCl. The nominally 0 Ca^2+^ solution contained no added Ca^2+^ and the 0.5 Ca^2+^ solution by adding 3.85 mM of CaCl2 to the same solution. Electrophysiological recordings were made using a MultiClamp 700B amplifier controlled by Clampex 10.6 via a Digidata 1550B (Molecular Devices); the sample frequency used to acquire data was 10 kHz with low-pass filtering at 2 kHz.

Statistical analysis of maximum currents, reversal potentials and I-V curves plots was made by IGOR pro (WaveMet rics, https://www.wavemetrics.com/). The extracellular ringer contained (in mM): 120 mM NaCl, 20 mM NaHCO_3_, 3 mM KCl, 2 mM CaCl_2_, 1 mM MgSO_4_, 10 mM HEPES, and 10 mM glucose, pH 7.4. and osmolarity to 310 mOsm, In some experiments to check the ionic permeability were used 2 types of solutions: one to assess the cationic or anionic permeability which had 14 mM of NaCl and the rest of components were replaced by sucrose, where before of each experiment was checked the osmolarity to be equal to the ringer solution, then in other set of experiments to check the sodium permeability, the NaCl and NaHCO_3_ were completely replaced by NMDG-Cl. The bath was grounded through a 3 M KCl agar bridge connected to an Ag/AgCl reference electrode. To activate the VRAC-like current by a natural stimuli, cells from patient 557 were perfused with a hypotonic solution reached by decreasing the concentration of NaCl to 90 mM and leaving the other component equally to the original extracellular ringer. To change solutions, the setup was equipped with six-barrel square glass (3SG700-5, Warner Instruments, Hamden, CT, USA) using the gravity-driven perfusion system Fast-Step SF-77B (Warner Instruments). All compounds and chemicals were obtained from MilliporeSigma, unless otherwise stated. All liquid junction potentials for the solutions used were calculated using the pClampex software (Axon Instruments), obtaining values between 1.2 and 5.8 mV.

### Calcium imaging experiments

Mouse Hypocampal neurons were pretreated with vehicle DMSO or Cocktail of drugs 72 hours before the calcium imaging recordings, solution containing the Ca2+ indicator Fluo-4 AM (5 μM; Thermo Fisher Scientific), 0.001% pluronic F-127 (20% solution in DMSO; Thermo Fisher Scientific), Ca²⁺ imaging experiments were performed using an inverted epifluorescence microscope (IX70; Olympus) equipped with a mercury fluorescence lamp (U-RF-T). Hypocampal neurons were placed in a recording dish on the microscope stage and continuously perfused by gravity-driven flow with oxygenated artificial cerebrospinal fluid (ACSF; 95% O₂ and 5% CO₂). Fluorescence was detected using a U-MWIB2 filter block (excitation filter, EX 460–90; dichroic mirror, DM 505; emission filter, EM 510IF) and a 40× objective (UPlanFL; numerical aperture, 0.75; Olympus). Images were acquired using an ORCA-Flash4.0 V3 digital CMOS camera (C13440-22CU; Hamamatsu) controlled by HCImage Live software (Hamamatsu), at a sampling frequency of 5 Hz and a resolution of 512 × 512 pixels. Fluorescence changes were quantified in regions of interest (ROIs) delineated around individual neurons using ImageJ version 1.51s (National Institutes of Health). Fluorescence signals were normalized to the baseline fluorescence according to the following equation: ΔF/F₀ = [F(t) − F₀]/F₀, where F₀ represents the mean fluorescence intensity and F(t) represents the fluorescence intensity at time t. Subsequent data analysis and figure preparation were performed using Igor Pro version 6.3.7.2 (WaveMetrics). Neurons were classified as responsive if they respond to a stimulation with 25 mM K⁺ eliciting a Ca²⁺ transient, where ΔF/F₀ exceeded the mean prestimulus fluorescence level, calculated over a 15-s baseline period, by more than three standard deviations for at least 5 s. In some experiments, photobleaching-related fluorescence decay was corrected mathematically using the exponential decay profile observed in non-responsive cells. To avoid image saturation, excitation light intensity was attenuated by ND4 and ND8 neutral density filters (Nikon, Tokyo, Japan).

### Immunostaining

For immunofluorescence experiments GSCs from patient S670 were rinsed three times in PBS and then fixed in 4% paraformaldehyde for 10 min at RT and kept at 4 °C until staining was performed. For the immunofluorescence staining, fixed cultures were rinsed three times in PBS to remove PFA, then a glycine solution (glycine 1 mM in PBS) was added for 5 min to reduce autofluorescence. After permeabilization with 0.2% Triton X-100 for 5 minutes, cells were incubated with 5% bovine serum albumin blocking solution at RT for 50 minutes. Then, coverslips were incubated with the Mouse monoclonal Sodium/Calcium exchanger monoclonal antibody (C2C12; #MA3-926) in ratio of 1:200, After 3 washes with PBS–0.01% Tween 20, the coverslips were incubated with a goat anti-mouse secondary antibody conjugated to Alexa Fluor 488 (Invitrogen, Thermo Fisher Scientific, A11029; 1:600) for 1 hour at RT in a dark and wet chamber. To visualize nuclei, incubation with DAPI 0.5 mg/mL (Merck-Sigma, 32670) in PBS for 15 minutes at RT was performed. Cover-slips were mounted with VECTASHIELD Vibrance Antifade Mounting Medium (Vector Laboratories, H-1700). For image acquisition, an inverted Nikon A1R confocal microscope was used. Z-stacks were acquired using the NIS Elements Advanced Research Software (Nikon) with ×40/0.95 NA air objective and a spatial resolution of 1024 pixels.

### Liposome Formulation and Functionalization

DPPC (1,2-dipalmitoyl-sn-glycero-3-phosphocholine; Avanti Polar Lipids, cat. 850355P), cholesterol (Avanti Polar Lipids, cat. 700000P), DPPG (1,2-dipalmitoyl-sn-glycero-3-phospho-(1′-rac-glycerol); Avanti Polar Lipids, cat. 200880), and DSPE-PEG2000-COOH (1,2-distearoyl-sn-glycero-3-phosphoethanolamine-N-[carboxy(polyethylene glycol)-2000]; Avanti Polar Lipids, cat. 880135P) were resuspended using Hamilton syringes (Hamilton Company, Franklin, MA, USA) in chloroform (cat. 438613, Carlo Erba, Cornaredo, MI, Italy) to obtain a 1 mg/mL stock solution, which was stored at −20 °C. These components were mixed in a single-neck round-bottom flask (cat. Z548871, Sigma-Aldrich, St. Louis, MO, USA) at a molar ratio of 50:30:10:10 (DPPC:cholesterol:DPPG:DSPE-PEG2000-COOH) and combined with the individual components of the drug cocktail (50, 100, and 200 µM). The organic solvent was evaporated under a gentle nitrogen stream overnight to form a thin lipid film. The film was hydrated with MES buffer (cat. 28390, ThermoFisher Scientific, Waltham, MA, USA), pH 6.0, incubated at room temperature for 15 min, and vortexed vigorously (30 s, twice). The lipid suspension was rapidly frozen in liquid nitrogen (−196 °C) for 5 min and thawed in a lukewarm water bath (50 °C) for 5 min. This freeze-thaw cycle was repeated 5 times to obtain multilamellar vesicles (MLVs). Then, to produce unilamellar vesicles, the suspension was extruded through a 100 nm pore size polycarbonate membrane for 51 passes using an Avanti mini-extruder (Avanti Polar Lipids, Alabaster, AL, USA) according to the manufacturer’s instructions.

For transferrin functionalization, carboxyl groups on the liposomal surface (10% DSPE-PEG2000-COOH) were activated using EDC/NHS chemistry. Liposomes were incubated with 2 mM EDC (cat. PG82079, ThermoFisher Scientific) in MES buffer (pH 6.0) for 15 min at room temperature. Next, 20 mM 2-mercaptoethanol was added to quench the EDC. Then, 5 mM NHS (cat. PG82071, ThermoFisher Scientific) and 20 µg human holo-transferrin (cat. T0665, Sigma-Aldrich) were added to the liposomes and incubated overnight at 4 °C to allow the formation of amide bonds between the activated carboxyl groups and the lysine residues of transferrin. To stop the reaction, 20 mM Tris was added and incubated for 20 min at room temperature. Unconjugated transferrin and other reagents were removed using 100 kDa Amicon Ultra-15 centrifugal filter units (cat. UFC910024, Merck) by centrifugation at 4,000 × *g* for 40 min at 4 °C.

### Vesicles characterization

#### Nanoparticle Tracking Analysis (NTA)

A NanoSight NS300 (NanoSight Ltd., UK), equipped with a 488 nm blue laser and an automated sample assistant for automated loading, was used. Liposome samples were diluted in Milli-Q water at an appropriate concentration (1:200) to maintain particle numbers within the optimal instrument detection range (under the threshold of 100 particles per field of view). Seven 60 s videos were captured for each sample at camera level 15. After visual inspection, videos with visible artifacts were removed, and five videos per replicate were chosen for analysis. Raw laser scattering and Brownian particle movement data were analyzed using NTA software (version 3.3; NanoSight Ltd., UK). Automatic settings were selected for maximal track length, minimum expected particle size, and blur. The minimum track length was set to 10, detection threshold to 5, sample viscosity to the viscosity of water, and temperature to 25 °C. The output data were presented as liposome size (modal hydrodynamic diameter in nm) and liposome concentration (number of particles/mL per mg/mL of lipid mix).

#### Atomic Force Microscopy (AFM) Imaging

Mica is a flaky mineral characterized by a negatively charged surface, serving as a suitable substrate for AFM imaging of biological samples. Briefly, 20 µL of liposomes, resuspended in PBS, were deposited onto a mica surface previously functionalized with poly-L-lysine and washed 5 times with 100 µL of PBS to remove debris or non-adhesive material. After a 15 min incubation, a 50 µL drop of PBS was added. Image acquisition was performed using an MFP-3D atomic force microscope (Asylum Research/Oxford Instruments) with an Olympus Biolever Mini tip in liquid tapping mode. For each sample, 5 images were acquired (5 µm × 5 µm, 512 × 512 pixels). Images were analyzed using Gwyddion software, which allows discrimination of vesicles according to diameter and height. Data were subsequently processed using IGOR Pro and GraphPad Prism software.

#### Deposition of Supported Lipid Bilayers (SLBs)

The standard protocol for vesicle fusion to obtain a planar SLB was carried out by depositing 50 µL of the final liposome suspension onto a freshly cleaved mica substrate (Nano-Tec V-1 grade, 10 mm diameter, 0.15–0.21 mm thickness). Samples were incubated at 50 °C for 20 min and then slowly cooled to 45 °C at a rate of 0.07 °C/s. Excess unfused vesicles were carefully washed away with 100 mM NaCl buffer. The SLB on mica was properly sealed in a temperature-controlled AFM cell and cooled to 17 °C at a rate of 0.02 °C/s prior to imaging. The temperature was subsequently increased at a rate of 0.02 °C/s, and AFM images were acquired at selected temperature checkpoints during the heating ramp after allowing the membrane to stabilize, thereby crossing the membrane transition temperatures to image SLB gel-domain formation.

#### Protein Quantification

The amount of transferrin associated with liposomes was quantified using the Bradford protein assay. To release liposome-bound transferrin and ensure complete protein solubilization, purified transferrin-functionalized liposomes were disrupted using Lysis Buffer [1 mM EDTA, 150 mM NaCl, 50 mM Tris-HCl, 0.5% sodium deoxycholate, and 1% NP-40 (all from Merck, St. Louis, MO, USA), supplemented with 1× protease inhibitor cocktail (cat. 78430, ThermoFisher Scientific, Waltham, MA, USA)]. Protein lysates were quantified using the Bradford assay (cat. 23238, ThermoFisher Scientific). Samples (5 µL of sample or standard) were mixed with 250 µL of Bradford reagent in a 96-well flat-bottom plate. A standard curve was generated using Bovine Serum Albumin (BSA) (cat. A4737, Sigma-Aldrich, St. Louis, MO, USA) at concentrations of 4, 2, 1, 0.5, and 0 µg/µL, prepared in the same lysis buffer. After a 15 min incubation at room temperature in the dark, absorbance was measured at 590 nm using a Synergy H1 Hybrid Reader (BioTek, Agilent Technologies, Santa Clara, CA, USA). Liposome-associated transferrin content was calculated from the calibration curve and expressed as µg of transferrin per mg of total lipid. Empty, non-functionalized liposomes processed identically and disrupted with the same detergent concentration were used as blanks.

#### Drug Encapsulation Efficiency

Drug encapsulation efficiency was quantified by UV–Vis spectrophotometry using a Synergy H1 Hybrid Reader (BioTek, Agilent Technologies). Liposomes were disrupted in Lysis Buffer and incubated for 15 min at room temperature. Samples were centrifuged at 10,000 × *g* for 15 min to remove insoluble lipid debris. Then, 20 µL of the supernatant was transferred to a 96-well plate and diluted with 80 µL of Lysis Buffer. Niflumic acid, bumetanide, and (R)-ketorolac were quantified at their respective absorbance maxima (290 nm, 230 nm, and 315 nm). Calibration curves were prepared using the corresponding free drugs (25 to 800 µM) diluted in Lysis Buffer. Empty liposomes in Lysis Buffer were used as blanks. Liposomal drug content was calculated from the calibration curve and expressed as encapsulation efficiency (%).

### Sex as a biological variable

Sex was not considered a biological variable in this study. Clinical information was available for three patients (S82, S192 and S197), all of whom were diagnosed with grade IV glioblastoma and had received no chemotherapy before tumor resection. Their ages at diagnosis were 49, 65, and 47 years, respectively. For the remaining patients, clinical information was limited to the diagnosis of high-grade glioblastoma.

